# A Pan-Cancer Blueprint of Early Tumor Microenvironment Reprogramming

**DOI:** 10.1101/2025.11.14.688523

**Authors:** Bhavya Bhavya, Shivangi Agrawal, Ekta Pathak, Rajeev Mishra

## Abstract

Metabolic disruption is a defining early event in tumorigenesis, yet its coordination with transcriptional and inflammatory remodeling remains unclear. Across fourteen first-stage solid tumors, we integrate transcriptomic, metabolic, immune, microRNA, and network-topological analyses to construct a blueprint of early tumor microenvironment reprogramming. Gene-network evolution revealed preferential attachment of proliferative hubs (FOXM1, CDK1, CCNB1, CDC20, TOP2A) accompanied by loss of neighborhood connectivity in key metabolic regulators (PPARA, PPARG, PRKACA/B, CREB1, SRC), indicating early erosion of lipid sensing, oxidative restraint, and metabolic-neuronal signaling. Pathway analysis demonstrated consistent suppression of fatty acid degradation and PPAR signaling, establishing a shift toward anabolic growth. ADH1B was uniformly downregulated across all cancers, linking impaired aldehyde metabolism to redox stress and inflammatory activation. Integration of inflammatory and apoptotic modules highlighted recurrent IRG-AG pairs, including IL11-MMP9-FGF10, forming a conserved inflammation-apoptosis-metabolism axis.

**Graphical abstract:** 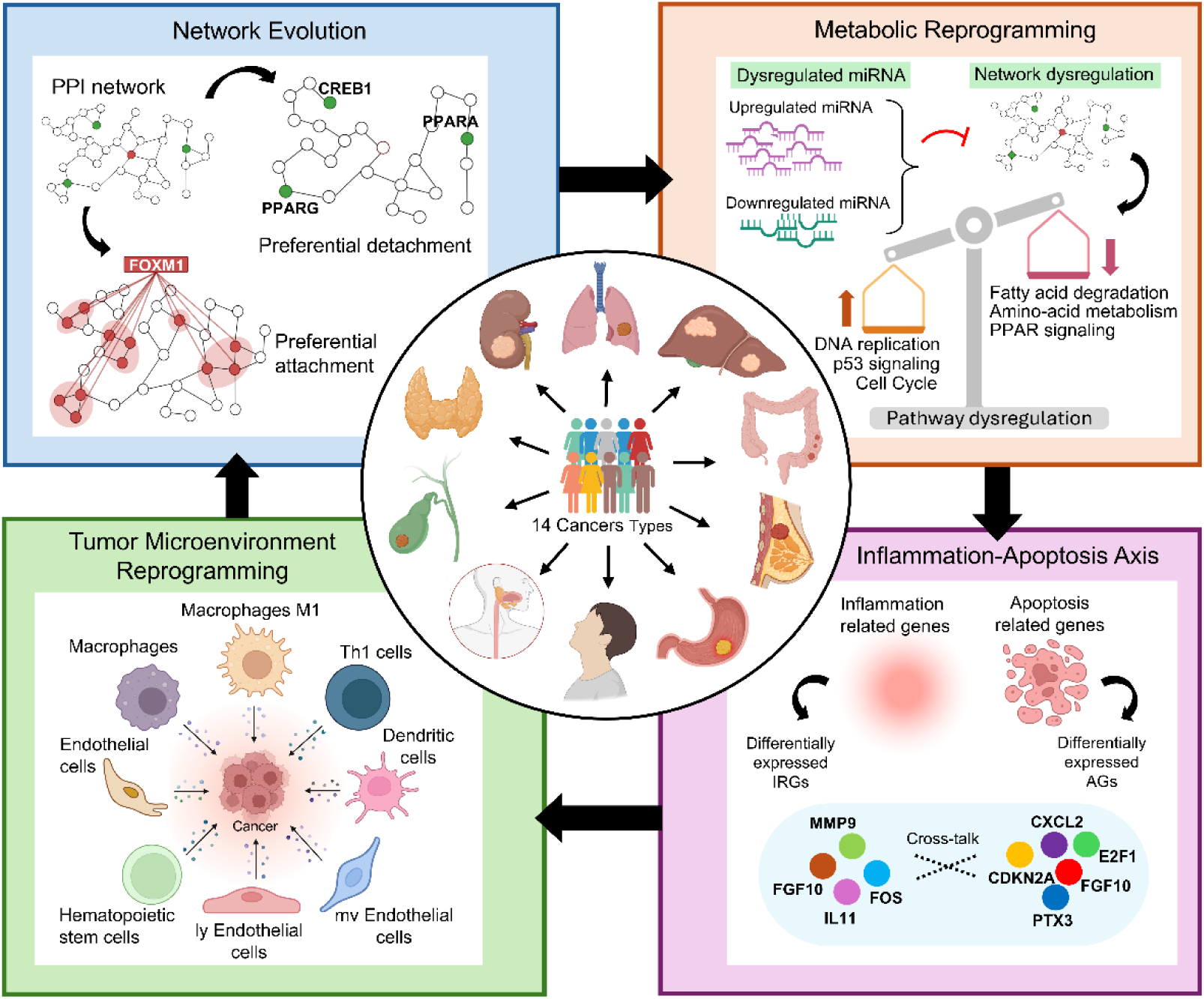

**Highlights:**

1. Network evolution reveals “trigger genes” shaping early oncogenic transformation
2. FOXM1 attachment and PPAR/CREB1 detachment coordinate metabolic reprogramming
3. Uniform ADH1B downregulation in 14 cancers links metabolic imbalance with inflammation
4. Integrated multi-omics uncovers a blueprint of early tumor-microenvironment shifts

## Introduction

Cancer remains one of the most pressing global health challenges, responsible for roughly ten million deaths annually (Bray et al., 2024). Malignant transformation is now widely understood as a systems-level transition in which transcriptional, metabolic, immune, and stromal programs reorganize in a coordinated manner, even though genomic alterations remain foundational to oncogenesis (Ahmed et al., 2023; Hanahan, 2022). Within this perspective, biological systems undergo dynamic remodeling of their interactions, where certain regulators gain network centrality through preferential attachment while others lose influence through preferential detachment, an organizing principle consistent with contemporary network-science models (Barabási and Albert, 1999; Mambetsariev et al., 2023).

The Pan-cancer efforts, notably those by The Cancer Genome Atlas (TCGA) and other large-scale consortia, have provided unprecedented opportunities to map shared oncogenic mechanisms across diverse tumor types (Hoadley et al., 2018; Weinstein et al., 2013). Yet, most analyses remain compartmentalized addressing mutations, signaling, or immunity in isolation. The tumor microenvironment (TME), a dynamic interface of tumor, stromal, and immune elements, represents the integrative nexus where these molecular events converge. The earliest stages of network remodeling, particularly the transitions shaping metabolic and immune circuits in the tumor microenvironment remain incompletely defined. Decoding the early TME at the systems scale—before extensive genomic chaos and immune escape offers a window into the origins of cancer as an emergent, adaptive network disorder. Evidence suggests that, well before the full emergence of genomic instability or advanced immune-evasion programs, nascent neoplastic ecosystems undergo substantial restructuring of cellular interactions, metabolic dependencies, and immunologic niches (De Visser and Joyce, 2023; Grant and Ferrer, 2025).

Cancer metabolism is increasingly recognized as both a driver and a consequence of network-level reprogramming (Kay and Zanivan, 2025). Metabolic reprogramming is deeply intertwined with these early alterations, shaping biosynthetic demand, redox balance, and nutrient utilization (Arner and Rathmell, 2023; Martínez-Reyes and Chandel, 2021; McGuirk et al., 2020; Peng-Winkler and Fendt, 2026). Simultaneously, metabolic cues recalibrate immunometabolic circuits that govern whether tissues become permissive or restrictive to malignant expansion (Arner and Rathmell, 2023; Mao et al., 2024). This metabolic-immune linkage is tightly embedded within the topology of gene-regulatory and signaling networks, wherein transcription factors, metabolic sensors, and cytokine mediators function as interconnected hubs guiding cell-state transitions(Kao et al., 2022; Martinez-Outschoorn et al., 2017).

Advances in multi-omics integration underscore that transcriptional, metabolic, and post-transcriptional layers operate as a coordinated system rather than isolated modalities (Chakraborty et al., 2024; He et al., 2023; Vucic et al., 2012). MicroRNAs further modulate metabolic flexibility and reinforce transcriptional programs influencing the cancer cell metabolism (Alamoudi et al., 2018), (Lee et al., 2024; Zong et al., 2023). Despite this progress, no unified cross-cancer framework yet explains how immune-stromal architecture, metabolic pathway remodeling, transcriptional hub dynamics, and miRNA regulation converge during the earliest phases of carcinogenesis.

To address this gap, we integrate transcriptomic, metabolic, immunologic, and miRNA layers across fourteen TCGA cancer types restricted to stage-I disease. We then apply a network-evolution framework to quantify preferential attachment and detachment and relate these topological shifts to metabolic and immune-microenvironmental processes. Our aim is to define shared axes of early tumor microenvironment remodeling and to outline a systems-level blueprint that may inform biomarker discovery, early detection, and metabolism-focused cancer prevention strategies.

## Results

### Network evolution from normal to first-stage cancer

To characterize transcriptional network remodeling at the onset of malignancy, we analyzed RNA-seq profiles from normal and first-stage samples across 14 cancers and identified significantly expressed genes for each condition (Table S1). PPI networks constructed from these gene sets exhibited scale-free degree distributions in all cancers, with a small number of highly connected hubs and many low-degree nodes (Figure 1A; Figure S1).

**Figure 1.**
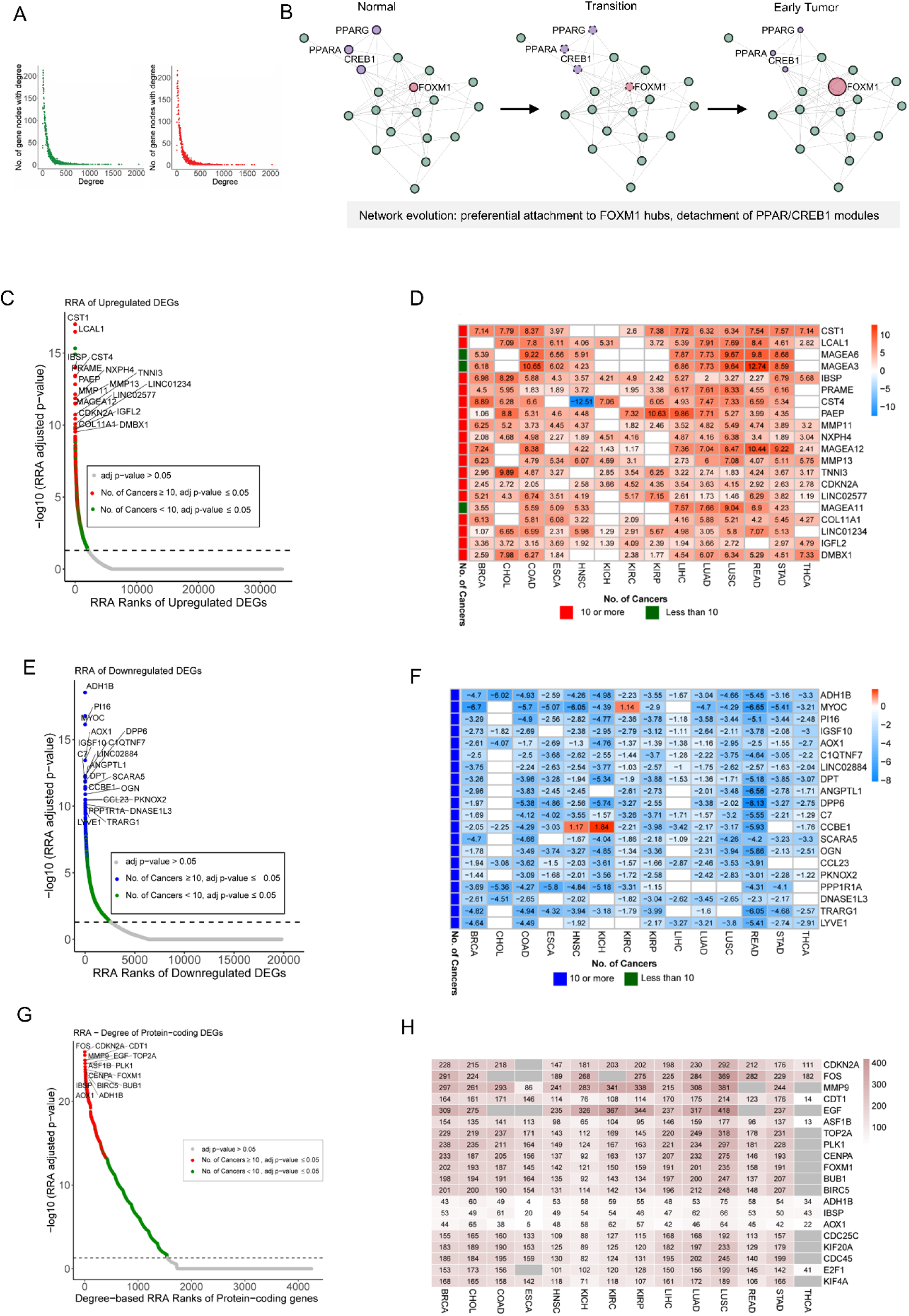
Transcriptomic and network-level RRA analyses across 14 first-stage cancers. A. Degree distributions of significantly expressed genes mapped onto their respective protein-protein interaction (PPI) networks in first-stage tumors and matched normal tissues. B. Gains and losses of interacting partners, illustrating preferential attachment and preferential detachment during network remodeling. C. RRA of upregulated genes. The dotted line marks the *padj* = 0.05 threshold. Red points denote statistically significant genes upregulated in ≥10 cancers; green points denote significant genes upregulated in <10 cancers; grey points are non-significant (*padj*>0.05). D. Heatmap of log₂FC values for the top 20 upregulated RRA genes across the 14 cancers. E. RRA of downregulated genes. The dotted line marks the *padj* = 0.05 threshold. Blue points denote significant genes downregulated in ≥10 cancers; green points denote significant genes downregulated in <10 cancers; grey points are non-significant. F. Heatmap of log₂FC values for the top 20 downregulated RRA genes across the 14 cancers. G. Degree-based RRA of protein-coding DEGs. The dotted line marks the *padj* = 0.05 threshold. Red points indicate significant DEGs present in ≥10 cancer networks; green points indicate significant DEGs present in <10 networks; grey points are non-significant. H. Heatmap of the degrees of the top 20 degree-based RRA genes across the 14 cancers.

Genes showing degree gain in ≥10 cancers were prioritized, and their mean degree change was computed. The ten most consistently expanded hubs were CDK1, CCNB1, BRCA1, BIRC5, KIF11, ASPM, CHEK1, FOXM1, TOP2A, and CDC6 (Table S2). Conversely, genes with degree loss in ≥10 cancers revealed a distinct set of top detaching nodes: SRC, AGT, DLG4, APP, PRKACA, FN1, PRKACB, NTRK2, SYP, and GNB1 (Table S3).

Pan-cancer preferential-attachment genes (top 50) formed a densely interconnected module with 11 genes—ASPM, BIRC5, BUB1, CCNA2, CCNB1, CDC20, CDK1, CENPF, KIF11, MAD2L1, TOP2A each exhibiting degree 49 and closeness centrality 1 (Figure S2A; Table S4). In contrast, the preferential-detachment network was sparsely connected, with SRC showing the highest degree (34) and closeness centrality (0.77) (Figure S2B; Table S4). Pathway enrichment revealed proliferative processes among attachment hubs and nervous system pathways among detachment genes (Table 1).

**Table 1.** Pathway Enrichment of genes with highest average degree increase/decrease.

| Pathway Enrichment of 50 genes with pan-cancer increase in degree |  |  |  |  |
| --- | --- | --- | --- | --- |
| KEGG Pathway | Count | % | p Value | Genes |
| Cell cycle | 18 | 36 | 6.24E-23 | CDCA5, KNL1, CDC6, NDC80, CCNA2, CDC20, CCNB1, CHEK1, CDK2, MCM3, CDK1, MCM4, MCM5, MCM6, FBXO5, BUB1, MCM2, MAD2L1 |
| DNA replication | 6 | 12 | 1.10E-07 | FEN1, MCM3, MCM4, MCM5, MCM6, MCM2 |
| Oocyte meiosis | 7 | 14 | 5.85E-06 | CDC20, CCNB1, CDK2, CDK1, FBXO5, BUB1, MAD2L1 |
| Progesterone-mediated oocyte maturation | 6 | 12 | 3.15E-05 | CCNA2, CCNB1, CDK2, CDK1, BUB1, MAD2L1 |
| p53 signaling pathway | 5 | 10 | 1.11E-04 | CCNB1, RRM2, CHEK1, CDK2, CDK1 |
| Cellular senescence | 6 | 12 | 1.64E-04 | CCNA2, CCNB1, CHEK1, CDK2, CDK1, FOXM1 |
| Fanconi anemia pathway | 4 | 8 | 8.22E-04 | FANCI, RAD51, BRCA1, BRCA2 |
| Viral carcinogenesis | 5 | 10 | 0.004795 | CCNA2, CDC20, CHEK1, CDK2, CDK1 |
| Human T-cell leukemia virus 1 infection | 5 | 10 | 0.006555 | CCNA2, CDC20, CHEK1, CDK2, MAD2L1 |
| Homologous recombination | 3 | 6 | 0.008423 | RAD51, BRCA1, BRCA2 |
| Pathway Enrichment of 50 genes with pan-cancer decrease in degree |  |  |  |  |
| Dopaminergic synapse | 20 | 40 | 1.05E-23 | GSK3B, CACNA1D, ARRB1, CACNA1C, ARRB2, COMT, GNG12, CREB1, GNG2, GNG7, GNAQ, GNB2, GNB1, GNB4, GNAS, GNB5, PLCB1, PRKACA, PLCB2, PRKACB |
| Serotonergic synapse | 18 | 36 | 2.34E-21 | APP, CACNA1D, CACNA1C, GNG12, GNG2, GNG7, GNAQ, GNB2, GNB1, GNB4, GNAS, KRAS, GNB5, PLCB1, PRKACA, HRAS, PLCB2, PRKACB |
| Relaxin signaling pathway | 18 | 36 | 2.09E-20 | SRC, ARRB1, ARRB2, GNG12, CREB1, GNG2, GNG7, GNB2, GNB1, GNB4, GNAS, KRAS, GNB5, PLCB1, PRKACA, HRAS, PLCB2, PRKACB |
| Cholinergic synapse | 17 | 34 | 1.39E-19 | CACNA1D, CACNA1C, GNG12, CREB1, GNG2, GNG7, GNAQ, GNB2, GNB1, GNB4, KRAS, GNB5, PLCB1, PRKACA, HRAS, PLCB2, PRKACB |
| Glutamatergic synapse | 17 | 34 | 1.39E-19 | CACNA1D, CACNA1C, GNG12, GNG2, GRK2, DLG4, GNG7, GNAQ, GNB2, GNB1, GNB4, GNAS, GNB5, PLCB1, PRKACA, PLCB2, PRKACB |
| Circadian entrainment | 16 | 32 | 3.83E-19 | CACNA1D, CACNA1C, GNG12, CREB1, GNG2, GNG7, GNAQ, GNB2, GNB1, GNB4, GNAS, GNB5, PLCB1, PRKACA, PLCB2, PRKACB |
| Chemokine signaling pathway | 19 | 38 | 6.45E-19 | GSK3B, SRC, ARRB1, ARRB2, GNG12, GNG2, GRK2, GNG7, GNAQ, GNB2, GNB1, GNB4, KRAS, GNB5, PLCB1, PRKACA, HRAS, PLCB2, PRKACB |
| Human cytomegalovirus infection | 18 | 36 | 2.98E-16 | GSK3B, SRC, GNG12, CREB1, GNG2, GNG7, GNAQ, GNB2, GNB1, GNB4, GNAS, KRAS, GNB5, PLCB1, PRKACA, HRAS, PLCB2, PRKACB |
| Apelin signaling pathway | 15 | 30 | 3.99E-15 | GNG12, GNG2, CDH1, GNG7, GNAQ, GNB2, GNB1, GNB4, KRAS, GNB5, PLCB1, PRKACA, HRAS, PLCB2, PRKACB |
| Morphine addiction | 13 | 26 | 1.63E-14 | ARRB1, ARRB2, GNG12, GNG2, GRK2, GNG7, GNB2, GNB1, GNB4, GNAS, GNB5, PRKACA, PRKACB |

### Pan-cancer differentially expressed and influential genes

Differential expression analysis across the 14 cancers identified 451 genes upregulated in ≥10 cancers and 159 genes downregulated in ≥10 cancers (Table S5; Figure S3). The top RRA-ranked upregulated genes were CST1, LCAL1, IBSP, PRAME, CST4, PAEP, MMP11, NXPH4, MAGEA12, and MMP13 (Figure 1C-D). The top RRA-ranked downregulated genes were ADH1B, MYOC, PI16, IGSF10, AOX1, C1QTNF7, LINC02884, DPT, ANGPTL1, and DPP6 (Figure 1E-F).

Gene-gene interaction (GGI) networks were constructed for protein-coding DEGs in each tumor type (Table S6). RRA integration of network degree and MCC centrality identified CDKN2A, FOS, MMP9, CDT1, and EGF as the top influential genes recurrently dysregulated in >10 cancers (Figure 1G-H).

### Pan-cancer influential DEGs involved in network evolution

Integration of DEGs with network-evolution modules revealed 28 pan-cancer upregulated genes with preferential-attachment properties, all of which were also topologically influential. FOXM1 occupied a central position, interacting with 46 genes, including 27 pan-cancer upregulated and influential nodes (Figure 2A). No preferential-detachment genes were both pan-cancer dysregulated and influential (Figure 2B).

**Figure 2.**
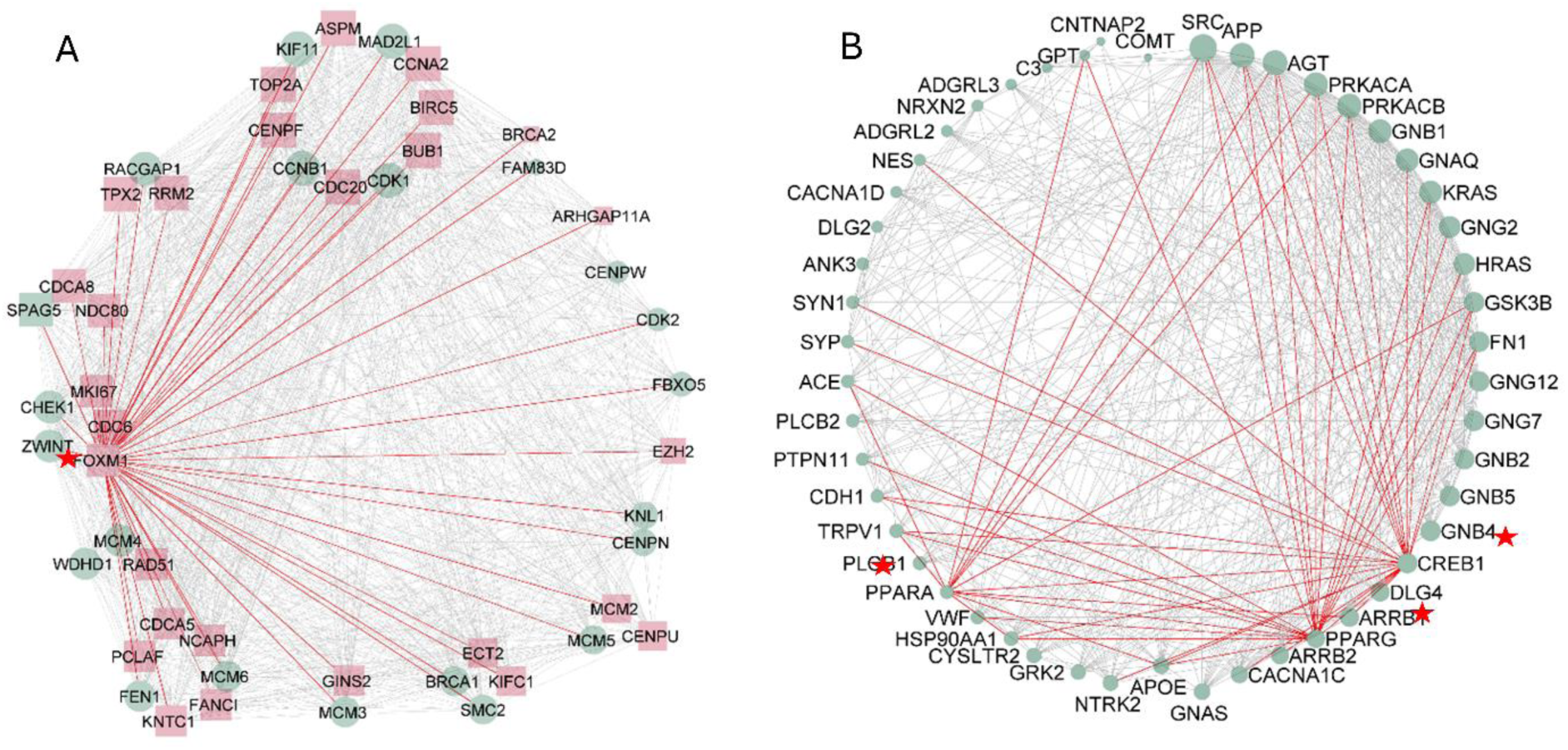
Pan-cancer influential genes contributing to network evolution. A. PPI network of the 50 genes exhibiting the highest average degree increase in first-stage cancers. B. PPI network of the 50 genes with the highest average degree decrease. Red and blue nodes denote pan-cancer upregulated and downregulated genes, respectively; square-shaped nodes highlight influential genes. Node size decreases with decreasing degree across cancers.

### Pan-cancer dysregulated metabolic pathways

KEGG pathway analysis revealed consistent metabolic and biosynthetic reprogramming across tumor types (Figure S4A-N). The most strongly upregulated pathways included Ribosome biogenesis in eukaryotes (COAD, LIHC, LUSC, READ, STAD) and DNA replication (HNSC, LUAD). Additional top pathways varied across cancers, including proteasome (BRCA), cell cycle (CHOL), spliceosome (ESCA), oxidative phosphorylation (KICH), graft-versus-host disease (KIRC), ribosome (KIRP), and bacterial invasion of epithelial cells (THCA).

Downregulated pathways were dominated by metabolic and stromal programs, including valine, leucine, and isoleucine degradation (ESCA, KIRC, THCA), hematopoietic cell lineage (LIHC, LUSC), and oxidative phosphorylation (HNSC, STAD). Additional cancer-specific downregulated pathways included regulation of lipolysis, complement and coagulation, bile secretion, fatty acid degradation, platelet activation, and calcium signaling (Figure S4A-N).

RRA of 300 pathways identified 22 significantly downregulated pathways (FDR < 0.05), of which 13 were downregulated in ≥10 cancers (Figure 3A-D). These included fatty acid degradation, valine/leucine/isoleucine degradation, PPAR signaling, propanoate metabolism, drug metabolism (cytochrome P450), butanoate metabolism, regulation of lipolysis, vascular smooth muscle contraction, calcium signaling, mineral absorption, adipocytokine signaling, fatty acid metabolism, and bile secretion. Across all cancers, we identified 10 pan-cancer upregulated pathways driven by upregulated genes and 13 pan-cancer downregulated pathways driven by downregulated genes (Table 2).

**Figure 3.**
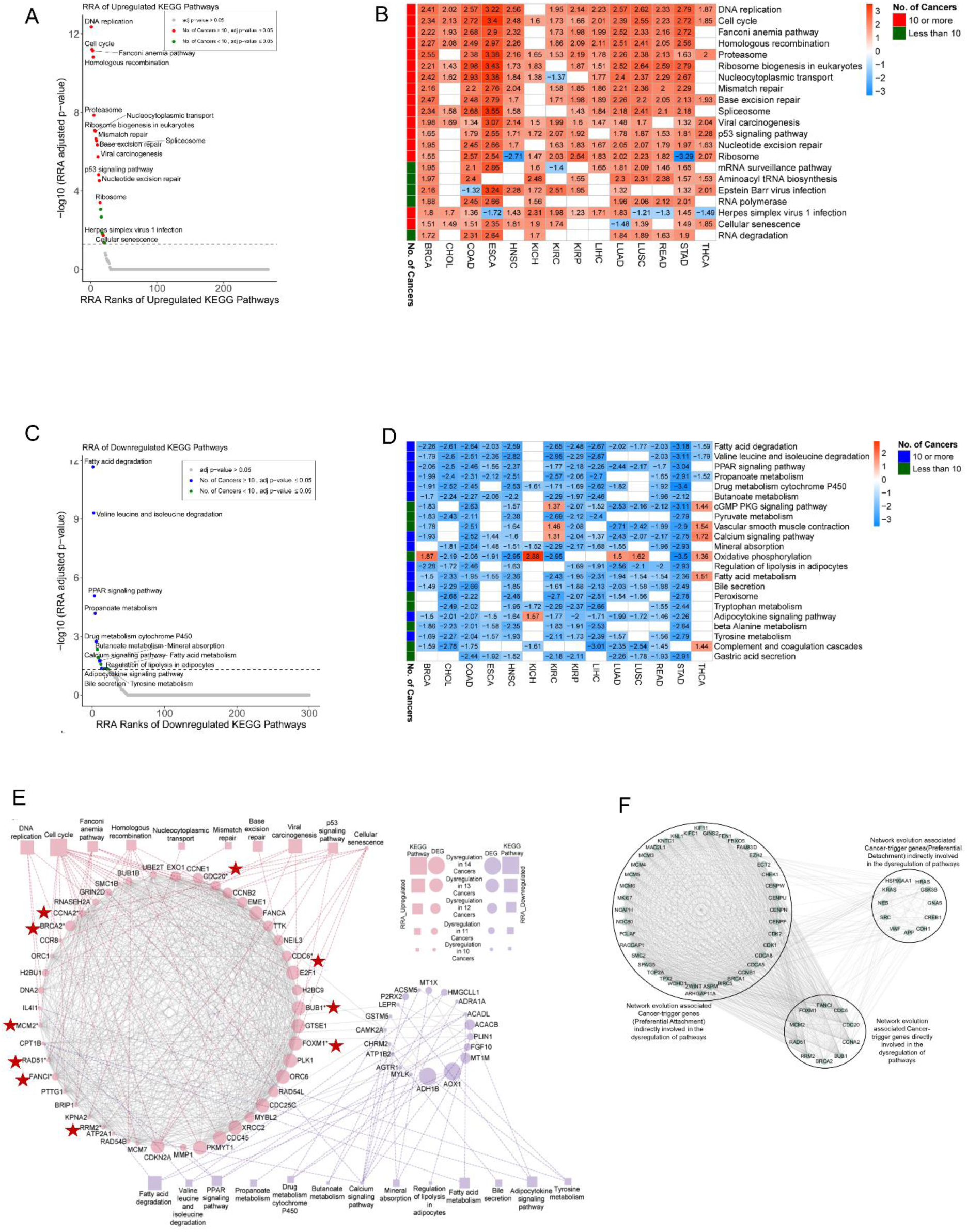
Pan-cancer pathway-level RRA analyses and gene-pathway integration. A. RRA of upregulated KEGG pathways. The dotted line marks *padj* = 0.05. Red points denote significant pathways upregulated in ≥10 cancers; green points denote significant pathways upregulated in <10 cancers; grey points are non-significant. B. Heatmap of NES values for significantly upregulated pathways. C. RRA of downregulated KEGG pathways, using the same color conventions (blue for ≥10 cancers). D. Heatmap of NES values for significantly downregulated pathways. E. PPI network linking pan-cancer dysregulated genes with dysregulated pathways. Upregulated and downregulated genes appear as red and blue circles; pathways appear as red and blue squares. Grey edges depict PPIs; dotted lines indicate gene-pathway associations. Starred genes denote those implicated in network evolution. F. Indirect associations between cancer-trigger genes and pathway dysregulation.

**Table 2.** Pan-cancer dysregulated genes that are associated with pan-cancer dysregulated pathways.

| Pan-cancer Dysregulated Pathway | Dysregulation of Pathway | No. of pan-cancer dysregulated genes associated with the pathway | Pan-cancer dysregulated genes associated with the pathway |
| --- | --- | --- | --- |
| Cell cycle | Upregulated | 20 | BUB1*, BUB1B, CCNA2*, CCNB2, CCNE1, CDC20*, CDC25C, CDC45, CDC6*, CDKN2A, E2F1, MCM2*, MCM7, ORC1, ORC6, PKMYT1, PLK1, PTTG1, SMC1B, TTK |
| Cellular senescence | Upregulated | 7 | CCNA2*, CCNB2, CCNE1, CDKN2A, E2F1, FOXM1*, MYBL2 |
| Fanconi anemia pathway | Upregulated | 7 | BRCA2*, BRIP1, EME1, FANCA, FANCI*, RAD51*, UBE2T |
| Homologous recombination | Upregulated | 7 | BRCA2*, BRIP1, EME1, RAD51*, RAD54B, RAD54L, XRCC2 |
| p53 signaling pathway | Upregulated | 5 | CCNB2, CCNE1, CDKN2A, GTSE1, RRM2* |
| DNA replication | Upregulated | 4 | DNA2, MCM2*, MCM7, RNASEH2A |
| Viral carcinogenesis | Upregulated | 4 | CCR8, CDKN2A, H2BC9, H2BU1 |
| Base excision repair | Upregulated | 1 | NEIL3 |
| Mismatch repair | Upregulated | 1 | EXO1 |
| Nucleocytoplasmic transport | Upregulated | 1 | KPNA2 |
| Calcium signaling pathway | Downregulated | 9 | ADRA1A, AGTR1, ATP2A1, CAMK2A, CHRM2, FGF10, GRIN2D, MYLK, P2RX2 |
| PPAR signaling pathway | Downregulated | 4 | ACADL, CPT1B, MMP1, PLIN1 |
| Adipocytokine signaling pathway | Downregulated | 3 | ACACB, CPT1B, LEPR |
| Drug metabolism cytochrome P450 | Downregulated | 3 | ADH1B, AOX1, GSTM5 |
| Fatty acid degradation | Downregulated | 3 | ACADL, ADH1B, CPT1B |
| Mineral absorption | Downregulated | 3 | ATP1B2, MT1M, MT1X |
| Tyrosine metabolism | Downregulated | 3 | ADH1B, AOX1, IL4I1 |
| Valine leucine and isoleucine degradation | Downregulated | 3 | AOX1, HMGCLL1, IL4I1 |
| Butanoate metabolism | Downregulated | 2 | ACSM5, HMGCLL1 |
| Fatty acid metabolism | Downregulated | 2 | ACADL, CPT1B |
| Bile secretion | Downregulated | 1 | ATP1B2 |
| Propanoate metabolism | Downregulated | 1 | ACACB |
| Regulation of lipolysis in adipocytes | Downregulated | 1 | PLIN1 |

\*Cancer trigger genes associated with evolution of gene interaction network from normal to first-stage cancer condition

Mapping network-evolution genes to dysregulated pathways identified 10 preferential-attachment cancer-trigger genes FOXM1, BUB1, CDC6, CDC20, CCNA2, BRCA2, MCM2, RAD51, FANCI, and RRM2 linked to pathways including the cell cycle, fanconi anemia, homologous recombination, p53 signaling, and DNA replication. In total, 42 upregulated and 19 downregulated genes were matched to pan-cancer dysregulated pathways, with an additional 51 evolution-linked interactors including 40 attachment and 11 detachment genes (Figure 3C-D). Notably, CREB1, a key preferential-detachment regulator, interacted with CCNA2, linking network evolution to dysregulated cell cycle and cellular senescence pathways.

### Pan-cancer differentially expressed IRGs and AGs

To define conserved immune-apoptotic interactions emerging at the onset of malignancy, we integrated 704 non-redundant immune-related genes (IRGs) and 1305 apoptosis-related genes (AGs) with the DEGs identified in each first-stage cancer. This yielded statistically significant upregulated and downregulated deIRGs and deAGs for all 14 tumor types, from which we reconstructed cancer-specific deIRG-deAG PPI networks (Figure S5-S18).

Across these networks, a strikingly recurrent set of six IRG-AG interaction pairs emerged in 11 cancers: CXCL2-MMP9, FOS-CDKN2A, FOS-E2F1, IL11-FGF10, IL11-MMP9, and PTX3-FGF10 (Figure 4A). These axes were absent only in specific tumor subsets—CXCL2-MMP9 in ESCA, READ, THCA; FOS-CDKN2A and FOS-E2F1 in COAD, ESCA, KIRC; IL11-FGF10 and PTX3-FGF10 in BRCA, CHOL, LIHC; and IL11-MMP9 in LIHC, READ, THCA.

**Figure 4.**
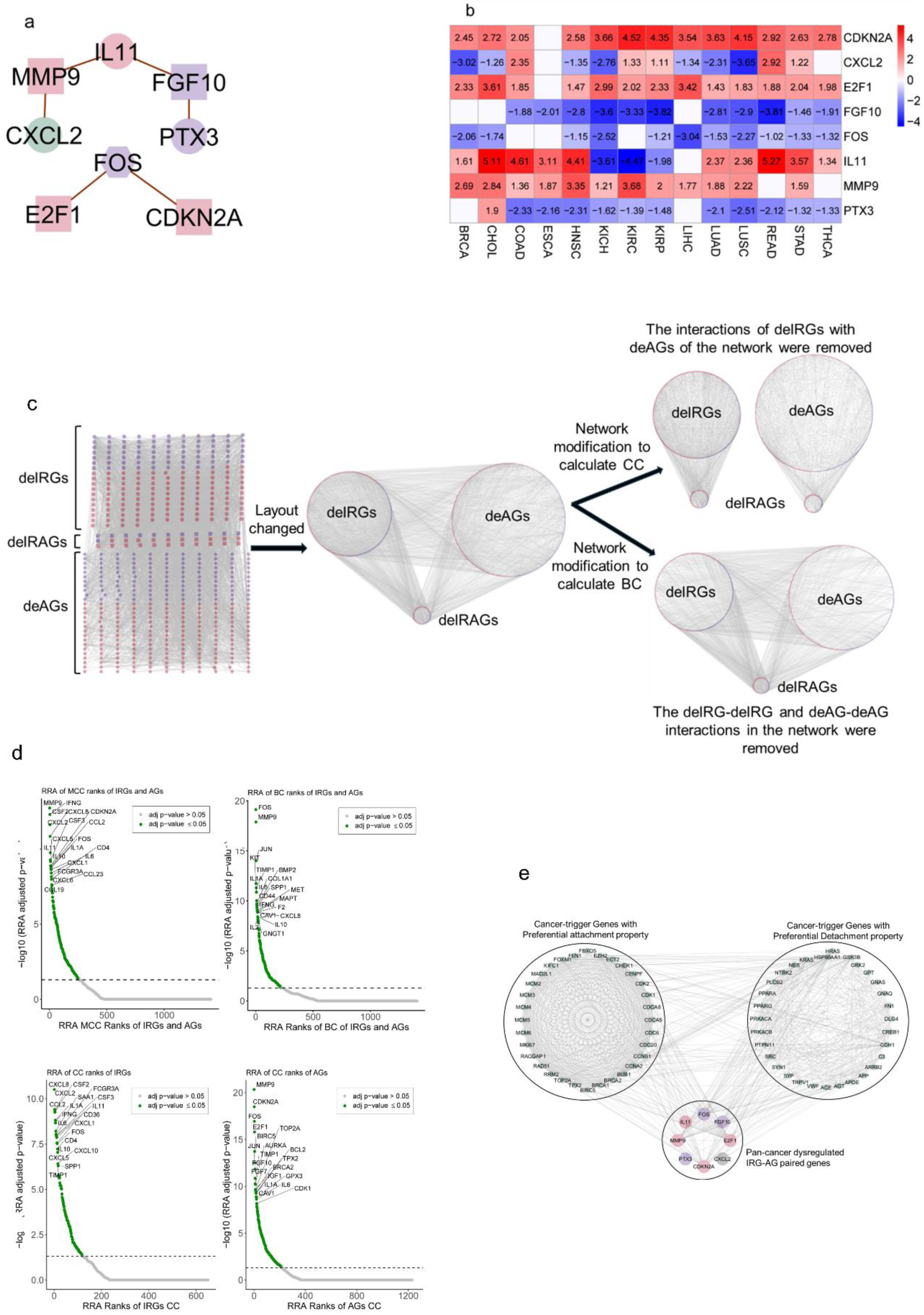
Dysregulated IRG-AGinteraction architecture and its association with network evolution. A. Interaction network of IRG-AGgene pairs recurrently dysregulated across cancers. Node shapes represent IRGs (circles), AGs (squares), and IRAGs (hexagons). Red and blue colors denote pan-cancer upregulated and downregulated genes; green marks pan-cancer DEGs. B. Heatmap of log₂FC values of IRG-AGgene pairs. C. Alterations in IRG-AGinteraction networks across 14 cancers. D. RRA of network parameters (MCC, BC, CC ranks for IRGs; CC ranks for AGs). Dotted line marks *padj* = 0.05; green points denote significant genes. E. Association of core dysregulated IRG-AGpairs with network evolution. Green nodes represent cancer-trigger genes; red and blue nodes represent pan-cancer upregulated and downregulated IRGs/AGs; the grey node denotes a pan-cancer DE IRG.

The genes forming these pairs displayed consistent pan-cancer dysregulation. CDKN2A, E2F1, and MMP9 were upregulated in 13, 13, and 12 cancers, respectively. FGF10, FOS, and PTX3 were downregulated in 11 cancers, with PTX3 uniquely upregulated in CHOL. IL11 was upregulated in 10 cancers but downregulated in KICH, KIRC, and KIRP, whereas CXCL2 showed a bidirectional pattern, upregulated in 5 and downregulated in 7 cancers (Figure 4B). Together, these findings reveal a conserved inflammatory-cell-cycle-stromal axis recurrently rewired across early tumors.

### Pan-cancer influential deIRGs and deAGs

To determine whether these dysregulated nodes exert broad network-level influence, we quantified centrality metrics for all deIRGs and deAGs in each cancer network. MCC ranks were computed from complete networks, and betweenness and closeness centrality (BC, CC) were derived from modified network topologies (Figure 4B).

RRA integration of MCC ranks across cancers identified 20 influential genes, including MMP9, CSF2, IFNG, CXCL2, CXCL8, IL11, CXCL5, CSF3, IL10, CDKN2A, CCL2, FOS, IL1A, CXCL1, IL6, FCGR3A, CD4, CXCL6, CCL19, and CCL23. Parallel RRA of BC ranks highlighted FOS, MMP9, JUN, IL1A, TIMP1, KIT, IL6, CD44, COL1A1, BMP2, SPP1, IFNG, MET, MAPT, F2, CAV1, CXCL8, IL10, GNGT1, and IL2 as key information-flow mediators.

CC-based RRA further identified top influential IRGs (CXCL8, CCL2, CXCL2, CSF2, IL1A, IFNG, IL6, SAA1, FCGR3A, CSF3, IL11, CD36, CXCL1, FOS, CD4, IL10, CXCL5, CXCL10, SPP1, TIMP1) and influential AGs (MMP9, CDKN2A, FOS, E2F1, BIRC5, FGF10, JUN, FGF7, TIMP1, AURKA, TOP2A, IGF1, BRCA2, TPX2, BCL2, IL1A, CAV1, GPX3, IL6, CDK1).

Notably, all eight genes forming the core IRG-AG pairs were consistently recovered as influential across multiple centrality frameworks (Figure 4C), positioning them as early pan-cancer immune-apoptotic regulators.

### Association of core dysregulated IRG-AGpairs with network evolution

We next evaluated whether the conserved IRG-AGaxis intersects with the cancer-trigger genes underlying network evolution from normal to first-stage cancer. The eight dysregulated IRG-AGgenes interacted extensively with 32 preferential-attachment and 31 preferential-detachment genes, including central master regulators FOXM1, PPARA, PPARG, and CREB1 (Figure 4D). This places the immune-apoptotic dysregulation directly within the architecture of early network remodeling.

### Multi-cancer differentially expressed miRNAs with dysregulated target gene sets

To assess whether post-transcriptional programs reinforce these early perturbations, we profiled differentially expressed miRNAs in each cancer (Table S8; Figure S19). RRA ranked 943 upregulated and 681 downregulated miRNAs across cancers (Figure S20). We then defined miRNAs whose expression correlated with dysregulation of their target gene sets (TGS) in each cancer (Figure S21A-N). RRA of miRNA_upTGS and miRNA_dnTGS identified 17 and 65 statistically significant miRNAs, respectively (Figure S22, S23).

Ten miRNAs were significantly RRA-ranked both for upregulation and for downregulated target gene sets: hsa-miR-106b-5p, hsa-miR-130b-3p, hsa-miR-141-3p, hsa-miR-182-5p, hsa-miR-200a-3p, hsa-miR-301a-3p, hsa-miR-429, hsa-miR-93-5p, hsa-miR-9-5p, and hsa-miR-96-5p (Figure 5A-B). Although no cancer showed this pattern for all 10 miRNAs, each miRNA displayed the up-miRNA/down-TGS signature in at least one cancer. Their occurrence spanned BRCA, COAD, LIHC, LUAD, LUSC, READ, STAD, KIRC, and KIRP (Figure 5C-D). This set represents a conserved module of multi-cancer upregulated miRNAs exerting broad repressive pressure on gene targets during early tumorigenesis.

**Figure 5.**
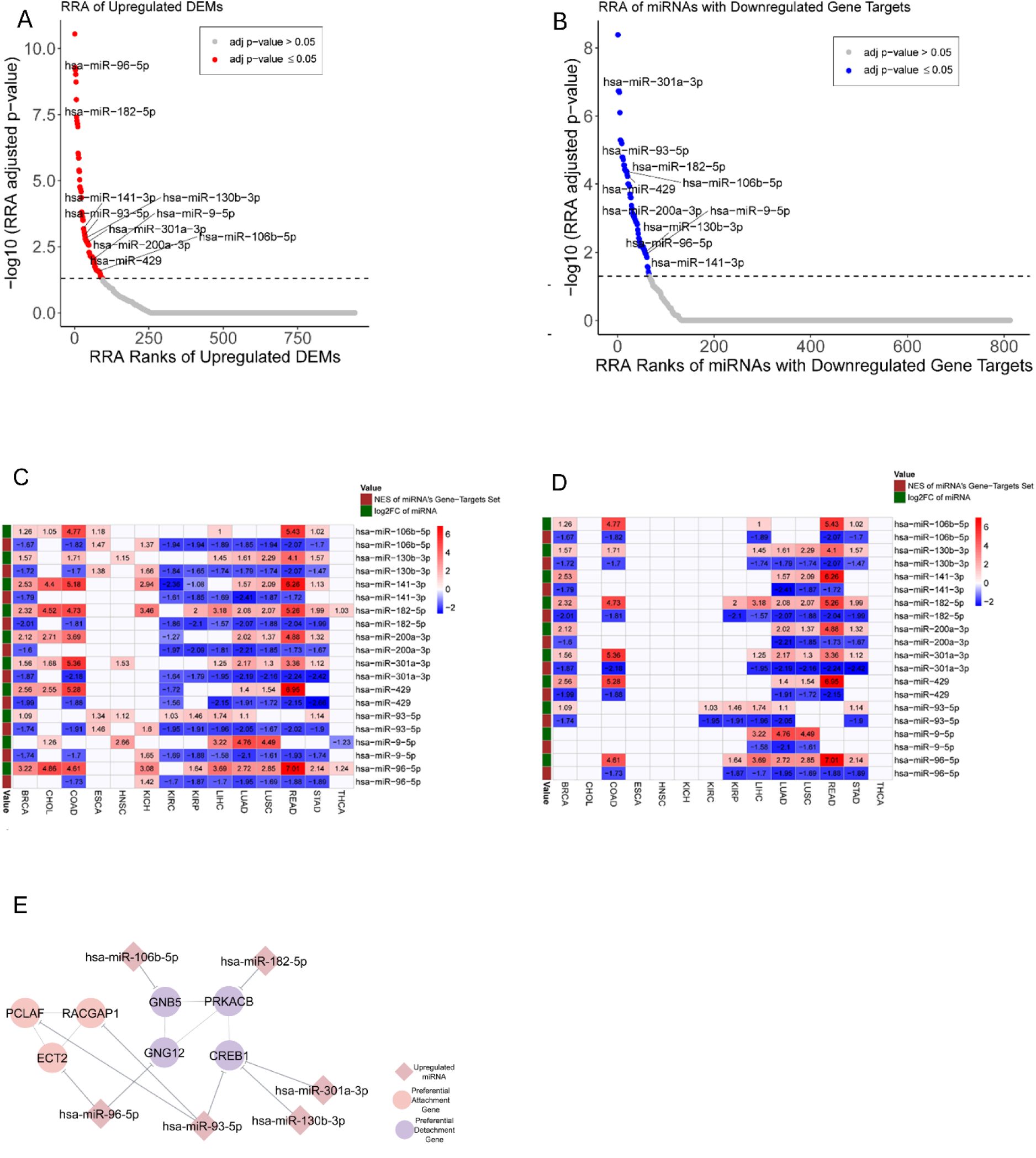
Pan-cancer miRNA deregulation and target suppression associated with cancer-trigger genes. A. RRA of upregulated miRNAs. The dotted line marks *padj* = 0.05; red points denote significant miRNAs. B. RRA of miRNAs whose target gene sets are downregulated. Blue points denote significant miRNAs; grey points are non-significant. C. Heatmap showing log₂FC and NES values for the 10 miRNAs significant in both upregulation and target-downregulation analyses. D. Heatmap showing the log₂FC and NES values for miRNAs upregulated in each cancer alongside their downregulated target sets. E. Cancer-trigger genes mapped as targets of these miRNAs.

### Association of miRNA dysregulation with network evolution

Integration of miRNA targets with the network evolution gene set revealed six of the ten multi-cancer upregulated miRNAs targeting seven network-evolution genes (Figure 5E). Among preferential-attachment genes, PCLAF and RACGAP1 were targets of hsa-miR-93-5p, while ECT2 was targeted by hsa-miR-96-5p. Among preferential-detachment genes, GNB5 was targeted by hsa-miR-106b-5p, PRKACB by hsa-miR-182-5p, and CREB1 by hsa-miR-301a-3p, hsa-miR-130b-3p, and hsa-miR-93-5p. These findings link early miRNA rewiring directly to topological drivers of network evolution, creating a multi-layered regulatory interface between miRNA dysregulation and early cancer-trigger genes.

### Differential infiltration of cells in cancers

To determine whether early tumors exhibit conserved immune-stromal reorganization, we compared infiltration scores of 64 cell types between normal and first-stage cancers across all 14 tumor types. This analysis revealed a reproducible pattern of increased infiltration of Mesenchymal stem cells, T helper 1 (Th1) cells, T helper 2 (Th2) cells, and Common lymphoid progenitors, indicating coordinated recruitment of pro-inflammatory and stromal remodeling compartments during early tumorigenesis. Conversely, Hematopoietic stem cells (HSCs), Adipocytes, Fibroblasts, and Chondrocytes consistently showed reduced infiltration, reflecting early depletion of homeostatic stromal and progenitor populations in first-stage cancers (Figure 6A).

**Figure 6.**
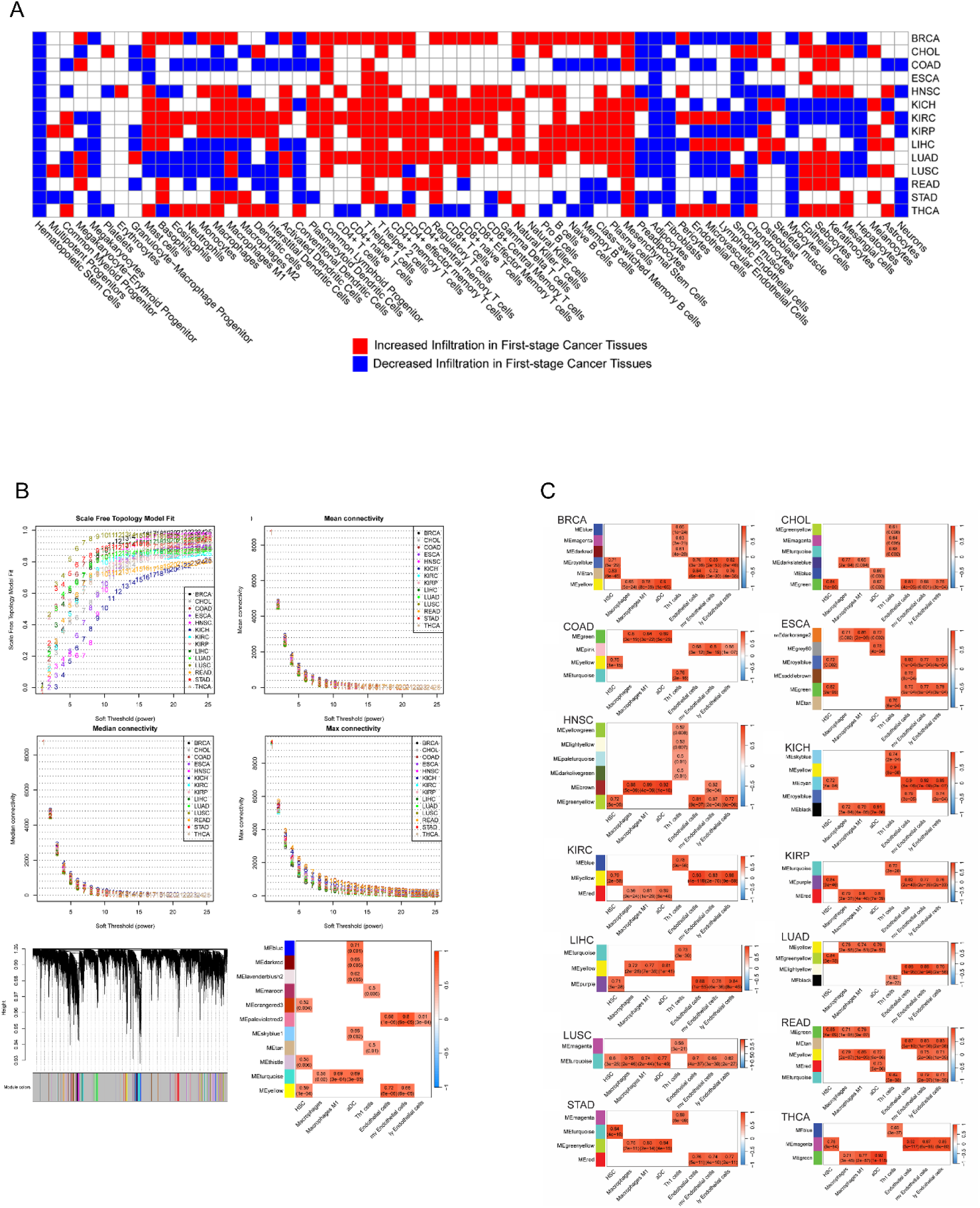
Cell-infiltration landscape and consensus/individual WGCNA across 14 cancers. A. Differential infiltration of 64 cell types between first-stage tumors and matched normal tissues. B. Pan-cancer Consensus WGCNA. Soft-threshold power (β) selection maintained a scale-free topology fit index ≥0.8. Plots show mean, median, and mode connectivity across β values. Gene dendrogram shows consensus module structure; heatmap shows correlations (Pearson coefficient and *p*-value) between module eigengenes and PCI cell types. C. Individual WGCNA for each cancer, showing high positive correlations between PCI-associated module eigengenes and PCI cells.

### Consensus Weighted Correlation Network Analysis (WGCNA) and validation

To identify gene modules underlying these infiltration changes, we performed consensus WGCNA across 17,333 genes after low-count filtering of the pan-cancer RNA-seq dataset. This analysis yielded 125 consensus modules of positively co-expressed genes. Module-trait correlations revealed 11 gene modules significantly and positively associated with infiltration scores of eight cell types, which we defined as pan-cancer infiltration-associated (PCI) cells: HSCs, activated dendritic cells (aDC), macrophages, macrophages M1, Th1 cells, endothelial cells, microvascular endothelial (mv endothelial) cells, and lymphatic endothelial (ly endothelial) cells (Figure 6B).

To confirm these associations within individual cancers, we performed separate WGCNA analyses for each tumor type (Figure S24). Because consensus WGCNA identified only positive correlations between modules and PCI cells, validation was restricted to modules showing strong positive correlations in each cancer (Figure 6C). These findings establish a reproducible cross-cancer link between PCI cell infiltration and coordinated transcriptional programs underpinning early microenvironmental remodeling.

### Pan-cancer network model

We next quantified the gene composition of the PCI-associated modules (csPCI-DEGs) (Figure 7A) and identified a subset of PCI-associated genes that were consistently differentially expressed across most cancers and present across all 14 tumor types (Figure 7B). From this subset, we selected core genes associated with specific PCI cell types.

**Figure 7.**
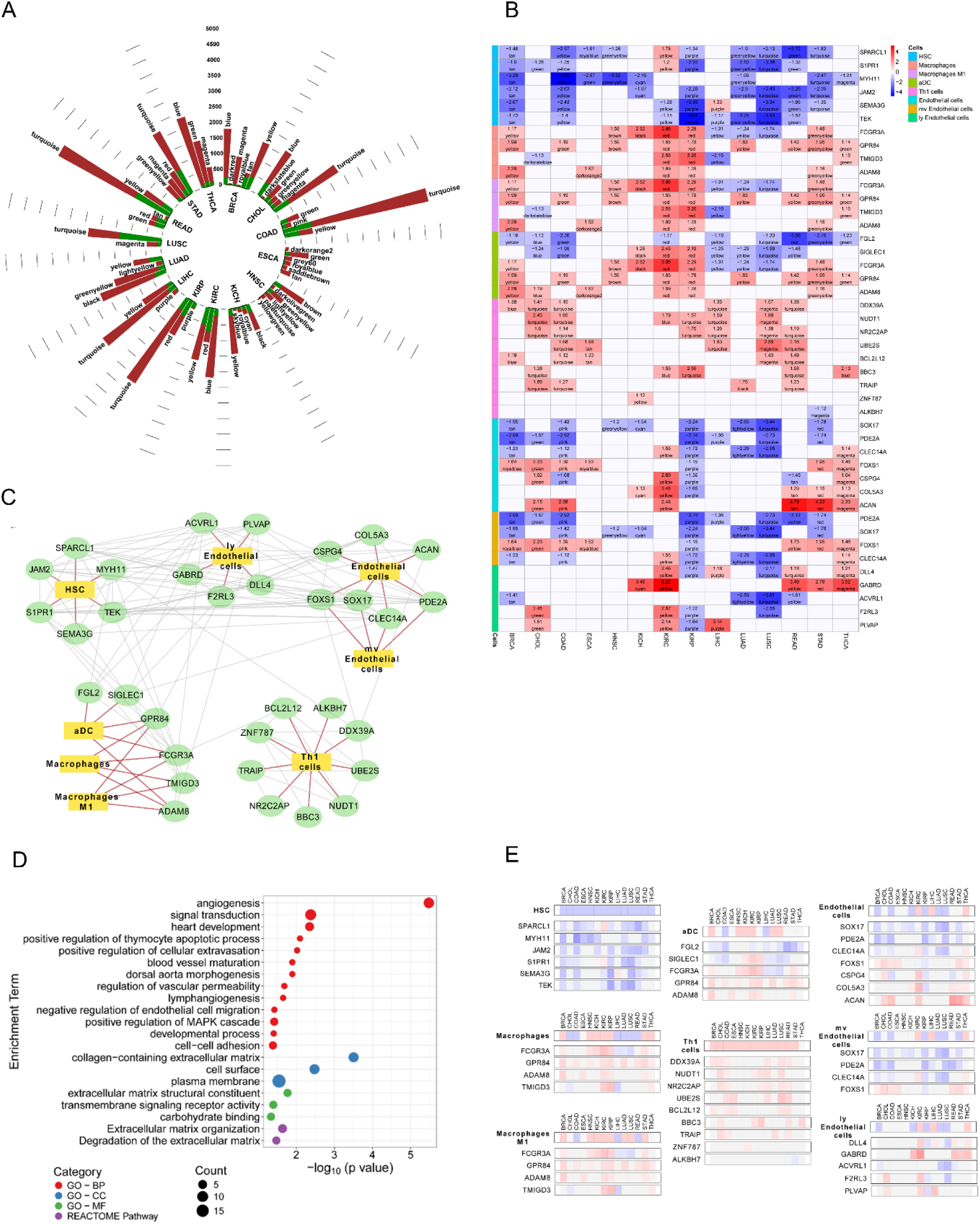
PCI-associated DEGs and the construction of the pan-cancer PCI network model. **A.** Number of genes and DEGs within PCI-associated modules across the 14 cancers; brown and green bars represent total genes and DEGs, respectively. **B.** Heatmap of log₂FC values for selected csPCI-DEGs; module membership is annotated in each cell. **C.** Pan-cancer network model of interactions among PCI-associated genes. **D.** Functional and pathway enrichment of genes used in constructing the PCI network model. **E.** Heatmaps of PCI-cell infiltration changes and log₂FC of associated csPCI-DEGs across cancers. Red/blue colors represent increased/decreased infiltration and gene upregulation/downregulation.

For HSC-associated infiltration, we identified SPARCL1, TEK, JAM2, SEMA3G, S1PR1, and MYH11. For Macrophage and Macrophage M1 infiltration, the selected genes were FCGR3A, GPR84, TMIGD3, and ADAM8. For aDCs, the associated genes were FGL2, SIGLEC1, FCGR3A, GPR84, and ADAM8. For Th1 cells, we identified DDX39A, NUDT1, NR2C2AP, UBE2S, BCL2L12, BBC3, TRAIP, ZNF787, and ALKBH7. For Endothelial cells, the set included SOX17, PDE2A, CLEC14A, FOXS1, CSPG4, COL5A3, and ACAN.

For mv endothelial infiltration, the overlapping genes were PDE2A, SOX17, FOXS1, and CLEC14A. For ly endothelial cells, we identified DLL4, AC026369.2, GABRD, ACVRL1, F2RL3, and PLVAP.

To integrate these findings into a unified framework of early microenvironmental communication, we constructed a GGI network using all selected PCI-associated genes. This network captured the pan-cancer interaction landscape through which PCI cells are interconnected via dysregulated gene programs (Figure 7C).

Network topology highlighted MYH11, S1PR1, SPARCL1, and TEK as the nodes with the highest degree, indicating central roles in early intercellular crosstalk. MYH11, FCGR3A, and TEK exhibited the highest closeness centrality, consistent with efficient signal propagation across the network. FCGR3A, MYH11, and DDX39A showed the highest betweenness centrality, identifying them as critical bridges linking sub-modules within the PCI network (Table 3).

**Table 3.** Network Topological Parameters of genes associated with TME-infiltrating cells.

| **Genes** | **Degree** | **Closeness Centrality** | **Betweenness Centrality** |
| --- | --- | --- | --- |
| MYH11 | **12** | **0.553571429** | **0.189150664** |
| S1PR1 | **12** | 0.508196721 | 0.08793177 |
| SPARCL1 | **12** | 0.5 | 0.076101181 |
| TEK | **12** | **0.516666667** | 0.087453252 |
| ACVRL1 | 10 | 0.492063492 | 0.033214062 |
| JAM2 | 10 | 0.484375 | 0.01675829 |
| CLEC14A | 9 | 0.430555556 | 0.013575773 |
| FCGR3A | 9 | **0.525423729** | **0.209453211** |
| PDE2A | 9 | 0.476923077 | 0.028810861 |
| PLVAP | 9 | 0.46969697 | 0.042890425 |
| COL5A3 | 6 | 0.418918919 | 0.019043352 |
| CSPG4 | 6 | 0.442857143 | 0.026379797 |
| DLL4 | 6 | 0.413333333 | 0.010295809 |
| F2RL3 | 6 | 0.402597403 | 0.032345134 |
| SIGLEC1 | 6 | 0.449275362 | 0.027311231 |
| SOX17 | 6 | 0.407894737 | 0.00119133 |
| ACAN | 5 | 0.455882353 | 0.036829634 |
| DDX39A | 5 | 0.449275362 | **0.182776847** |
| SEMA3G | 5 | 0.436619718 | 0.007339107 |
| UBE2S | 5 | 0.373493976 | 0.104290835 |
| ZNF787 | 5 | 0.369047619 | 0.07515617 |
| BCL2L12 | 4 | 0.407894737 | 0.132043942 |
| FGL2 | 4 | 0.424657534 | 0.001792115 |
| NUDT1 | 4 | 0.326315789 | 0.026779314 |
| TMIGD3 | 4 | 0.418918919 | 0.012398175 |
| FOXS1 | 3 | 0.424657534 | 0.015497758 |
| GPR84 | 3 | 0.369047619 | 0.004050179 |
| TRAIP | 3 | 0.281818182 | 0.064516129 |
| ADAM8 | 2 | 0.3875 | 0.001433692 |
| GABRD | 2 | 0.356321839 | 9.32E-04 |
| BBC3 | 1 | 0.271929825 | 0 |
| NR2C2AP | 1 | 0.221428571 | 0 |
| ALKBH7 | 0 | 0 | 0 |

### Pan-cancer blueprint of early TME reprogramming

To integrate the multi-layered alterations observed across cancers, we mapped all pan-cancer results onto the PCI-based interaction network, generating a unified blueprint of early TME reprogramming (Figure 8A). This integrative model captures how transcriptional regulators, immune-stromal programs, and early network-evolution genes converge to remodel the tumor microenvironment at first presentation.

**Figure 8.**
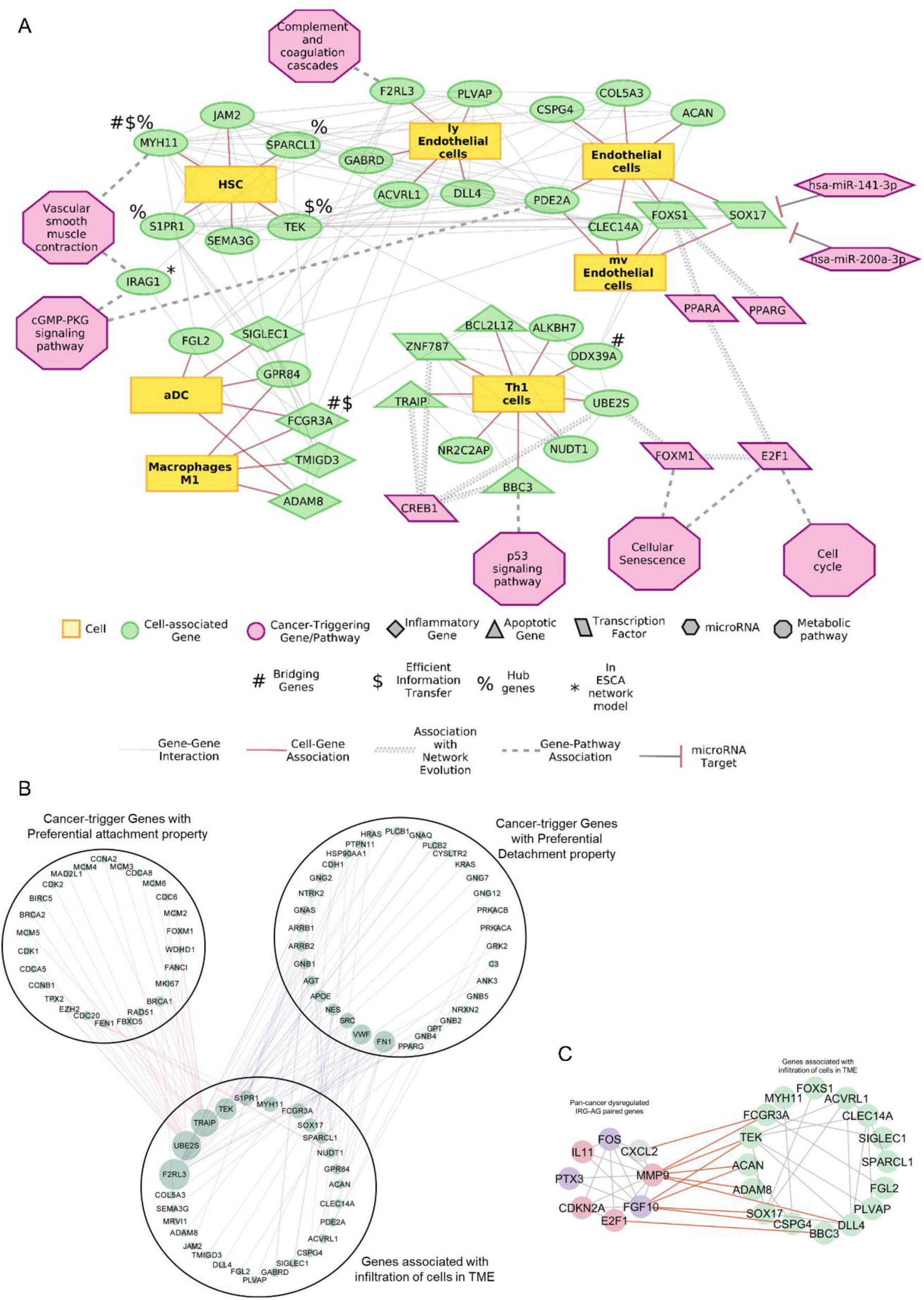
Integrated pan-cancer blueprint of early TME reprogramming and network evolution. A. Schematic blueprint summarizing early TME reprogramming across the 14 cancers. B. PPI network linking TME-infiltrating-cell-associated genes with cancer-trigger genes involved in network evolution. Node size reflects interaction count. Red and blue edges denote associations with preferential attachment and preferential detachment genes, respectively. C. PPI network linking TME-infiltrating-cell-associated genes with pan-cancer dysregulated inflammatory-apoptotic genes involved in network evolution. Red, blue, and grey nodes denote pan-cancer upregulated, downregulated, and differentially expressed inflammatory-apoptotic genes; green nodes denote blueprint genes. Red edges depict interactions involving dysregulated inflammatory-apoptotic genes.

Within this blueprint, UBE2S, a Th1 cell-associated gene, was positioned downstream of FOXM1, which in turn is transcriptionally regulated by E2F1, a core driver of the inflammation-apoptosis axis. CREB1, a preferential-detachment master regulator, targeted multiple Th1-associated genes—TRAIP, ZNF787, BBC3, and UBE2S indicating a direct regulatory link between detachment-type network rewiring and immune infiltration. For endothelial-associated programs, FOXS1 was identified as a transcriptional target of PPARA and PPARG, while PPARA itself was regulated by E2F1, placing endothelial remodeling downstream of the same proliferative axis controlling FOXM1.

Functional mapping revealed that FOXM1 was associated with dysregulated cellular senescence, whereas E2F1 impacted both cellular senescence and the cell cycle. F2RL3 aligned with altered complement and coagulation cascades, MYH11 and IRAG1 with vascular smooth muscle contraction, PDE2A and IRAG1 with cGMP-PKG signaling, and BBC3 with p53 signaling, collectively highlighting that the early TME blueprint integrates vascular, apoptotic, inflammatory, and senescence pathways.

Gene-level functional classes further contextualized this landscape. FOXS1, SOX17, and ZNF787 encoded transcription factors; SIGLEC1, FCGR3A, TMIGD3, and ADAM8 were inflammatory-response genes; BCL2L12, TRAIP, and BBC3 participated in apoptosis. Notably, SOX17 was targeted by hsa-miR-141-3p and hsa-miR-200a-3p, situating miRNA regulation within endothelial-associated TME remodeling.

Network topology underscored MYH11, S1PR1, TEK, and SPARCL1 as high-degree hubs. MYH11, FCGR3A, and TEK demonstrated the highest closeness centrality, consistent with efficient signal propagation, while FCGR3A, MYH11, and DDX39A exhibited the highest betweenness centrality, marking them as critical connectors between TME-infiltration modules (Table 3).

Finally, integration with network-evolution data revealed that the blueprint closely interfaces with early cancer-trigger processes. Twenty-seven TME-infiltration-associated genes interacted with 26 preferential-attachment and 34 preferential-detachment genes. F2RL3 showed the strongest connectivity to detachment-type cancer-trigger genes, whereas UBE2S and TRAIP exhibited the highest interactions with attachment-type triggers. Among detachment-driven genes, FN1 displayed the strongest linkage to Pan-Cancer Blueprint genes (Figure 8B). These blueprint components also aligned with the pan-cancer dysregulated inflammation-apoptosis axis (Figure 8C), positioning the early TME as a direct output of network remodeling and dysregulated immuno-apoptotic circuits.

## Discussion

Cancer initiation represents a systems-level transition in which a previously homeostatic tissue adopts a new organizational state defined by instability, metabolic reprogramming, and inflammatory plasticity. By interrogating first-stage tumors across fourteen TCGA cancers, this work uncovers a conserved, pan-cancer blueprint in which transcriptional reprogramming, metabolic imbalance, inflammatory-apoptotic crosstalk, and network evolution converge to form an early malignant state. Importantly, because all analyses were restricted to stage-I tumors, the observed patterns reflect events proximal to tumor initiation rather than later adaptations acquired during progression. The resulting integrated model identifies a compact set of gene-network triggers centrally FOXM1, PPARA, PPARG, CREB1 and a dense cohort of cell-cycle regulators that collectively drive early cancer network rewiring, metabolic deviation, and inflammatory-apoptotic imbalance.

Across all cancers studied, the gene interaction networks retained a scale-free topology in both normal and first-stage tumor tissues, consistent with Barabási and Albert’s formulation that biological networks evolve via preferential attachment (Barabási and Albert, 1999). Yet the transition from normal to early tumor state involved marked redistributions in connectivity. A small group of genes—CDK1, CCNB1, BRCA1, BIRC5, KIF11, ASPM, CHEK1, FOXM1, TOP2A and CDC6—gained interactions across ten or more cancers, forming a densely interconnected core in which eleven nodes (ASPM, BIRC5, BUB1, CCNA2, CCNB1, CDC20, CDK1, CENPF, KIF11, MAD2L1 and TOP2A) were maximally connected. These genes, uniformly upregulated, collectively define a shared proliferative signal across cancers, supported by enriched pathways in cell cycle, DNA replication, homologous recombination. Their synchronous overexpression and coordinated connectivity strengthening indicate a pan-cancer shift towards a hyper-mitotic state that appears fundamental to malignant inception.

In parallel, a distinct cohort—SRC, AGT, DLG4, APP, PRKACA, FN1, PRKACB, NTRK2, SYP and GNB1—showed consistent loss of interactions, forming a sparse, preferentially detaching network enriched for synaptic and neurotransmission pathways. These genes were neither commonly dysregulated nor influential in pan-cancer networks, indicating functional attenuation rather than active regulation. Their collective down-modulation suggests early suppression of neuronal-like signaling, consistent with literature linking neuro-modulatory circuits to anti-tumor immunity (Amit et al., 2025; Mancusi and Monje, 2023). The concordant decrease of these nodes across cancers supports the interpretation that early malignant ecosystems turn down neural-metabolic cross-talk to insulate the emerging proliferative core. This pattern suggests that cancer network evolution may involve simultaneous strengthening of mitotic hubs and systematic decommissioning of anti-proliferative neuro-modulatory modules.

FOXM1 emerged as the central orchestrator of this attachment subnetwork, interacting with 46 of the 50 preferentially attaching genes. FOXM1’s documented regulation of M-phase progression, DNA repair, immune modulation, metabolic control and therapy resistance (Hsu et al., 2024; Li et al., 2024; Raghuwanshi et al., 2024; Timilsina et al., 2025; Wang et al., 2025) is consistent with this dominant topological positioning. Its pan-cancer upregulation and hub-like influence indicate that FOXM1 is not merely another proliferative transcription factor but functions as the architect of the early malignant network state. The finding that FOXM1 also participates in the pan-cancer upregulated Cellular Senescence pathway highlights its role in subverting senescence-associated defenses, thereby shifting normal tissues toward proliferative immortality.

In contrast, the transcription factors PPARA, PPARG and CREB1 characterized the preferential detachment network. PPARG was pan-cancer downregulated, while PPARA and CREB1 interacted with detaching nodes, consistent with their roles in lipid sensing, apoptosis regulation, T-cell metabolic competence, macrophage polarization and neuromodulation(Li et al., 2023; Rudko et al., 2020; Zeng et al., 2022). Early loss of these regulators deprives normal tissues of lipid-immune homeostasis, contributing to the metabolic and inflammatory plasticity characteristic of tumors. CREB1’s interaction with CCNA2 links inflammatory-metabolic detachment directly to the proliferative engine, suggesting a point of early cross-regulatory vulnerability. The observation that pan-cancer network evolution is driven by simultaneous FOXM1-centered attachment and PPAR/CREB-mediated detachment represents a previously unrecognized dual-axis model of early tumorigenic network reprogramming.

Metabolic pathway analysis further supported this model. GSEA across the fourteen cancers revealed consistent upregulation of Ribosome Biogenesis, proteasome, cell cycle, DNA replication, and multiple DNA repair pathways, demonstrating strong proliferation-centric metabolic orchestration. The downregulated landscape, in contrast, was dominated by fatty acid degradation, valine-leucine-isoleucine degradation, PPAR Signaling, adipocytokine signaling, calcium signaling and drug metabolism via cytochrome p450. These changes demonstrate a broad collapse of lipid catabolism, branched-chain amino acid breakdown, and pharmacologic detoxification—metabolic features compatible with increased lipid storage, amino acid conservation and treatment resistance.

Among all genes, ADH1B was the most consistently downregulated across all fourteen cancers, and fatty acid degradation was the most consistently downregulated pathway. ADH1B’s reduced expression implies aldehyde accumulation, which promotes DNA damage and genomic instability (Woo and Jia, 2024). Its recently described suppression of tumor stemness via cAMP/PKA/CREB1 signaling (Shi et al., 2025) places ADH1B at the intersection of metabolic detoxification, stemness acquisition and CREB1 activity, making it a plausible early metabolic trigger of malignant transformation. The concordant pan-cancer repression of ADH1B and fatty acid degradation suggests that aldehyde-induced genomic instability and lipid accumulation are universal metabolic primers of early tumorigenesis.

Integration of pathway-level and network-level data revealed that ten preferentially attaching cancer-trigger genes (FOXM1, BUB1, CDC6, CDC20, CCNA2, BRCA2, MCM2, RAD51, FANCI and RRM2) directly contribute to dysregulated pathways governing DNA replication fidelity, cell cycle control, senescence and homologous recombination. Their dual role as (i) initiators of network rewiring and (ii) effectors of pathway-level dysregulation confirms them as core, conserved drivers of malignant state acquisition rather than downstream consequences of tissue-specific alterations.

Finally, an integrated analysis of inflammation and apoptosis revealed six pan-cancer dysregulated IRG-AGpairs: CXCL2-MMP9, MMP9-IL11, IL11-FGF10, PTX3-FGF10, FOS-CDKN2A and FOS-E2F1. These eight genes were also among the most influential nodes in their respective networks, demonstrating a consistent early imbalance linking extracellular matrix remodeling (MMP9), chemokine-driven leukocyte trafficking (CXCL2, CXCL5), cytokine-driven proliferative signaling (IL11), pro-proliferative transcriptional regulation (FOS, E2F1), growth factor signaling (FGF10) and tumor-suppressive cell-cycle control (CDKN2A). Their cross-module placement underscores that inflammation-apoptosis crosstalk is not a late-stage adaptation but an early, reproducible property of tumor emergence. The hierarchical clustering based on IRGs also mirrored tissue-of-origin lineages, suggesting that inflammatory dysregulation partially recapitulates germ-layer constraints, whereas apoptotic dysregulation does not—consistent with apoptosis being a more universally hijacked mechanism.

The core IRG-AGgene sets interacted extensively with preferential attachment/detachment regulators, including FOXM1, PPARA, PPARG and CREB1, indicating that metabolic reprogramming, proliferative rewiring and inflammatory-apoptotic imbalance are not isolated events but components of a single early tumorigenic program. This interlocked architecture suggests that early cancer does not evolve via sequential hallmarks but by a coordinated systems-level shift in which proliferation, metabolism and inflammation become mutually reinforcing modules.

Taken together, these findings propose a unifying model of early tumorigenesis in which FOXM1-driven proliferation, PPAR/CREB-governed metabolic-neuromodulatory detachment, ADH1B-associated aldehyde accumulation, suppression of lipid catabolism, and inflammatory-apoptotic disequilibrium converge to form a conserved malignant attractor state. This state is ancestrally encoded and phenotypically plastic, enabling rapid adaptation while maintaining hallmark properties of early cancer. The identification of this tightly interlinked network highlights several potential early intervention targets—most prominently FOXM1, ADH1B and the IL11-MMP9-FGF10 axis—whose modulation may destabilize the nascent malignant program before it becomes clinically entrenched.

### Limitations of the study

Although this study provides a unified pan-cancer blueprint of early tumor microenvironment reprogramming, several limitations warrant consideration. The analysis relies on bulk first-stage TCGA transcriptomes, which constrain cell-type resolution and cannot fully dissect epithelial-stromal heterogeneity. Network-evolution inferences are based on correlation-derived topology and do not establish causal directionality among trigger genes, metabolic regulators, and inflammatory-apoptotic axes.. Pathway-level shifts, particularly in fatty acid degradation, PPAR signaling, and aldehyde metabolism reflect conserved transcriptomic signatures but require functional validation in experimental models. Finally, while the pan-cancer framework highlights robust, cross-tissue convergence, cancer-type-specific nuances may be underrepresented by the emphasis on shared early axes.

## Methods

### Data Acquisition

RNA-seq (raw counts; STAR output) and miRNA-seq data for first-stage tumors and matched normal tissues were retrieved for 14 TCGA cancers (BRCA, CHOL, COAD, ESCA, HNSC, KICH, KIRC, KIRP, LIHC, LUAD, LUSC, READ, STAD, THCA) from the GDC Portal, restricted to tumor types with ≥5 normal and ≥5 stage-I samples [36, 37].

### Network-Evolution Framework: Construction of PPI Networks

Significantly expressed genes in normal and stage-I tissues were identified using edgeR [38] For each cancer, two condition-specific protein-protein interaction (PPI) networks were constructed (STRING via Cytoscape-stringApp) [39-41]. Degree distributions were computed using NetworkAnalyzer [42] to assess scale-free architecture.

To quantify preferential attachment and detachment, genes present in both networks were assigned Δdegree values (cancer−normal). Genes exhibiting increased or decreased degree in ≥10 cancers were averaged across cancers. The top 50 genes with largest mean gain (“attachment set”) and loss (“detachment set”) were used to build summary PPI networks. Transcription-factor regulators were annotated using Harmonizome 3.0 [43, 44]. KEGG pathway enrichment of attachment/detachment genes was performed in DAVID [45-48].

### Differential Expression and Robust Rank Aggregation (RRA) Framework

Differentially expressed genes (DEGs) were computed using DESeq2 [49, 50] applying padj < 0.05 and |log2FC| >1. DEGs of each cancer were separated into up-and down-regulated sets and ranked by log2FC. Robust Rank Aggregation [51] with Benjamini-hochnerg correction [52] generated pan-cancer ranked lists for up- and down-regulated genes. Genes significantly dysregulated (FDR < 0.05) in ≥10 cancers were designated pan-cancer up- and down-regulated genes.

Pan-cancer dysregulated genes were assembled into a PPI network (stringApp), and influential genes were identified using Degree and MCC metrics (NetworkAnalyzer; cytoHubba; [53]). Parallel DEG-specific PPI networks were constructed for each cancer, and Degree/MCC rankings were again subjected to RRA to identify pan-cancer influential genes. These were intersected with genes showing preferential degree gain/loss to obtain influential drivers of network evolution.

### Gene Set Enrichment Analysis and RRA of Pathways

GSEA [54-56] was performed on normalized RNA-seq counts (DESeq2) using KEGG gene sets retrieved via KEGGREST [46]. Dysregulated pathways (|NES|>1, padj < 0.05) were divided into up- and down-pathways and subjected to RRA to identify pan-cancer dysregulated pathways [51]. Pan-cancer dysregulated pathways were mapped to pan-cancer dysregulated genes to identify pathway-linked driver genes.

### Inflammatory-Apoptotic Gene Analysis

Inflammatory-response genes (IRGs) and Apoptosis Genes (AGs) were compiled from QuickGO [57, 58] and MSigDB [59, 60]. Overlap defined IRAGs. DEGs of each cancer were intersected with IRGs/AGs/IRAGs. PPI networks of deIRGs-deAGs were generated (stringApp; score cutoff 0.04). Influential genes were ranked using MCC, and modified networks were used to compute Closeness and Betweenness centralities (cytoHubba; NetworkAnalyzer). Centrality-based gene ranks across cancers were integrated via RRA [51] to obtain pan-cancer influential IRGs and AGs. Their connections with attachment/detachment genes were analyzed to assess network-evolution coupling.

### miRNA Differential Expression, RRA, and Target-Set GSEA

MicroRNA-seq data were retrieved [36, 37], normalized in DESeq2, and DEMs identified (padj < 0.05, |log2FC| >1). DEMs were ranked and subjected to RRA to obtain pan-cancer up- and down-regulated miRNAs.

miRNA-target predictions were obtained from miRWalk [61, 62]. TargetScan [63], miRDB [64], and miRTarBase [65]. Targets predicted/validated by ≥3 resources were retained. Target-set GSEA (fgsea) identified miRNAs with up- or down-regulated target gene sets (miRNA_upTGS, miRNA_dnTGS). Their NES-ranked lists underwent RRA, yielding pan-cancer miRNAs with dysregulated target sets. Overlaps between dysregulated miRNAs and dysregulated target-set miRNAs defined Up_miRNA_Dn_TGS and Dn_miRNA_Up_TGS. Their targets were intersected with pan-cancer DEGs, KEGG RRA pathways, and network-evolution genes.

### Immune/Stromal Cell Infiltration and WGCNA

TPM data were used to compute infiltration scores of 64 cell types using xCell (Aran, Hu, and Butte 2017). Differential infiltration was assessed via t-tests (p ≤0.05).

Consensus WGCNA [66] was applied separately to normal and stage-I cancers (vst-normalized DESeq2 data; networkType=“signed”; β selected by scale-free topology R² ≥0.8; minModuleSize=10) to identify pan-normal modules and pan-cancer modules. Module-trait correlations with xCell infiltration defined pan-normal-infiltrating (PNI) and pan-cancer-infiltrating (PCI) cell types (|r|≥0.5, p<0.05). Individual WGCNAs were then run for each cancer to derive cancer-specific PCI-associated DEGs (csPCI-DEGs), using correlation thresholds |r|≥0.7 (relaxed to ≥0.6 or ≥0.5 when necessary).

### PCI-Gene Network Modeling

A gene-gene interaction (GGI) network for PCI-associated genes was built using GeneMANIA [67]. Influential genes were computed (NetworkAnalyzer). Genes most consistently present across cancers were used to derive a pan-cancer PCI network model, from which cancer-specific subnetworks were obtained. Functional enrichment of network genes was performed in DAVID [48].

### Construction of the Pan-Cancer Blueprint of Early TME Reprogramming

The final blueprint integrated:

1. genes showing preferential attachment/detachment,
2. pan-cancer influential genes,
3. pan-cancer dysregulated DEGs and pathways,
4. pan-cancer influential IRGs/AGs,
5. miRNA regulatory layers,(6) PCI-network architecture.

Master-regulator TF targets (Harmonizome 3.0) were mapped to the TME model, and interactions with cancer-trigger genes were assembled using STRING PPI networks [39, 40].

## Acknowledgments

We gratefully acknowledge the following sources of support: Bhavya for the support from ICMR-JRF and SRF from ICMR, Govt. of India; S.A. for project fellow support from DST-CURIE-2022-80, Government of India; R.M. gratefully acknowledges for the research support from DST-CURIE-2022-80(G), Government of India.

## Authors’ Contribution

Bhavya performed computational and bioinformatics experiments, analyzed data, generated figures, and co-wrote the manuscript. SA analyzed data and generated figures. EP guided the research, cross-checked bioinformatic methods and analyses, co-wrote and edited the manuscript. RM conceived and designed the study, supervised the research work, analyzed data, and wrote the manuscript. All authors have reviewed and approved the final version of the manuscript.

