## Supplementary Material for "A Pan-Cancer Blueprint of Early Tumor Microenvironment Reprogramming"

**Corresponding address:**

*#**Rajeev Mishra, Ph.D.** Bioinformatics Department, MMV, Institute of Science, Banaras Hindu University, Varanasi-221005**, India**

***Ekta Pathak, Ph.D.** , Institute of Diabetes and Obesity, Helmholtz Zentrum, München, Neuherberg, **Germany**

;

**Table S1** Number of significantly expressed genes with degree change in the networks of normal and first-stage cancer conditions of the 14 cancers.

| S. No. | Type of cancer | No. of first-stage cancer tissue samples | No. of normal tissue samples | No. of Significantly Expressed genes in first-stage cancer condition | No. of Significantly Expressed genes in normal condition | Gain/Loss in number of Significantly Expressed Genes in First-stage Cancer | No. of Significantly Expressed genes in both normal and first-stage conditions of a cancer | No. of genes with Decreased Degree in cancer condition | No. of genes with Increased Degree in cancer condition |
|---|---|---|---|---|---|---|---|---|---|
| 1 | Breast invasive carcinoma (BRCA) | 183 | 113 | 14657 | 15080 | Loss of 423 | 14477 | 7102 | 1629 |
| 2 | Cholangiocarcinoma (CHOL) | 18 | 9 | 13638 | 12865 | Gain of 773 | 12419 | 1446 | 8396 |
| 3 | Colon adenocarcinoma (COAD) | 81 | 41 | 13777 | 14712 | Loss of 935 | 13603 | 9017 | 782 |
| 4 | Esophageal carcinoma (ESCA) | 16 | 11 | 13760 | 13184 | Gain of 576 | 12918 | 1146 | 7845 |
| 5 | Head and Neck squamous cell carcinoma (HNSC) | 25 | 44 | 13936 | 14185 | Loss of 249 | 13636 | 4850 | 3364 |
| 6 | Kidney Chromophobe (KICH) | 20 | 25 | 13552 | 14916 | Loss of 1364 | 13447 | 10330 | 335 |
| 7 | Kidney renal clear cell carcinoma (KIRC) | 272 | 72 | 14672 | 14953 | Loss of 281 | 14276 | 5719 | 3552 |
| 8 | Kidney renal papillary cell carcinoma (KIRP) | 172 | 32 | 14241 | 14942 | Loss of 701 | 13960 | 7402 | 2482 |
| 9 | Liver hepatocellular carcinoma (LIHC) | 171 | 50 | 13288 | 13425 | Loss of 137 | 12882 | 4552 | 3921 |
| 10 | Lung adenocarcinoma (LUAD) | 294 | 59 | 14666 | 14548 | Gain of 118 | 14184 | 2798 | 6033 |
| 11 | Lung squamous cell carcinoma (LUSC) | 245 | 49 | 14873 | 15130 | Loss of 257 | 14450 | 5804 | 3733 |
| 12 | Rectum adenocarcinoma (READ) | 30 | 10 | 13164 | 14202 | Loss of 1038 | 12949 | 9185 | 690 |
| 13 | Stomach adenocarcinoma (STAD) | 53 | 32 | 14629 | 14384 | Gain of 245 | 13990 | 2623 | 6628 |
| 14 | Thyroid carcinoma (THCA) | 283 | 59 | 14591 | 14767 | Loss of 176 | 14323 | 5119 | 2828 |

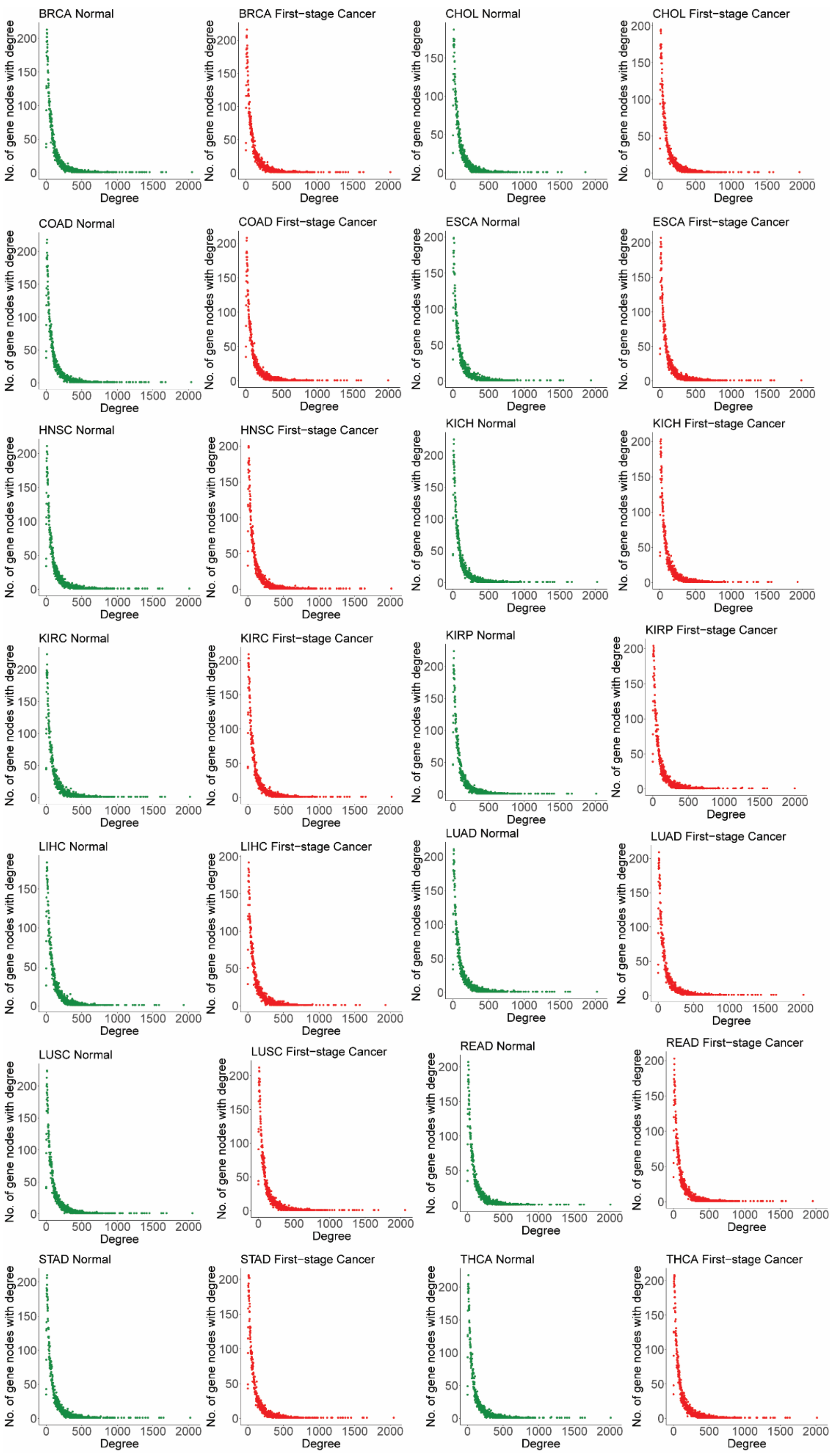


**Figure S1** Degree Distribution of significantly expressed genes in their PPI networks for 14 first-stage cancers and their respective normal conditions

**Table S2** Top 50 genes having the highest average increase in degree in the first-stage cancer

| S. No. | Gene Name | BRCA | CHOL | COAD | ESCA | HNSC | KICH | KIRC | KIRP | LIHC | LUAD | LUSC | READ | STAD | THCA | No. of cancers in which there is an increase of degree in cancer | Average increase in degree |
|---|---|---|---|---|---|---|---|---|---|---|---|---|---|---|---|---|---|
| 1 | CDK1 | 6 | 114 | | 27 | 5 | | 21 | 26 | 65 | 22 | 16 | | 13 | 16 | 11 | 30.09 |
| 2 | CCNB1 | 3 | 106 | | 20 | | | 21 | 23 | 59 | 7 | 9 | | 2 | 14 | 10 | 26.4 |
| 3 | BRCA1 | | 102 | | 18 | 4 | | 16 | 12 | 47 | 13 | 13 | | 12 | 9 | 10 | 24.6 |
| 4 | BIRC5 | | 92 | | 12 | | 4 | 17 | 25 | 56 | 9 | 1 | | 4 | 17 | 10 | 23.7 |
| 5 | KIF11 | 1 | 97 | | 16 | 5 | | 16 | 21 | 52 | 13 | 10 | | 10 | 12 | 11 | 23 |
| 6 | ASPM | | 87 | | 11 | | 2 | 16 | 20 | 50 | 6 | 7 | | 8 | 17 | 10 | 22.4 |
| 7 | CHEK1 | 1 | 99 | | 11 | 1 | | 19 | 22 | 57 | 9 | 8 | | 3 | 16 | 11 | 22.36 |
| 8 | FOXM1 | 1 | 88 | | 9 | | | 14 | 18 | 55 | 8 | 9 | | 8 | 12 | 10 | 22.2 |
| 9 | TOP2A | 2 | 98 | | 15 | 2 | 1 | 18 | 25 | 61 | 12 | 7 | | 6 | 14 | 12 | 21.75 |
| 10 | CDC6 | | 91 | | 11 | 2 | 3 | 17 | 26 | 51 | 6 | 7 | | 4 | 14 | 11 | 21.09 |
| 11 | TPX2 | 1 | 84 | | 7 | | 9 | 17 | 22 | 48 | 3 | 4 | | | 13 | 10 | 20.8 |
| 12 | MAD2L1 | | 88 | | 9 | 2 | | 13 | 19 | 47 | 8 | 9 | | 3 | 9 | 10 | 20.7 |
| 13 | CCNA2 | 2 | | | 25 | 5 | | 25 | 30 | 59 | 12 | 13 | | 13 | 17 | 10 | 20.1 |
| 14 | NCAPH | 3 | 81 | | 7 | | 9 | 17 | 24 | 48 | 6 | 6 | | 1 | 14 | 11 | 19.63 |
| 15 | RACGAP1 | | 81 | | 9 | | 7 | 14 | 17 | 43 | 7 | 1 | | 1 | 14 | 10 | 19.4 |
| 16 | CDCA5 | 1 | 91 | 1 | 8 | | 9 | 17 | 24 | 52 | 5 | 4 | | 2 | 16 | 12 | 19.16 |
| 17 | RRM2 | | 89 | | 7 | 2 | 4 | 16 | 18 | 48 | 7 | 2 | | 1 | 15 | 11 | 19 |
| 18 | KIFC1 | 1 | 77 | | 9 | | 6 | 15 | 20 | 41 | 5 | 2 | | | 14 | 10 | 19 |
| 19 | CENPF | 1 | 90 | 2 | 8 | | 7 | 15 | 22 | 53 | 4 | 4 | | 2 | 14 | 12 | 18.5 |
| 20 | MKI67 | 3 | 85 | | 12 | 1 | 2 | 17 | 25 | 43 | 4 | 6 | | 2 | 16 | 12 | 18 |
| 21 | FEN1 | | 80 | 3 | 8 | 3 | 2 | 14 | 21 | 50 | 6 | 6 | | 4 | 13 | 12 | 17.5 |
| 22 | BUB1 | 6 | | | 14 | | 4 | 20 | 29 | 57 | 11 | 8 | | 6 | 18 | 10 | 17.3 |
| 23 | ZWINT | 1 | 76 | | 6 | 1 | 5 | 13 | 19 | 44 | 3 | 3 | | | 13 | 11 | 16.72 |
| 24 | CDK2 | | 62 | | 17 | 5 | | 14 | 9 | 26 | 11 | 4 | | 12 | 6 | 10 | 16.6 |

| | | | | | | | | | | | | | | | | | |
|---|---|---|---|---|---|---|---|---|---|---|---|---|---|---|---|---|---|
| 25 | MCM4 | 1 | 79 | | 8 | 1 | | 11 | 16 | 44 | 2 | 8 | | 1 | 9 | 11 | 16.36 |
| 26 | SPAG5 | 3 | 83 | 1 | 8 | 1 | 5 | 16 | 22 | 50 | 2 | 5 | | 2 | 13 | 13 | 16.23 |
| 27 | CENPN | 4 | 73 | | 14 | 1 | 4 | 11 | 20 | 39 | 7 | 6 | | 3 | 11 | 12 | 16.08 |
| 28 | KNTC1 | 2 | 72 | | 6 | | 6 | 11 | 17 | 43 | 3 | 3 | | 1 | 12 | 11 | 16 |
| 29 | CDC20 | 5 | | | 15 | 4 | | 19 | 28 | 56 | 7 | 7 | | 5 | 14 | 10 | 16 |
| 30 | ECT2 | 3 | 73 | 1 | 8 | | 6 | 11 | 19 | 43 | 5 | 4 | | 2 | 14 | 12 | 15.75 |
| 31 | CENPU | 3 | 68 | | 11 | | 4 | 10 | 15 | 38 | 6 | 5 | | 2 | 11 | 11 | 15.73 |
| 32 | EZH2 | | 64 | | 24 | 5 | | 6 | 1 | 18 | 10 | 15 | | 12 | 2 | 10 | 15.7 |
| 33 | GINS2 | 1 | 73 | | 6 | 2 | 6 | 11 | 17 | 41 | 2 | 3 | | | 9 | 11 | 15.54 |
| 34 | BRCA2 | 2 | 59 | | 12 | 3 | | 11 | 8 | 29 | 12 | 12 | | 14 | 4 | 11 | 15.09 |
| 35 | PCLAF | 2 | 67 | | 4 | | 4 | 12 | 19 | 40 | 3 | 3 | | 1 | 11 | 11 | 15.09 |
| 36 | CENPW | 4 | 63 | | 12 | | 2 | 8 | 16 | 35 | 6 | 5 | | 3 | 9 | 11 | 14.81 |
| 37 | CDCA8 | 4 | | | 14 | 2 | 2 | 15 | 23 | 58 | 10 | 11 | | 9 | 15 | 11 | 14.81 |
| 38 | ARHGAP11A | 3 | 62 | | 10 | 1 | 5 | 13 | 18 | 40 | 4 | 4 | | 5 | 12 | 12 | 14.75 |
| 39 | MCM6 | 1 | 67 | | 8 | | 3 | 10 | 12 | 33 | 1 | 4 | | | 7 | 10 | 14.6 |
| 40 | RAD51 | 1 | | 4 | 16 | 4 | | 14 | 21 | 57 | 10 | 11 | | 8 | 14 | 11 | 14.54 |
| 41 | FANCI | 1 | 77 | 3 | 9 | 1 | 4 | 12 | 19 | 46 | 6 | 7 | 3 | 3 | 12 | 14 | 14.5 |
| 42 | KNL1 | 1 | 69 | | 7 | 1 | 4 | 9 | 18 | 43 | 5 | 2 | | 2 | 12 | 12 | 14.42 |
| 43 | FBXO5 | 3 | 67 | 1 | 8 | | 4 | 11 | 18 | 37 | 4 | 5 | | 3 | 12 | 12 | 14.42 |
| 44 | MCM3 | | 61 | | 6 | | 2 | 10 | 16 | 31 | 3 | 4 | | 1 | 9 | 10 | 14.3 |
| 45 | MCM2 | 2 | 69 | | 10 | 2 | | 9 | 12 | 33 | 2 | 8 | | 2 | 8 | 11 | 14.27 |
| 46 | SMC2 | 3 | 64 | | 10 | | 2 | 7 | 13 | 35 | 5 | 6 | | 3 | 9 | 11 | 14.27 |
| 47 | MCM5 | | 63 | | 4 | | 4 | 8 | 14 | 32 | 3 | 4 | | 1 | 9 | 10 | 14.2 |
| 48 | NDC80 | 1 | | | 12 | 2 | 7 | 16 | 21 | 51 | 11 | 10 | | 10 | 14 | 11 | 14.09 |
| 49 | WDHD1 | 1 | 67 | | 4 | 1 | 7 | 9 | 18 | 39 | 5 | 3 | | 2 | 12 | 12 | 14 |
| 50 | FAM83D | 1 | 56 | 1 | 6 | | 9 | 13 | 18 | 34 | 2 | 3 | | | 11 | 11 | 14 |

**Table S3** Top 50 genes having the highest average decrease in degree in the first-stage cancer

| S. No. | Gene Name | BRCA | CHOL | COAD | ESCA | HNSC | KICH | KIRC | KIRP | LIHC | LUAD | LUSC | READ | STAD | THCA | No. of cancers in which there is decrease of degree in cancer | Average decrease in degree |
|---|---|---|---|---|---|---|---|---|---|---|---|---|---|---|---|---|---|
| 1 | SRC | 38 | | 87 | | 3 | 85 | 3 | 67 | 47 | | 29 | 92 | | 5 | 10 | 45.6 |
| 2 | AGT | 41 | | 62 | | 10 | 77 | 12 | 69 | 19 | | 36 | 68 | 2 | 11 | 11 | 37 |
| 3 | DLG4 | 26 | | 78 | | 14 | 58 | 22 | 42 | 24 | | 12 | 76 | 22 | 12 | 11 | 35.09 |
| 4 | APP | 30 | | 54 | | 16 | 56 | 28 | 60 | 30 | 9 | 19 | 61 | 11 | 18 | 12 | 32.67 |
| 5 | PRKACA | 36 | | 65 | | 15 | 65 | 15 | 52 | 10 | 4 | 25 | 71 | 5 | 15 | 12 | 31.5 |
| 6 | FN1 | 23 | | 29 | | 2 | 83 | 25 | 60 | 25 | | 13 | 43 | | 10 | 10 | 31.3 |
| 7 | PRKACB | 32 | | 62 | | 15 | 59 | 17 | 49 | 9 | 7 | 24 | 67 | 4 | 13 | 12 | 29.83 |
| 8 | NTRK2 | 26 | | 58 | | 14 | 46 | 14 | 40 | | 7 | 6 | 53 | 15 | 12 | 11 | 26.45 |
| 9 | SYP | 13 | | 61 | | 9 | 35 | 17 | 34 | 22 | 5 | | 57 | 16 | 19 | 11 | 26.18 |
| 10 | GNB1 | 22 | | 56 | | 4 | 36 | | 40 | 6 | | 17 | 62 | 7 | 3 | 10 | 25.3 |
| 11 | GNG2 | 19 | | 54 | | 4 | 33 | | 35 | 10 | | 16 | 56 | 11 | 1 | 10 | 23.9 |
| 12 | CACNA1C | 11 | | 49 | | 4 | 40 | 18 | 38 | | 7 | 15 | 49 | 4 | 11 | 11 | 22.36 |
| 13 | APOE | 13 | 8 | 32 | | 13 | 58 | 25 | 43 | 11 | 3 | 23 | 31 | | 8 | 12 | 22.33 |
| 14 | ARRB2 | 19 | | 44 | | 4 | 37 | | 39 | 20 | 2 | 17 | 45 | 6 | 7 | 11 | 21.81 |
| 15 | KRAS | 22 | | 28 | | 14 | 52 | 14 | 30 | 13 | | | 33 | 3 | 8 | 10 | 21.7 |
| 16 | GNAQ | 26 | | 47 | | 5 | 30 | | 29 | 9 | | 10 | 50 | 3 | 6 | 10 | 21.5 |
| 17 | CDH1 | 21 | | 23 | | 3 | 62 | 22 | 39 | 24 | | 1 | 17 | | 2 | 10 | 21.4 |
| 18 | ARRB1 | 15 | | 42 | | 4 | 34 | | 33 | 18 | 2 | 13 | 44 | 9 | 7 | 11 | 20.09 |
| 19 | SYN1 | 14 | | 52 | | 1 | 34 | 10 | 24 | | 2 | | 43 | 12 | 8 | 10 | 20 |
| 20 | PPARG | 15 | | 26 | | 8 | 48 | 13 | 33 | 7 | 2 | 14 | 30 | | | 10 | 19.6 |
| 21 | NRXN2 | 12 | | 31 | | 5 | 33 | 21 | 34 | 12 | | 5 | 38 | 13 | 11 | 11 | 19.54 |
| 22 | GNAS | 20 | | 37 | | 1 | 31 | 2 | 35 | 13 | 2 | 13 | 40 | 6 | | 11 | 18.18 |
| 23 | GRK2 | 21 | | 33 | | 2 | 33 | | 23 | 12 | 1 | 5 | 35 | | 11 | 10 | 17.6 |

| | | | | | | | | | | | | | | | | | |
|---|---|---|---|---|---|---|---|---|---|---|---|---|---|---|---|---|---|
| 24 | COMT | 19 | 20 | 21 | | 9 | 36 | 15 | 26 | 4 | | 6 | 25 | | 12 | 11 | 17.54 |
| 25 | TRPV1 | 16 | | 39 | | 3 | 30 | 7 | 25 | 8 | | 3 | 41 | 14 | 7 | 11 | 17.54 |
| 26 | ADGRL3 | | | 33 | | 5 | 41 | 6 | | 13 | 3 | 23 | 39 | 2 | 8 | 10 | 17.3 |
| 27 | ACE | 16 | | 29 | | 10 | 34 | 18 | 36 | 8 | 7 | 13 | 24 | 4 | 6 | 12 | 17.08 |
| 28 | GNG7 | 13 | | 43 | | 6 | 25 | 3 | 34 | 8 | 4 | 13 | 42 | 11 | 2 | 12 | 17 |
| 29 | ANK3 | 11 | | 35 | | | 33 | 6 | 22 | 9 | | 3 | 29 | 7 | 12 | 10 | 16.7 |
| 30 | GPT | 16 | 27 | 24 | | 7 | 38 | 10 | 24 | 9 | 6 | 18 | 16 | | 5 | 12 | 16.67 |
| 31 | CREB1 | 18 | | 36 | | 6 | 30 | 5 | 36 | 7 | 2 | 7 | 40 | 3 | 9 | 12 | 16.58 |
| 32 | PLCB2 | 16 | | 35 | | 4 | 31 | 6 | 27 | 8 | | 5 | 31 | 6 | 11 | 11 | 16.36 |
| 33 | GNB2 | 11 | | 37 | | 5 | 25 | 5 | 26 | 6 | | 11 | 43 | 10 | 1 | 11 | 16.36 |
| 34 | PTPN11 | 16 | | 27 | | | 26 | | 15 | 9 | 6 | 13 | 39 | 4 | 8 | 10 | 16.3 |
| 35 | HSP90AA1 | 13 | | 33 | | 11 | 48 | 3 | 20 | 5 | | 5 | 36 | 2 | 3 | 11 | 16.27 |
| 36 | GNB4 | 10 | | 37 | | 6 | 25 | 6 | 27 | 5 | | 11 | 43 | 8 | 1 | 11 | 16.27 |
| 37 | GNB5 | 14 | | 37 | | 5 | 28 | 7 | 28 | 6 | 3 | 15 | 38 | 9 | 5 | 12 | 16.25 |
| 38 | CYSLTR2 | 25 | | 34 | | 3 | 29 | 5 | 28 | 12 | 4 | 13 | | 11 | 14 | 11 | 16.18 |
| 39 | CACNA1D | 5 | | 39 | | | 31 | 13 | 23 | 8 | 5 | 8 | 35 | 2 | 8 | 11 | 16.09 |
| 40 | C3 | 15 | 18 | 24 | | 4 | 29 | 15 | 27 | 11 | | 18 | 29 | 1 | 2 | 12 | 16.08 |
| 41 | CNTNAP2 | 16 | | 33 | 4 | 8 | 24 | 17 | | 15 | 8 | 13 | 34 | 10 | 9 | 12 | 15.92 |
| 42 | GNG12 | 9 | | 38 | | 6 | 25 | 7 | 27 | 7 | 6 | 12 | 40 | 10 | 3 | 12 | 15.83 |
| 43 | ADGRL2 | 3 | | 33 | | 2 | 32 | 9 | 25 | 12 | 5 | 18 | 37 | 2 | 9 | 12 | 15.58 |
| 44 | GSK3B | 13 | | 22 | | 3 | 44 | 14 | 30 | 16 | | 1 | 17 | 1 | 9 | 11 | 15.45 |
| 45 | HRAS | 9 | | 25 | | 10 | 28 | 8 | 21 | 11 | | | 33 | 4 | 5 | 10 | 15.4 |
| 46 | VWF | 11 | | 20 | | 5 | 29 | 8 | 29 | 12 | 7 | 24 | 23 | | 1 | 11 | 15.36 |
| 47 | PLCB1 | 16 | | 34 | | 7 | 29 | 5 | 28 | 5 | 1 | 8 | 30 | 7 | 13 | 12 | 15.25 |
| 48 | PPARA | 14 | 22 | 21 | | 13 | 37 | 5 | 26 | 9 | 4 | 10 | 21 | | 1 | 12 | 15.25 |
| 49 | DLG2 | 13 | | 39 | | 7 | 22 | 13 | 23 | 10 | 3 | 8 | | 18 | 11 | 11 | 15.18 |
| 50 | NES | 9 | | 33 | | 1 | 29 | 8 | 20 | 15 | 2 | | 29 | | 5 | 10 | 15.1 |

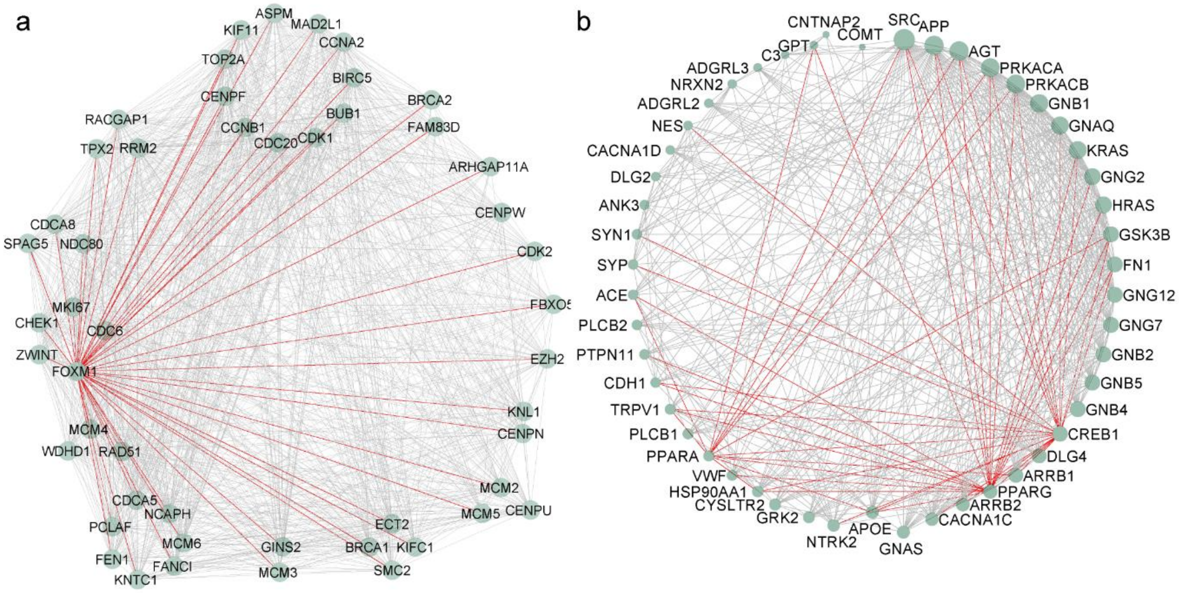


**Figure S2** PPI networks of genes with highest average degree change (a) PPI network between the 50 genes having the highest average increase in degree in the first-stage cancer (b) PPI network between the 50 genes having the highest average decrease in degree in the first-stage cancer. The red edges represent the interaction of genes with the transcription factors

**Table S4** The Degree and Closeness Centrality of 50 genes with highest average degree increase/decrease in their networks

| S. No. | Top 50 genes with highest average increase of degree in cancer | | | Top 50 genes with highest average decrease of degree in cancer | | |
|---|---|---|---|---|---|---|
| | Gene Name | Degree | Closeness Centrality | Gene Name | Degree | Closeness Centrality |
| 1 | ASPM | 49 | 1 | SRC | 34 | 0.765625 |
| 2 | BIRC5 | 49 | 1 | APP | 30 | 0.720588235 |
| 3 | BUB1 | 49 | 1 | AGT | 30 | 0.720588235 |
| 4 | CCNA2 | 49 | 1 | PRKACB | 29 | 0.710144928 |
| 5 | CCNB1 | 49 | 1 | PRKACA | 29 | 0.710144928 |
| 6 | CDC20 | 49 | 1 | GNB1 | 28 | 0.7 |
| 7 | CDK1 | 49 | 1 | KRAS | 27 | 0.680555556 |
| 8 | CENPF | 49 | 1 | GNAQ | 27 | 0.690140845 |
| 9 | KIF11 | 49 | 1 | GNG2 | 25 | 0.671232877 |
| 10 | MAD2L1 | 49 | 1 | HRAS | 25 | 0.662162162 |
| 11 | TOP2A | 49 | 1 | GSK3B | 23 | 0.653333333 |
| 12 | RACGAP1 | 48 | 0.98 | CREB1* | 23 | 0.644736842 |
| 13 | RRM2 | 48 | 0.98 | GNB2 | 23 | 0.653333333 |
| 14 | TPX2 | 48 | 0.98 | GNB5 | 23 | 0.653333333 |
| 15 | CDCA8 | 47 | 0.960784314 | GNG7 | 23 | 0.653333333 |
| 16 | NDC80 | 47 | 0.960784314 | GNG12 | 23 | 0.653333333 |
| 17 | SPAG5 | 47 | 0.960784314 | GNB4 | 23 | 0.653333333 |
| 18 | CDC6 | 46 | 0.942307692 | FN1 | 23 | 0.653333333 |
| 19 | CHEK1 | 46 | 0.942307692 | ARRB1 | 21 | 0.636363636 |
| 20 | FOXM1* | 46 | 0.942307692 | DLG4 | 21 | 0.636363636 |
| 21 | MKI67 | 46 | 0.942307692 | PPARG* | 20 | 0.620253165 |
| 22 | ZWINT | 46 | 0.942307692 | ARRB2 | 19 | 0.620253165 |
| 23 | MCM4 | 45 | 0.924528302 | CACNA1C | 18 | 0.6125 |
| 24 | RAD51 | 45 | 0.924528302 | GNAS | 17 | 0.576470588 |
| 25 | WDHD1 | 45 | 0.924528302 | APOE | 17 | 0.597560976 |
| 26 | CDCA5 | 44 | 0.907407407 | NTRK2 | 16 | 0.590361446 |
| 27 | FANCI | 44 | 0.907407407 | CYSLTR2 | 15 | 0.569767442 |
| 28 | FEN1 | 44 | 0.907407407 | GRK2 | 15 | 0.569767442 |
| 29 | KNTC1 | 44 | 0.907407407 | HSP90AA1 | 14 | 0.569767442 |
| 30 | MCM6 | 44 | 0.907407407 | PPARA* | 13 | 0.550561798 |
| 31 | NCAPH | 44 | 0.907407407 | VWF | 13 | 0.556818182 |
| 32 | PCLAF | 44 | 0.907407407 | TRPV1 | 13 | 0.569767442 |
| 33 | GINS2 | 43 | 0.890909091 | PLCB1 | 13 | 0.544444444 |
| 34 | MCM3 | 43 | 0.890909091 | PTPN11 | 12 | 0.550561798 |
| 35 | BRCA1 | 42 | 0.875 | SYN1 | 12 | 0.556818182 |
| 36 | ECT2 | 42 | 0.875 | SYP | 12 | 0.556818182 |
| 37 | KIFC1 | 42 | 0.875 | ACE | 12 | 0.544444444 |
| 38 | SMC2 | 42 | 0.875 | CDH1 | 12 | 0.538461538 |

| | | | | | | |
|---|---|---|---|---|---|---|
| 39 | CENPU | 41 | 0.859649123 | PLCB2 | 12 | 0.538461538 |
| 40 | MCM2 | 41 | 0.859649123 | ANK3 | 11 | 0.544444444 |
| 41 | MCM5 | 41 | 0.859649123 | ADGRL2 | 10 | 0.538461538 |
| 42 | CENPN | 39 | 0.830508475 | CACNA1D | 10 | 0.5 |
| 43 | KNL1 | 39 | 0.830508475 | DLG2 | 10 | 0.544444444 |
| 44 | EZH2 | 37 | 0.803278689 | NES | 10 | 0.532608696 |
| 45 | FBXO5 | 36 | 0.790322581 | NRXN2 | 10 | 0.510416667 |
| 46 | CDK2 | 34 | 0.765625 | ADGRL3 | 9 | 0.485148515 |
| 47 | CENPW | 33 | 0.753846154 | C3 | 8 | 0.52688172 |
| 48 | ARHGAP11A | 28 | 0.7 | GPT | 8 | 0.505154639 |
| 49 | BRCA2 | 25 | 0.671232877 | CNTNAP2 | 5 | 0.462264151 |
| 50 | FAM83D | 25 | 0.671232877 | COMT | 4 | 0.449541284 |

*Transcription Factor

**Table S5** Number of DEGs between first-stage cancer and normal tissue sample data for 14 TCGA cancers

| S. No. | Type of cancer | No. of DEGs | No. of Upregulated DEGs | No. of Downregulated DEGs |
|---|---|---|---|---|
| 1. | BRCA | 11030 | 7061 | 3969 |
| 2. | CHOL | 7053 | 4538 | 2515 |
| 3. | COAD | 14057 | 9981 | 4076 |
| 4. | ESCA | 1818 | 802 | 1016 |
| 5. | HNSC | 7175 | 3483 | 3692 |
| 6. | KICH | 11146 | 4607 | 6539 |
| 7. | KIRC | 17942 | 14184 | 3758 |
| 8. | KIRP | 10642 | 6404 | 4238 |
| 9. | LIHC | 7772 | 6178 | 1594 |
| 10. | LUAD | 16000 | 13097 | 2903 |
| 11. | LUSC | 16218 | 11023 | 5195 |
| 12. | READ | 9902 | 5344 | 4558 |
| 13. | STAD | 10431 | 7178 | 3253 |
| 14. | THCA | 6345 | 3244 | 3101 |

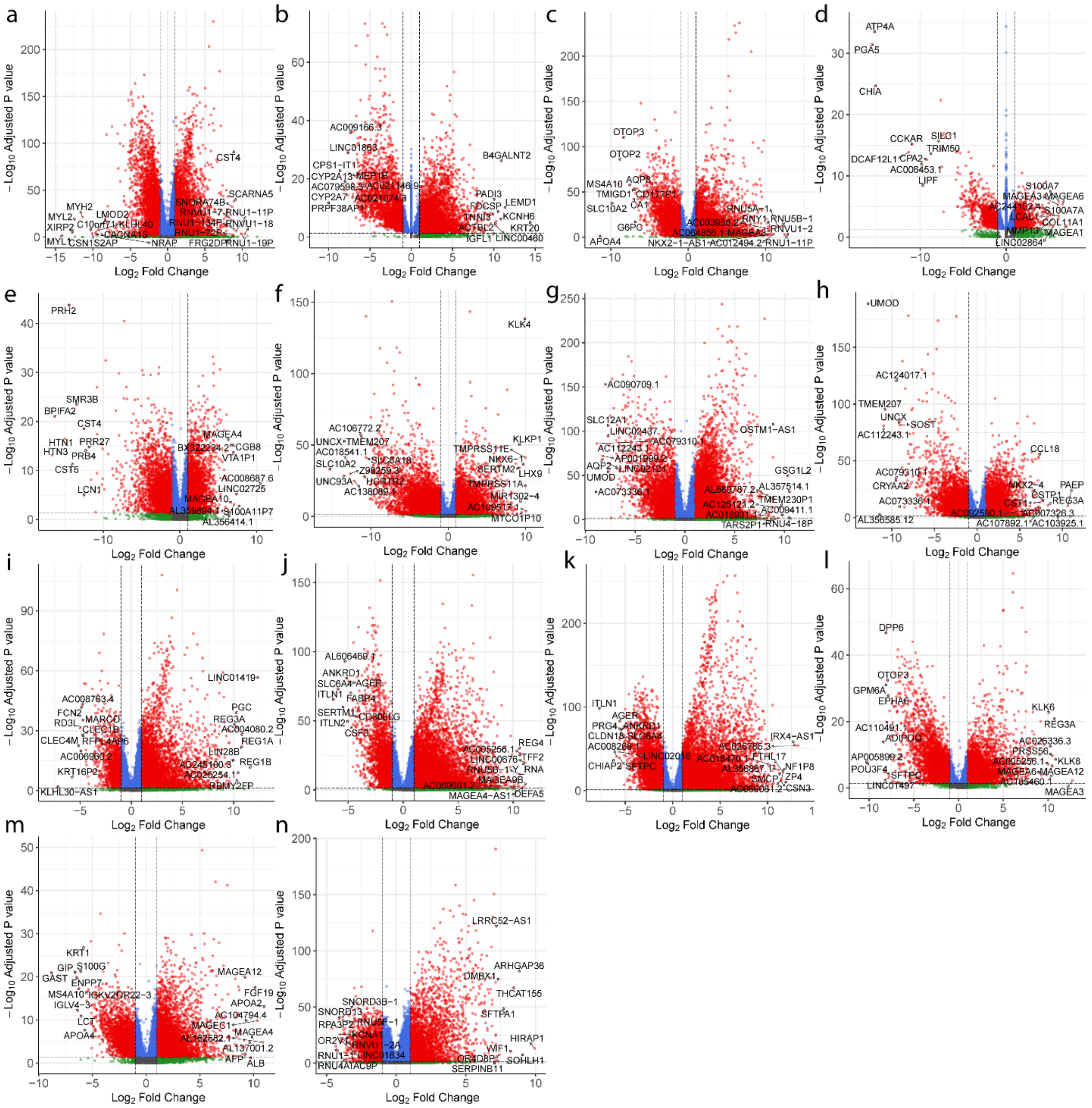

**Figure S3** Volcano plots for the differential gene expression analysis results of the 14 cancers (a) BRCA (b) CHOL (c) COAD (d) ESCA (e) HNSC (f) KICH (g) KIRC (h) KIRP (i) LIHC (j) LUAD (k) LUSC (l) READ (m) STAD (n) THCA. The red dots indicate the statistically significant DEGs (|log2FC| > 1 and padj < 0.05) between normal and first-stage cancer. The green dots indicate genes with padj >=0.05. The blue dots indicate the genes with |log2FC| <= 1. The grey dots represent the genes with |log2FC| < 1 and padj >= 0.05. The gene names of top 10 upregulated and downregulated genes for each cancer are mentioned in the plots.

**Table S6** Number of protein-coding DEGs in the 14 cancers and their PPI networks

| S. No. | Cancer | No. of protein-coding DEGs | No. of protein-coding DEGs in the PPI network |
|---|---|---|---|
| 1 | BRCA | 4761 | 4730 |
| 2 | CHOL | 4867 | 4851 |
| 3 | COAD | 5374 | 5321 |
| 4 | ESCA | 1173 | 1167 |
| 5 | HNSC | 3802 | 3776 |
| 6 | KICH | 5298 | 5257 |
| 7 | KIRC | 5545 | 5450 |
| 8 | KIRP | 4793 | 4735 |
| 9 | LIHC | 3779 | 3739 |
| 10 | LUAD | 5286 | 5234 |
| 11 | LUSC | 6881 | 6835 |
| 12 | READ | 5441 | 5396 |
| 13 | STAD | 4070 | 4042 |
| 14 | THCA | 2869 | 2848 |

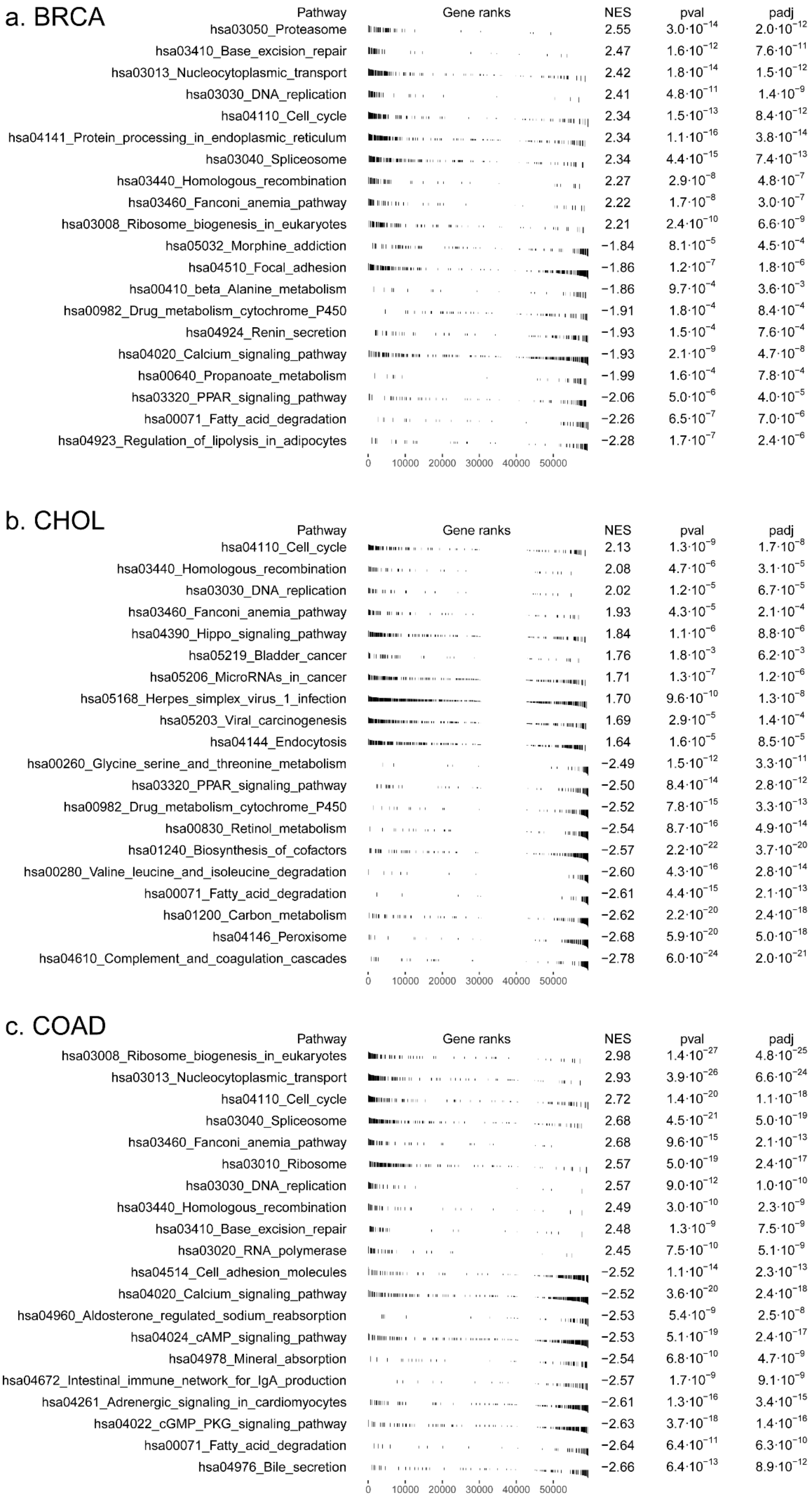


**Figure S4** GSEA tables of 14 cancers representing the top 10 upregulated and downregulated KEGG Pathways (a) BRCA (b) CHOL (c) COAD (d) ESCA (e) HNSC (f) KICH (g) KIRC (h) KIRP (i) LIHC (j) LUAD (k) LUSC (l) READ (m) STAD (n) THCA (Detailed results in Supplementary Table S2_1)

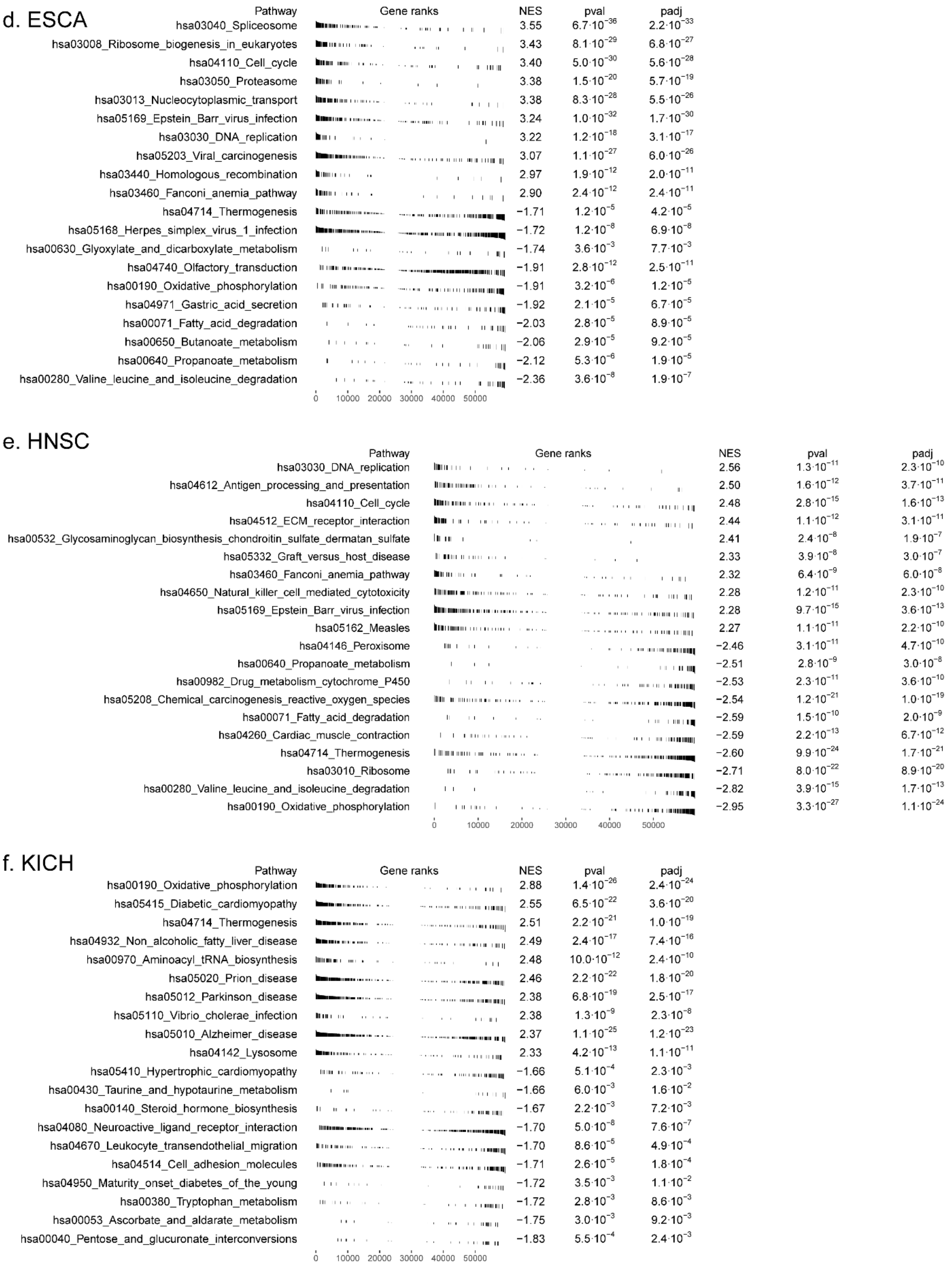


**Figure S4** (continued). GSEA tables of 14 cancers representing the top 10 upregulated and downregulated KEGG Pathways. (a) BRCA (b) CHOL (c) COAD (d) ESCA (e) HNSC (f) KICH (g) KIRC (h) KIRP (i) LIHC (j) LUAD (k) LUSC (l) READ (m) STAD (n) THCA (Detailed results in Supplementary Table S2_1)

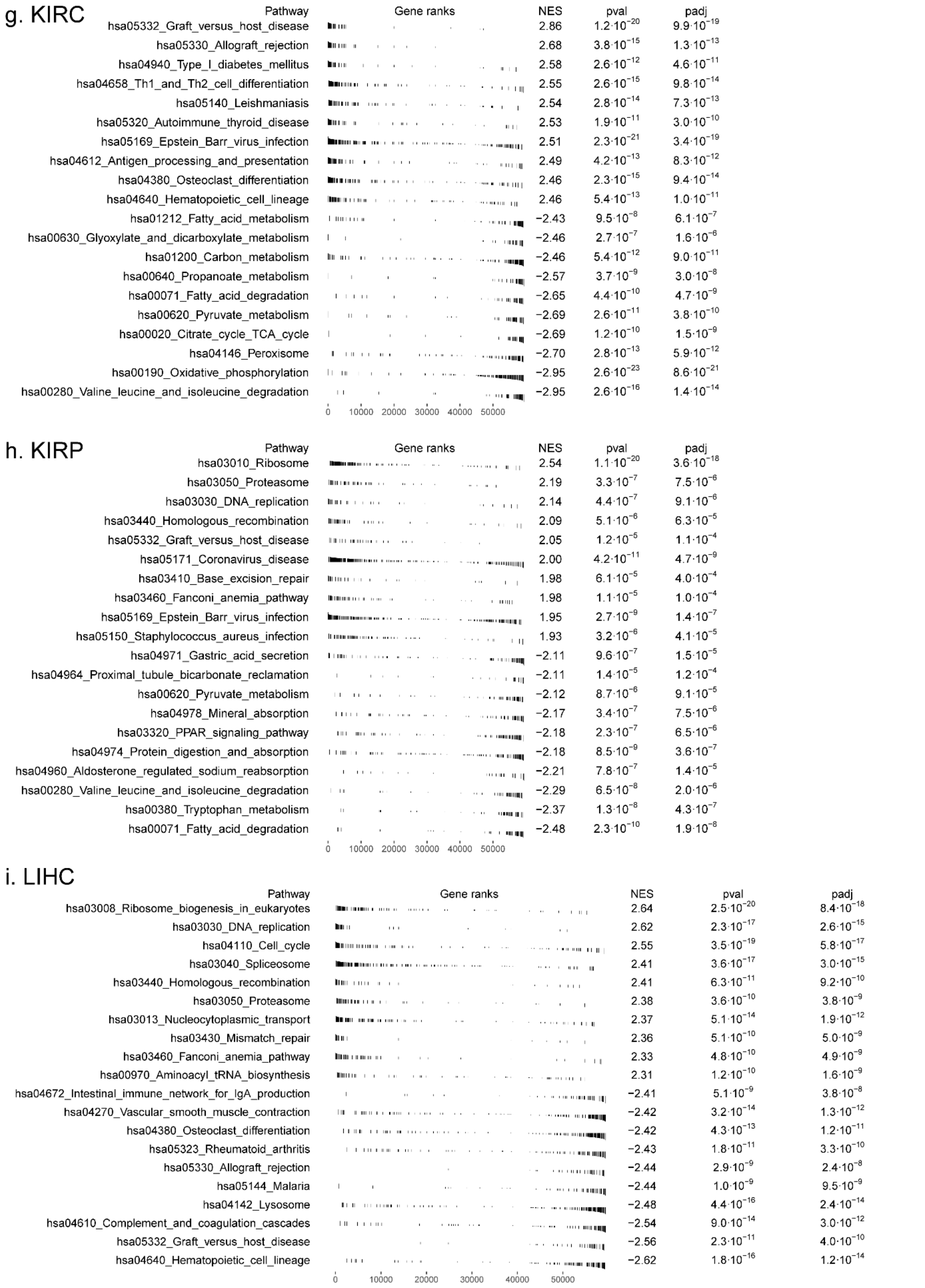


**Figure S4** (continued). GSEA tables of 14 cancers representing the top 10 upregulated and downregulated KEGG Pathways. (a) BRCA (b) CHOL (c) COAD (d) ESCA (e) HNSC (f) KICH (g) KIRC (h) KIRP (i) LIHC (j) LUAD (k) LUSC (l) READ (m) STAD (n) THCA

(Detailed results in Supplementary Table S2_1)

j. LUAD

| Pathway | NES | pval | padj |
|---|---|---|---|
| hsa03030_DNA_replication | 2.57 | $1.2\cdot10^{-13}$ | $2.4\cdot10^{-12}$ |
| hsa03008_Ribosome_biogenesis_in_eukaryotes | 2.52 | $6.7\cdot10^{-16}$ | $3.7\cdot10^{-14}$ |
| hsa03460_Fanconi_anemia_pathway | 2.52 | $7.2\cdot10^{-13}$ | $1.1\cdot10^{-11}$ |
| hsa03440_Homologous_recombination | 2.51 | $6.9\cdot10^{-12}$ | $8.9\cdot10^{-11}$ |
| hsa03013_Nucleocytoplasmic_transport | 2.40 | $2.8\cdot10^{-13}$ | $4.5\cdot10^{-12}$ |
| hsa04110_Cell_cycle | 2.39 | $7.1\cdot10^{-14}$ | $1.7\cdot10^{-12}$ |
| hsa00970_Aminoacyl_tRNA_biosynthesis | 2.30 | $1.2\cdot10^{-9}$ | $9.1\cdot10^{-9}$ |
| hsa03050_Proteasome | 2.26 | $4.2\cdot10^{-8}$ | $2.1\cdot10^{-7}$ |
| hsa03410_Base_excision_repair | 2.26 | $5.8\cdot10^{-8}$ | $2.7\cdot10^{-7}$ |
| hsa01232_Nucleotide_metabolism | 2.22 | $5.0\cdot10^{-9}$ | $3.1\cdot10^{-8}$ |
| hsa04514_Cell_adhesion_molecules | −2.55 | $7.5\cdot10^{-15}$ | $2.3\cdot10^{-13}$ |
| hsa04670_Leukocyte_transendothelial_migration | −2.56 | $2.5\cdot10^{-12}$ | $3.5\cdot10^{-11}$ |
| hsa04923_Regulation_of_lipolysis_in_adipocytes | −2.56 | $2.3\cdot10^{-9}$ | $1.5\cdot10^{-8}$ |
| hsa05332_Graft_versus_host_disease | −2.59 | $1.0\cdot10^{-8}$ | $5.9\cdot10^{-8}$ |
| hsa04380_Osteoclast_differentiation | −2.60 | $7.6\cdot10^{-14}$ | $1.7\cdot10^{-12}$ |
| hsa04640_Hematopoietic_cell_lineage | −2.66 | $3.9\cdot10^{-14}$ | $1.0\cdot10^{-12}$ |
| hsa05144_Malaria | −2.68 | $2.2\cdot10^{-11}$ | $2.5\cdot10^{-10}$ |
| hsa04270_Vascular_smooth_muscle_contraction | −2.71 | $1.4\cdot10^{-16}$ | $9.2\cdot10^{-15}$ |
| hsa04924_Renin_secretion | −2.73 | $2.6\cdot10^{-13}$ | $4.4\cdot10^{-12}$ |
| hsa04611_Platelet_activation | −2.74 | $6.6\cdot10^{-17}$ | $6.6\cdot10^{-15}$ |

k. LUSC

| Pathway | NES | pval | padj |
|---|---|---|---|
| hsa03008_Ribosome_biogenesis_in_eukaryotes | 2.64 | $2.5\cdot10^{-20}$ | $8.4\cdot10^{-18}$ |
| hsa03030_DNA_replication | 2.62 | $2.3\cdot10^{-17}$ | $2.6\cdot10^{-15}$ |
| hsa04110_Cell_cycle | 2.55 | $3.5\cdot10^{-19}$ | $5.8\cdot10^{-17}$ |
| hsa03040_Spliceosome | 2.41 | $3.6\cdot10^{-17}$ | $3.0\cdot10^{-15}$ |
| hsa03440_Homologous_recombination | 2.41 | $6.3\cdot10^{-11}$ | $9.2\cdot10^{-10}$ |
| hsa03050_Proteasome | 2.38 | $3.6\cdot10^{-10}$ | $3.8\cdot10^{-9}$ |
| hsa03013_Nucleocytoplasmic_transport | 2.37 | $5.1\cdot10^{-14}$ | $1.9\cdot10^{-12}$ |
| hsa03430_Mismatch_repair | 2.36 | $5.1\cdot10^{-10}$ | $5.0\cdot10^{-9}$ |
| hsa03460_Fanconi_anemia_pathway | 2.33 | $4.8\cdot10^{-10}$ | $4.9\cdot10^{-9}$ |
| hsa00970_Aminoacyl_tRNA_biosynthesis | 2.31 | $1.2\cdot10^{-10}$ | $1.6\cdot10^{-9}$ |
| hsa04672_Intestinal_immune_network_for_IgA_production | −2.41 | $5.1\cdot10^{-9}$ | $3.8\cdot10^{-8}$ |
| hsa04270_Vascular_smooth_muscle_contraction | −2.42 | $3.2\cdot10^{-14}$ | $1.3\cdot10^{-12}$ |
| hsa04380_Osteoclast_differentiation | −2.42 | $4.3\cdot10^{-13}$ | $1.2\cdot10^{-11}$ |
| hsa05323_Rheumatoid_arthritis | −2.43 | $1.8\cdot10^{-11}$ | $3.3\cdot10^{-10}$ |
| hsa05330_Allograft_rejection | −2.44 | $2.9\cdot10^{-9}$ | $2.4\cdot10^{-8}$ |
| hsa05144_Malaria | −2.44 | $1.0\cdot10^{-9}$ | $9.5\cdot10^{-9}$ |
| hsa04142_Lysosome | −2.48 | $4.4\cdot10^{-16}$ | $2.4\cdot10^{-14}$ |
| hsa04610_Complement_and_coagulation_cascades | −2.54 | $9.0\cdot10^{-14}$ | $3.0\cdot10^{-12}$ |
| hsa05332_Graft_versus_host_disease | −2.56 | $2.3\cdot10^{-11}$ | $4.0\cdot10^{-10}$ |
| hsa04640_Hematopoietic_cell_lineage | −2.62 | $1.8\cdot10^{-16}$ | $1.2\cdot10^{-14}$ |

l. READ

| Pathway | NES | pval | padj |
|---|---|---|---|
| hsa03008_Ribosome_biogenesis_in_eukaryotes | 2.59 | $4.0\cdot10^{-17}$ | $1.3\cdot10^{-14}$ |
| hsa00970_Aminoacyl_tRNA_biosynthesis | 2.38 | $1.6\cdot10^{-10}$ | $7.5\cdot10^{-9}$ |
| hsa03030_DNA_replication | 2.33 | $1.2\cdot10^{-8}$ | $3.0\cdot10^{-7}$ |
| hsa03013_Nucleocytoplasmic_transport | 2.29 | $3.4\cdot10^{-11}$ | $2.3\cdot10^{-9}$ |
| hsa04110_Cell_cycle | 2.23 | $2.7\cdot10^{-10}$ | $10.0\cdot10^{-9}$ |
| hsa03460_Fanconi_anemia_pathway | 2.16 | $4.6\cdot10^{-7}$ | $5.7\cdot10^{-6}$ |
| hsa03050_Proteasome | 2.13 | $1.7\cdot10^{-6}$ | $1.8\cdot10^{-5}$ |
| hsa03020_RNA_polymerase | 2.12 | $6.7\cdot10^{-6}$ | $5.6\cdot10^{-5}$ |
| hsa03040_Spliceosome | 2.10 | $8.4\cdot10^{-10}$ | $2.8\cdot10^{-8}$ |
| hsa03440_Homologous_recombination | 2.05 | $8.2\cdot10^{-6}$ | $5.9\cdot10^{-5}$ |
| hsa04924_Renin_secretion | −2.02 | $2.7\cdot10^{-6}$ | $2.7\cdot10^{-5}$ |
| hsa04024_cAMP_signaling_pathway | −2.02 | $4.3\cdot10^{-11}$ | $2.4\cdot10^{-9}$ |
| hsa00071_Fatty_acid_degradation | −2.03 | $2.7\cdot10^{-5}$ | $1.5\cdot10^{-4}$ |
| hsa00280_Valine_leucine_and_isoleucine_degradation | −2.03 | $8.5\cdot10^{-6}$ | $5.9\cdot10^{-5}$ |
| hsa04080_Neuroactive_ligand_receptor_interaction | −2.04 | $3.5\cdot10^{-15}$ | $3.9\cdot10^{-13}$ |
| hsa04514_Cell_adhesion_molecules | −2.04 | $1.5\cdot10^{-9}$ | $4.5\cdot10^{-8}$ |
| hsa04713_Circadian_entrainment | −2.07 | $5.0\cdot10^{-8}$ | $9.2\cdot10^{-7}$ |
| hsa04261_Adrenergic_signaling_in_cardiomyocytes | −2.09 | $2.5\cdot10^{-10}$ | $10.0\cdot10^{-9}$ |
| hsa04022_cGMP_PKG_signaling_pathway | −2.12 | $1.9\cdot10^{-11}$ | $1.6\cdot10^{-9}$ |
| hsa04020_Calcium_signaling_pathway | −2.17 | $1.0\cdot10^{-15}$ | $1.7\cdot10^{-13}$ |

**Figure S4** (continued). GSEA tables of 14 cancers representing the top 10 upregulated and downregulated KEGG Pathways. (a) BRCA (b) CHOL (c) COAD (d) ESCA (e) HNSC (f)

KICH (g) KIRC (h) KIRP (i) LIHC (j) LUAD (k) LUSC (l) READ (m) STAD (n) THCA (Detailed results in Supplementary Table S2_1)

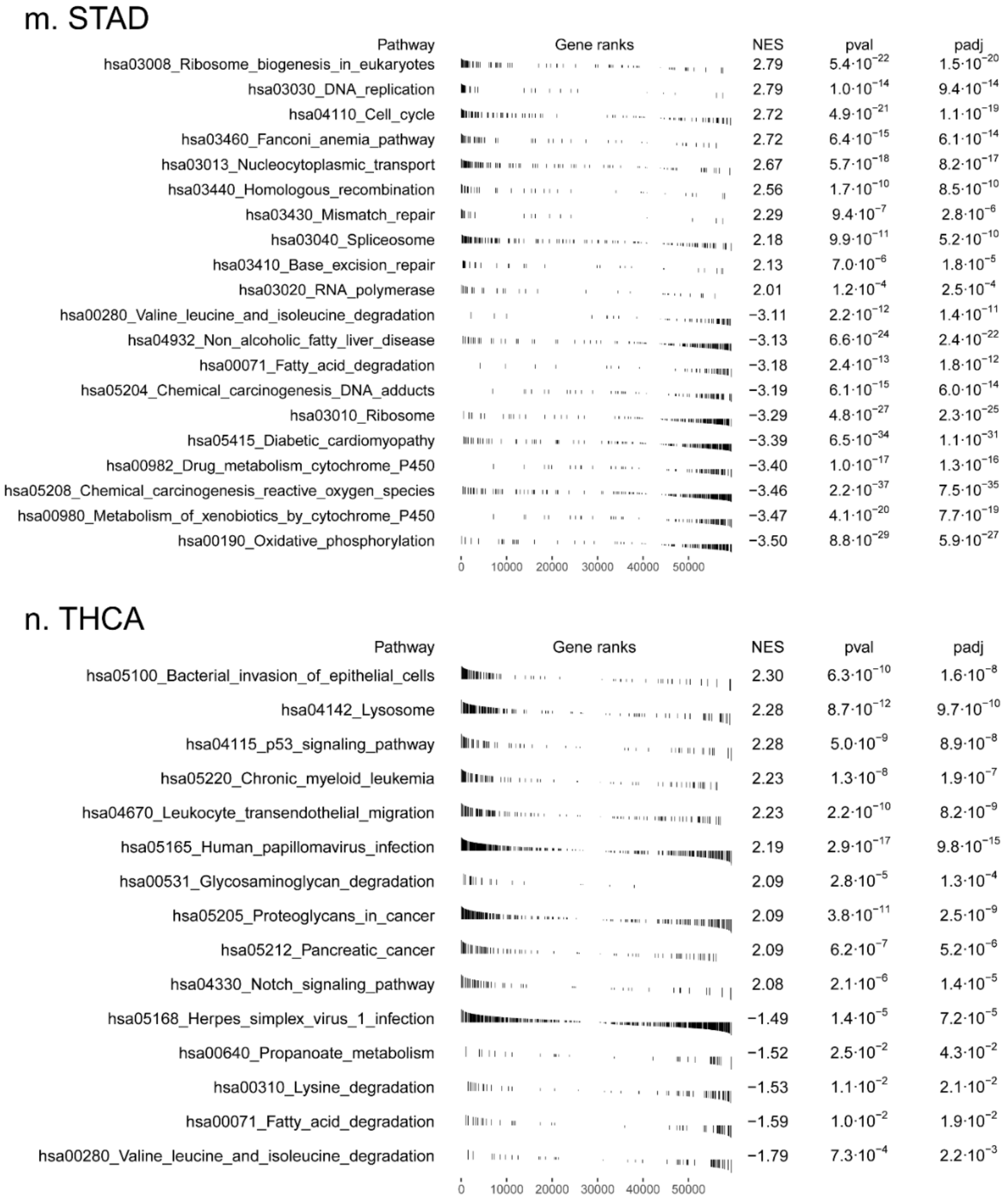


**Figure S4** (continued). GSEA tables of 14 cancers representing the top 10 upregulated and downregulated KEGG Pathways. (a) BRCA (b) CHOL (c) COAD (d) ESCA (e) HNSC (f) KICH (g) KIRC (h) KIRP (i) LIHC (j) LUAD (k) LUSC (l) READ (m) STAD (n) THCA (Detailed results in Supplementary Table S2_1)

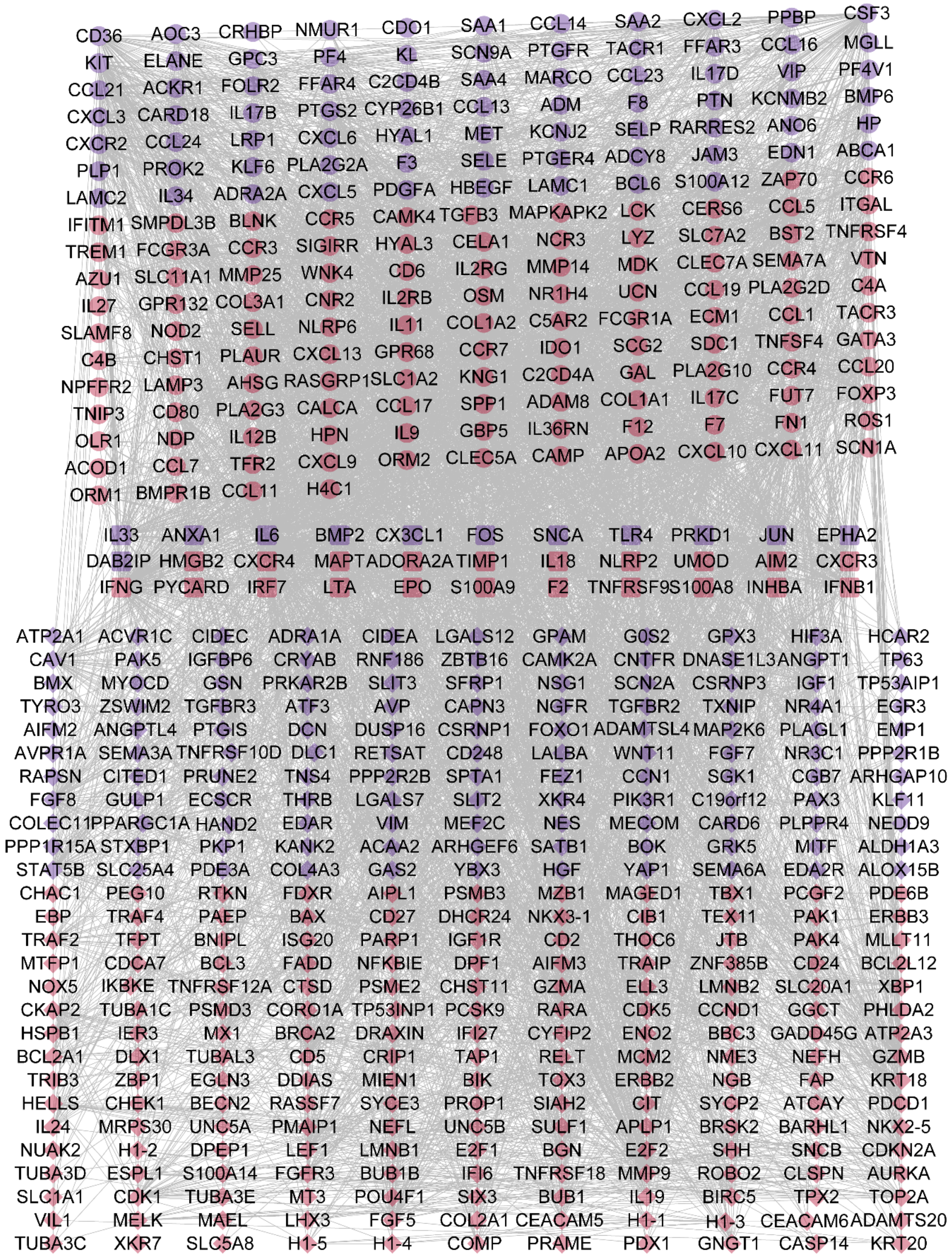


**Figure S5** PPI network between the deIRGs and deAGs in first-stage BRCA . The round nodes located at the upper section of the network represent the IRGs, the diamond-shaped nodes located at the lower section of the network represent the AGs and the rounded-square nodes located at the middle section of the network represent the genes that are IRGs as well as AGs. The red and blue nodes represent the upregulated and downregulated genes, respectively, in first-stage BRCA.

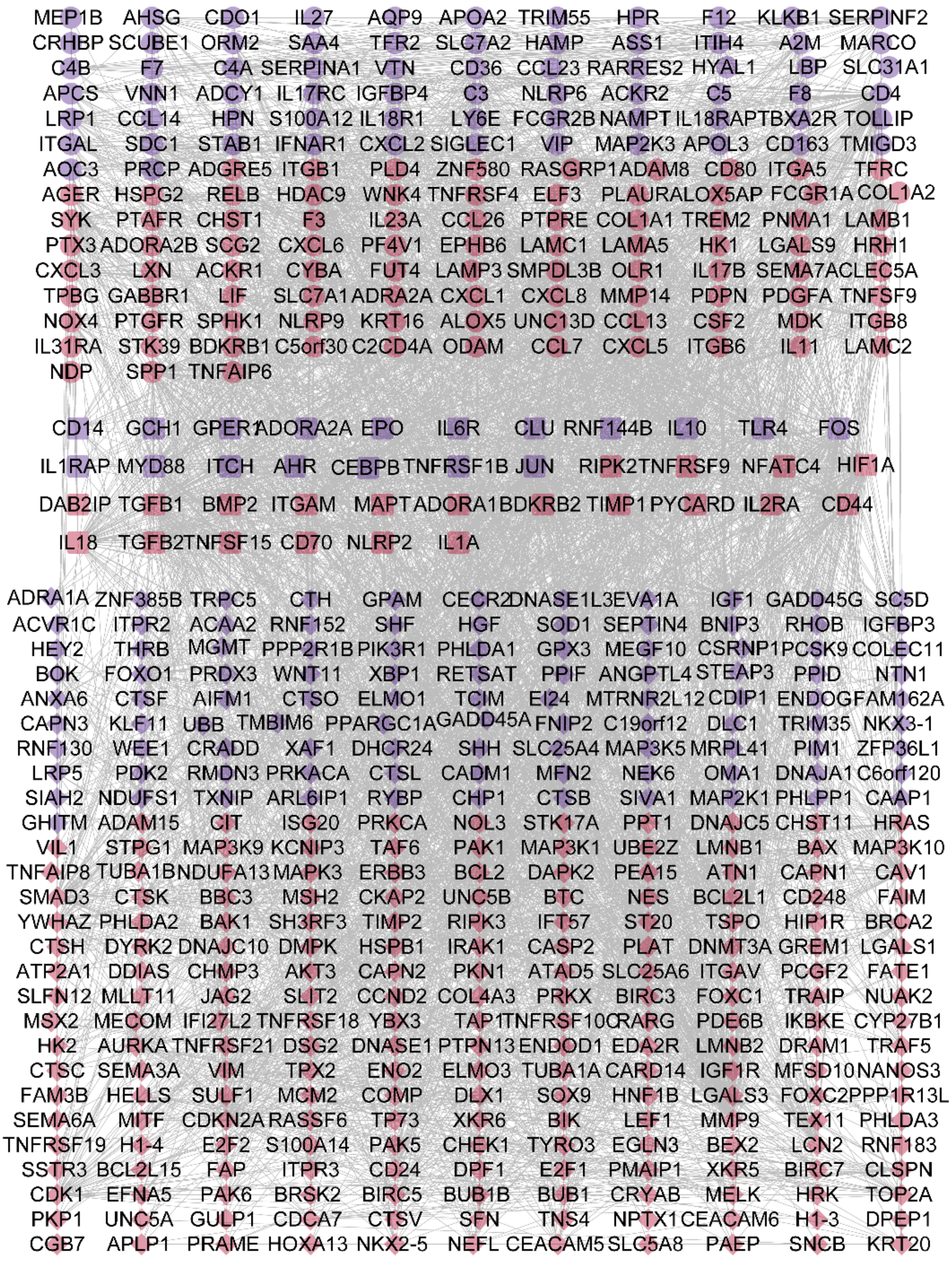


**Figure S6** PPI network between the deIRGs and deAGs in first-stage CHOL . The round nodes located at the upper section of the network represent the IRGs, the diamond-shaped nodes located at the lower section of the network represent the AGs and the rounded-square nodes located at the middle section of the network represent the genes that are IRGs as well as AGs. The red and blue nodes represent the upregulated and downregulated genes, respectively, in first-stage CHOL.

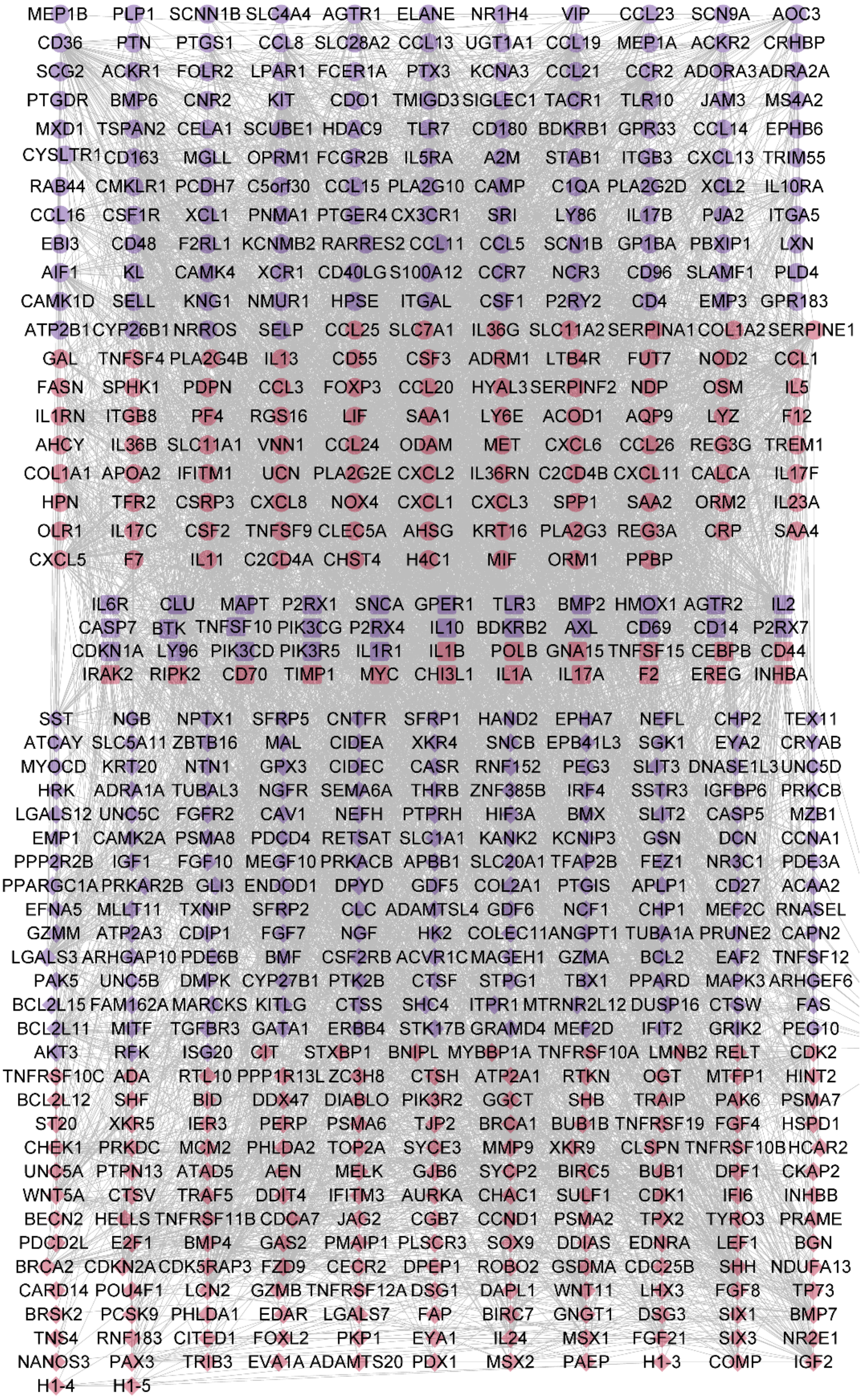


**Figure S7** PPI network between the deIRGs and deAGs in first-stage COAD . The round nodes located at the upper section of the network represent the IRGs, the diamond-shaped nodes located at the lower section of the network represent the AGs and the rounded-square nodes located at the middle section of the network represent the genes that are IRGs as well as AGs. The red and blue nodes represent the upregulated and downregulated genes, respectively, in first-stage COAD.

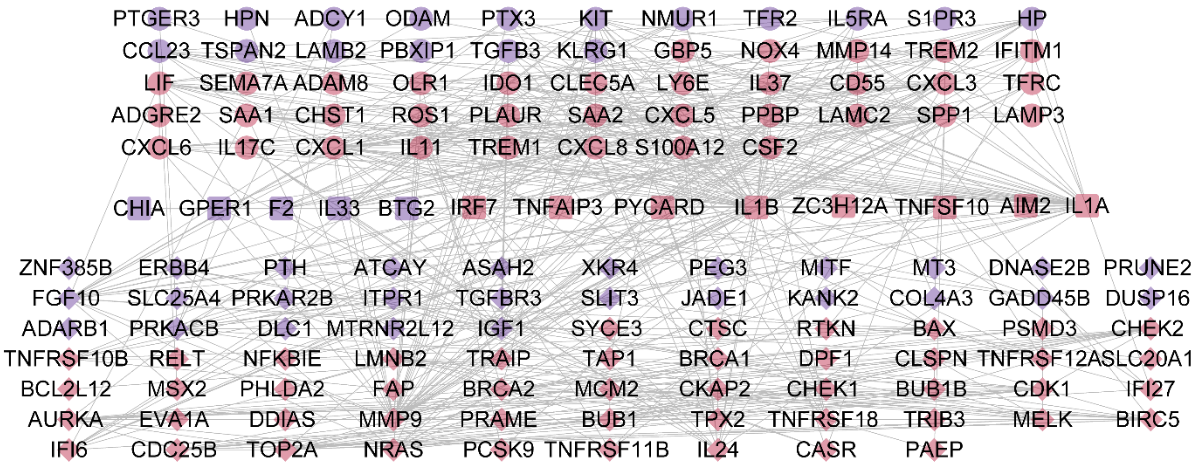


**Figure S8** PPI network between the deIRGs and deAGs in first-stage ESCA . The round nodes located at the upper section of the network represent the IRGs, the diamond-shaped nodes located at the lower section of the network represent the AGs and the rounded-square nodes located at the middle section of the network represent the genes that are IRGs as well as AGs. The red and blue nodes represent the upregulated and downregulated genes, respectively, in first-stage ESCA.

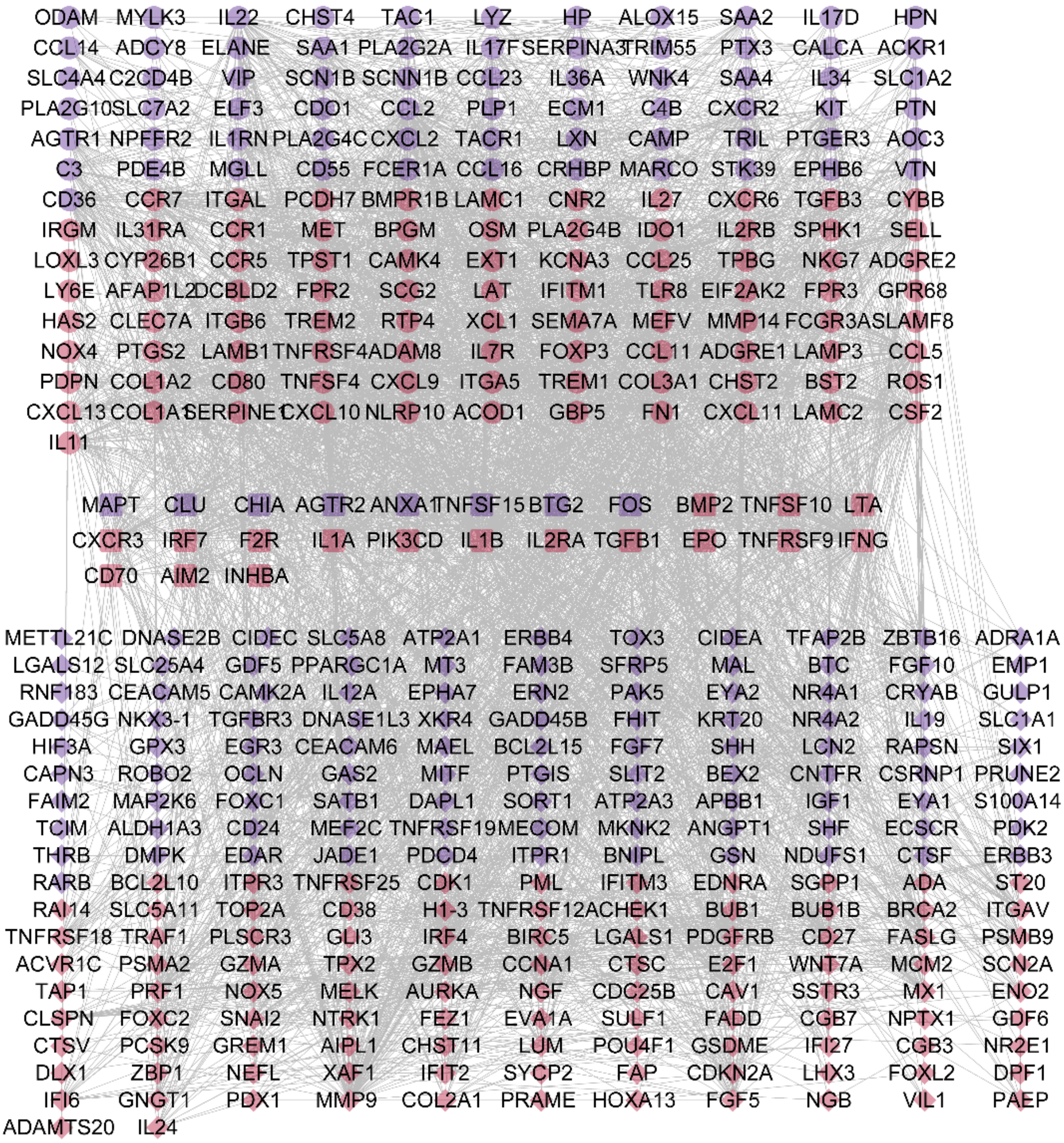


**Figure S9** PPI network between the deIRGs and deAGs in first-stage HNSC . The round nodes located at the upper section of the network represent the IRGs, the diamond-shaped nodes located at the lower section of the network represent the AGs and the rounded-square nodes located at the middle section of the network represent the genes that are IRGs as well as AGs. The red and blue nodes represent the upregulated and downregulated genes, respectively, in first-stage HNSC.

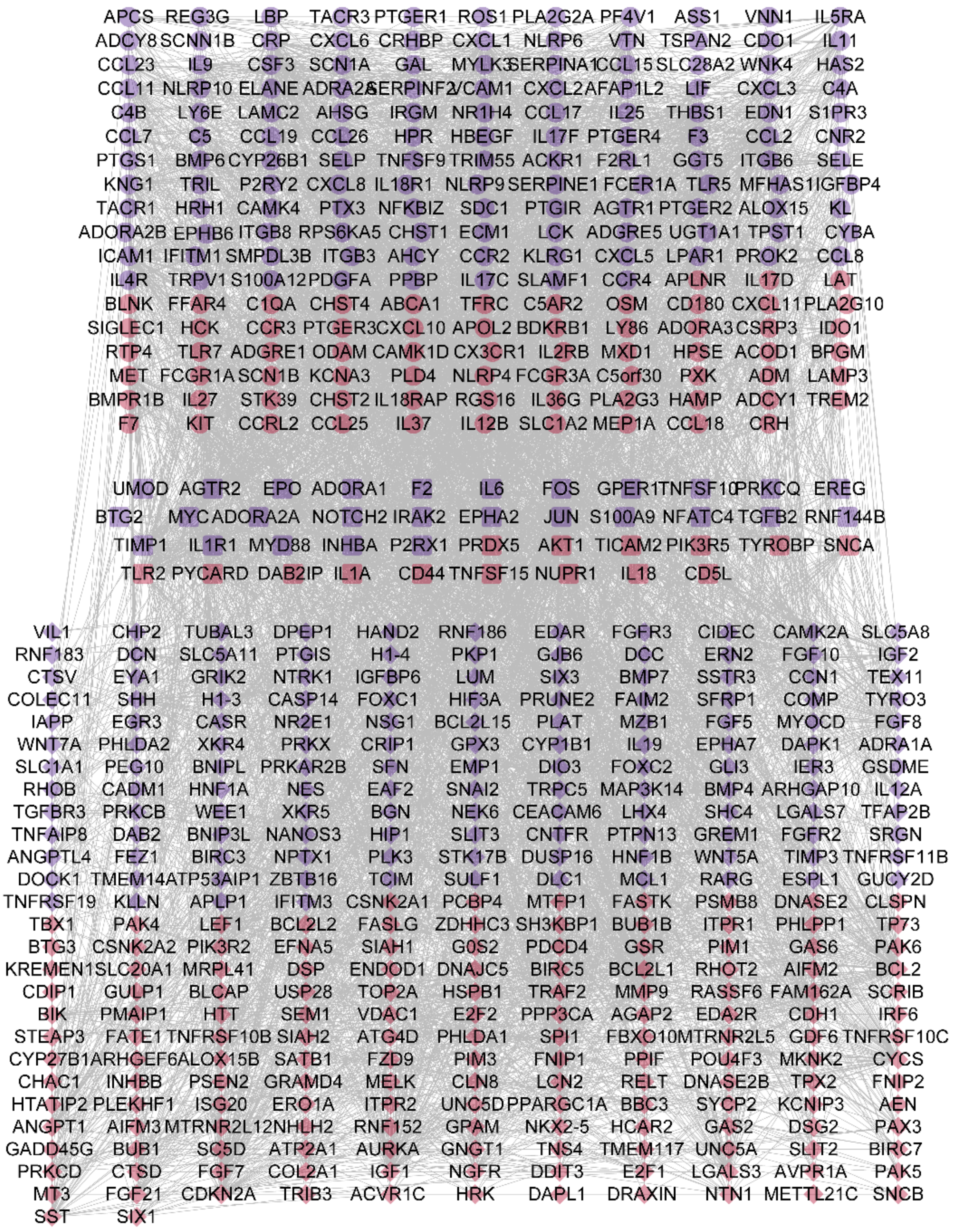


**Figure S10** PPI network between the deIRGs and deAGs in first-stage KICH . The round nodes located at the upper section of the network represent the IRGs, the diamond-shaped nodes located at the lower section of the network represent the AGs and the rounded-square nodes located at the middle section of the network represent the genes that are IRGs as well as AGs. The red and blue nodes represent the upregulated and downregulated genes, respectively, in first-stage KICH.

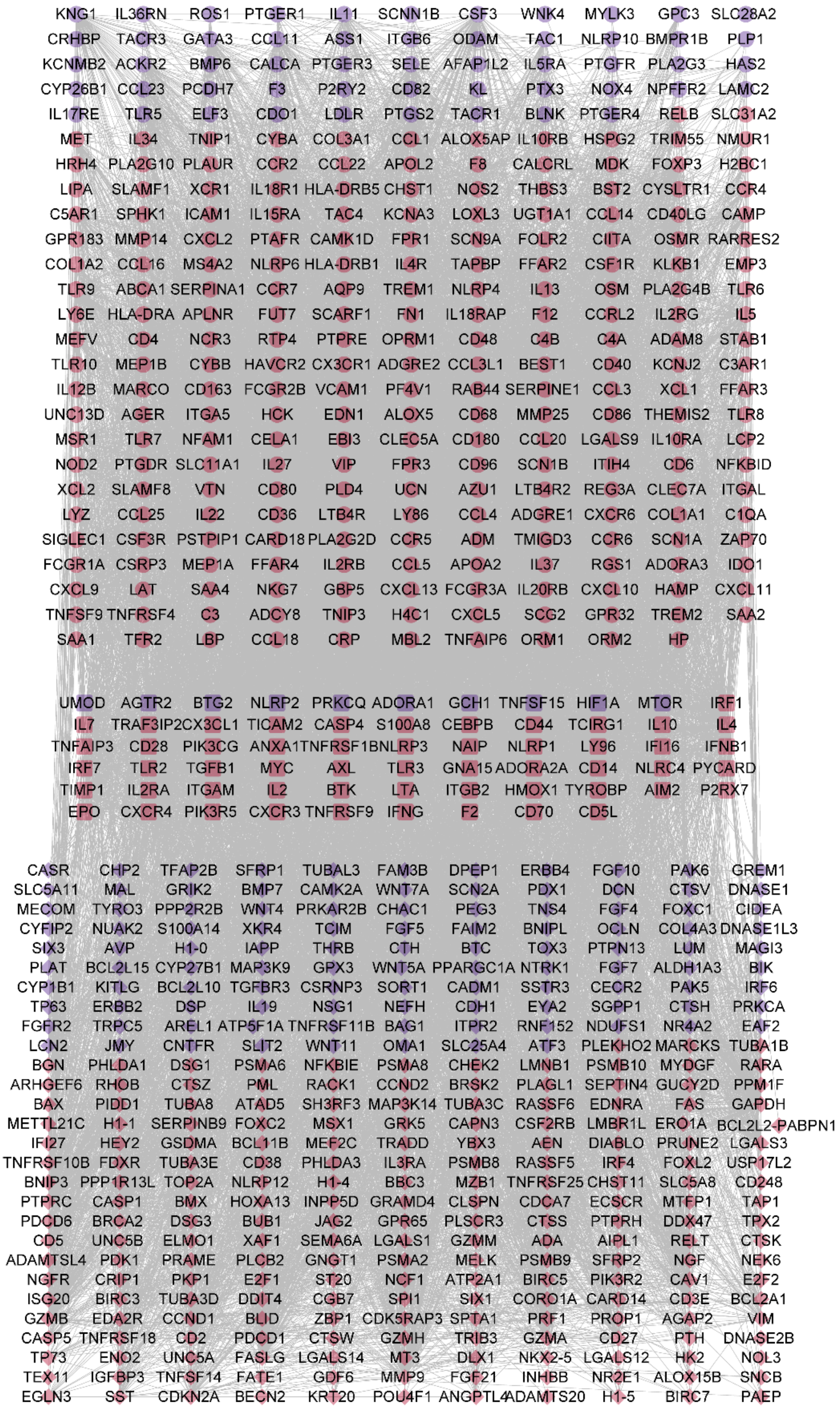

**Figure S11** PPI network between the deIRGs and deAGs in first-stage KIRC . The round nodes located at the upper section of the network represent the IRGs, the diamond-shaped nodes located at the lower section of the network represent the AGs and the rounded-square nodes located at the middle section of the network represent the genes that are IRGs as well as AGs. The red and blue nodes represent the upregulated and downregulated genes, respectively, in first-stage KIRC.

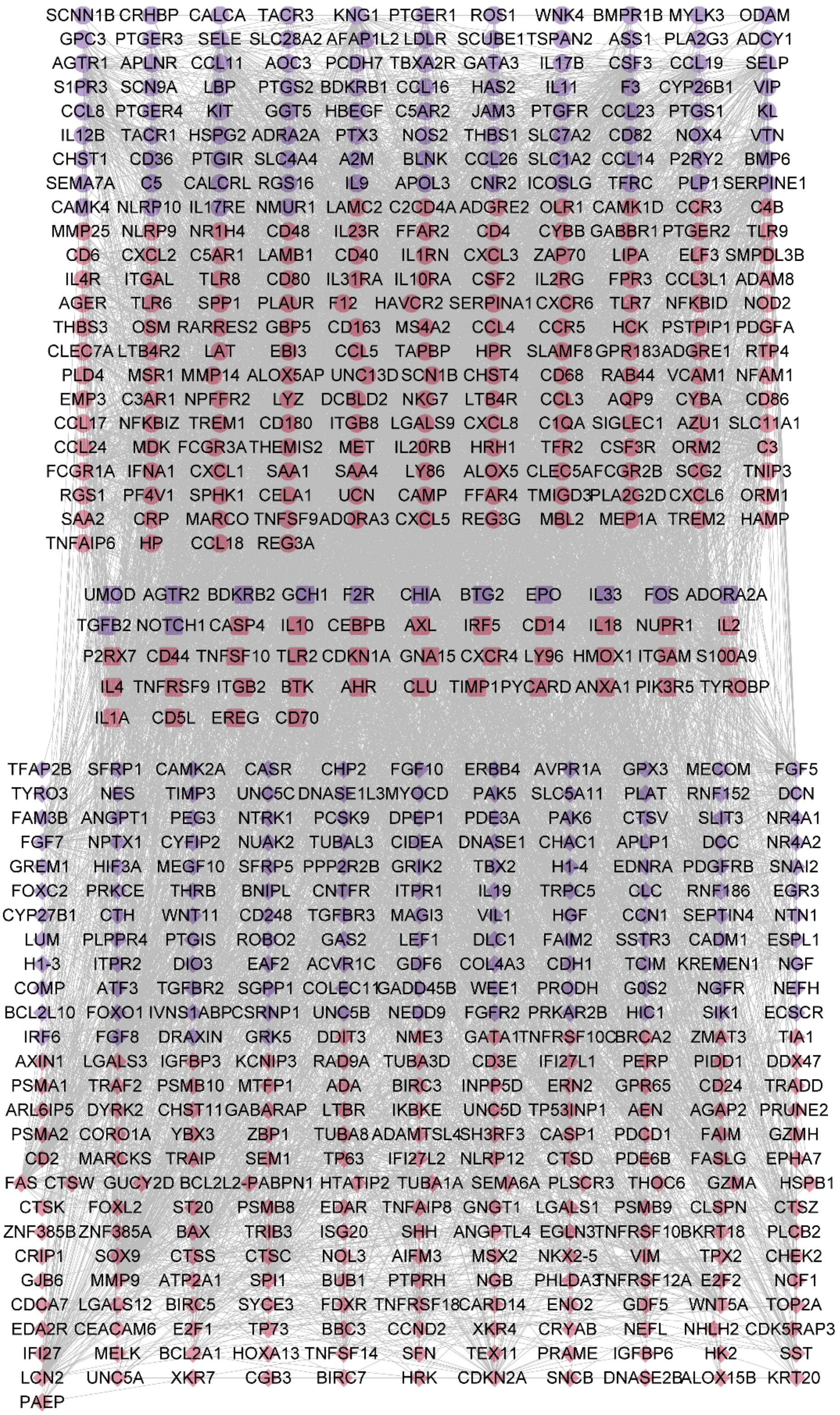

**Figure S12** PPI network between the deIRGs and deAGs in first-stage KIRP . The round nodes located at the upper section of the network represent the IRGs, the diamond-shaped nodes located at the lower section of the network represent the AGs and the rounded-square nodes located at the middle section of the network represent the genes that are IRGs as well as AGs. The red and blue nodes represent the upregulated and downregulated genes, respectively, in first-stage KIRP.

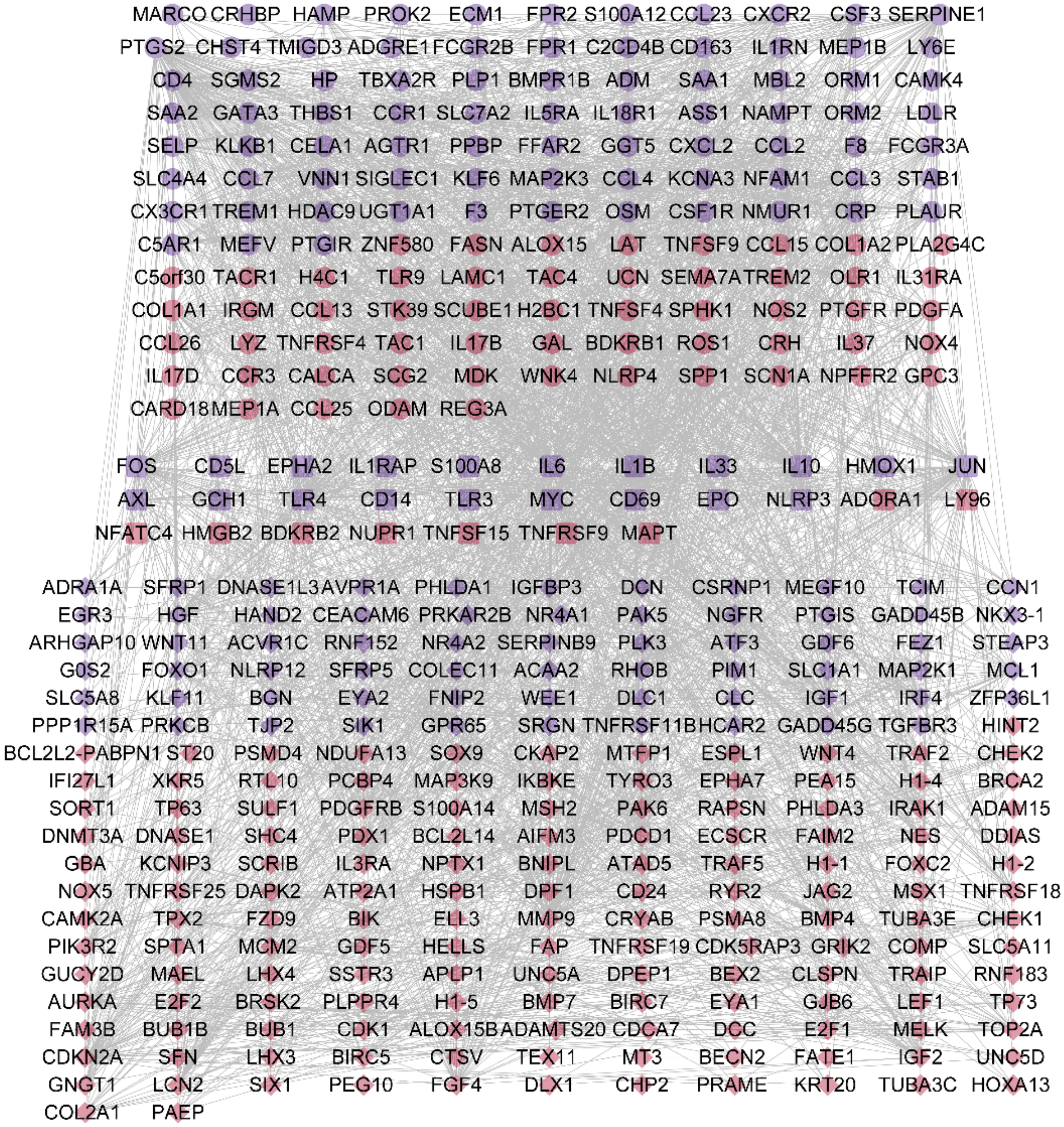


**Figure S13** PPI network between the deIRGs and deAGs in first-stage LIHC . The round nodes located at the upper section of the network represent the IRGs, the diamond-shaped nodes located at the lower section of the network represent the AGs and the rounded-square nodes located at the middle section of the network represent the genes that are IRGs as well as AGs. The red and blue nodes represent the upregulated and downregulated genes, respectively, in first-stage LIHC.

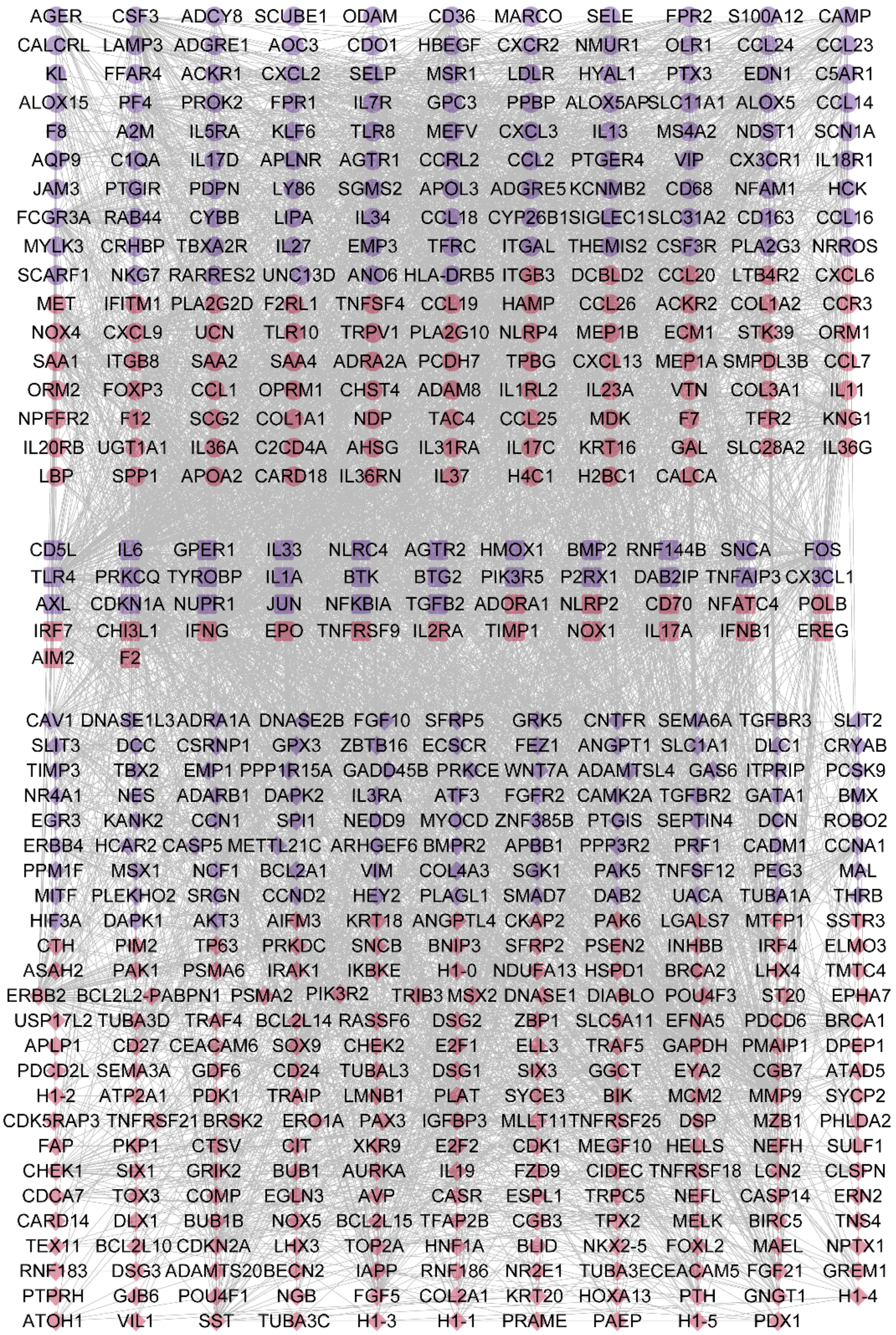


**Figure S14** PPI network between the deIRGs and deAGs in first-stage LUAD . The round nodes located at the upper section of the network represent the IRGs, the diamond-shaped nodes located at the lower section of the network represent the AGs and the rounded-square nodes located at the middle section of the network represent the genes that are IRGs as well as AGs. The red and blue nodes represent the upregulated and downregulated genes, respectively, in first-stage LUAD.

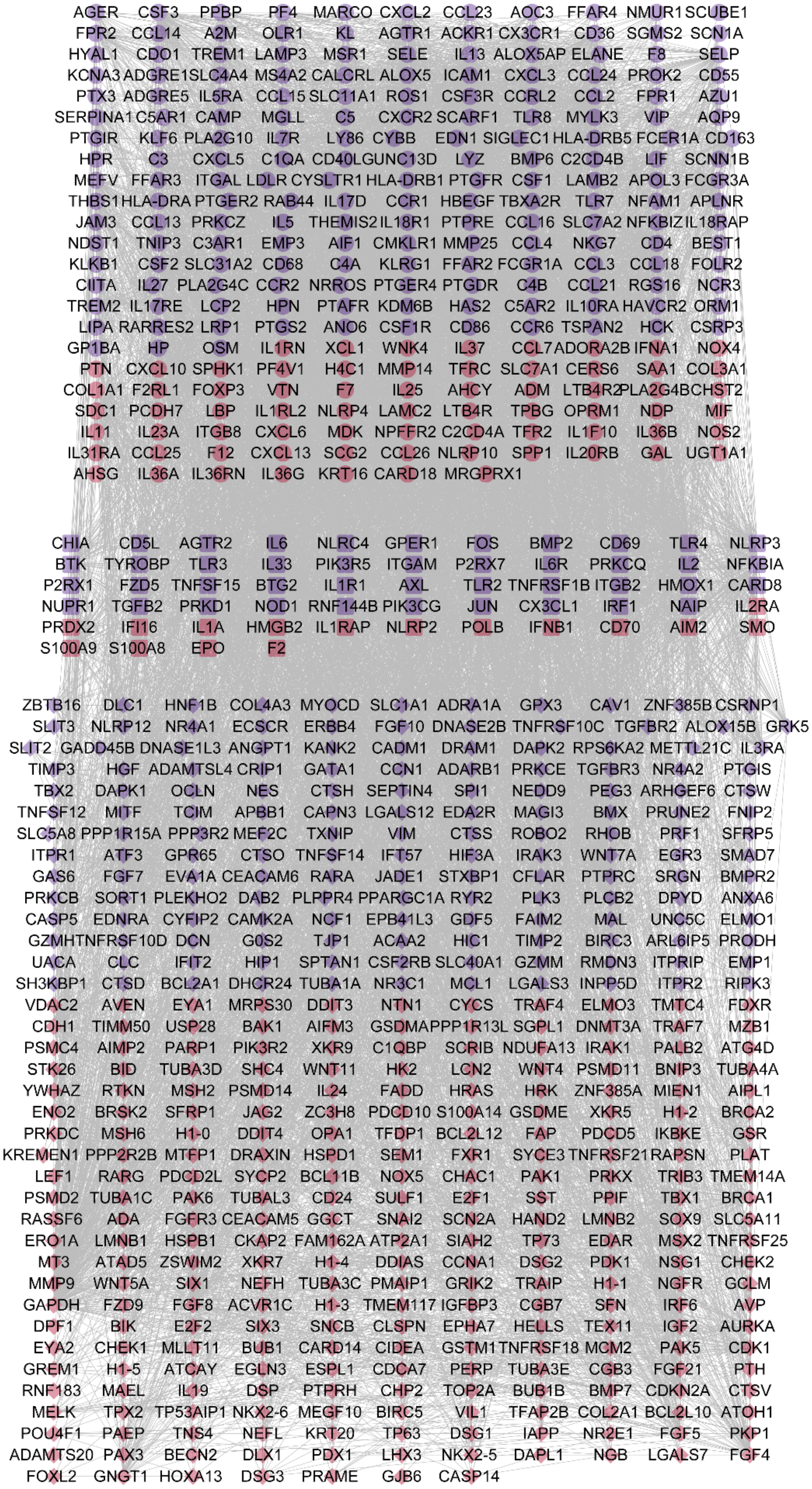


**Figure S15** PPI network between the deIRGs and deAGs in first-stage LUSC . The round nodes located at the upper section of the network represent the IRGs, the diamond-shaped nodes located at the lower section of the network represent the AGs and the rounded-square nodes located at the middle section of the network represent the genes that are IRGs as well as AGs. The red and blue nodes represent the upregulated and downregulated genes, respectively, in first-stage LUSC.

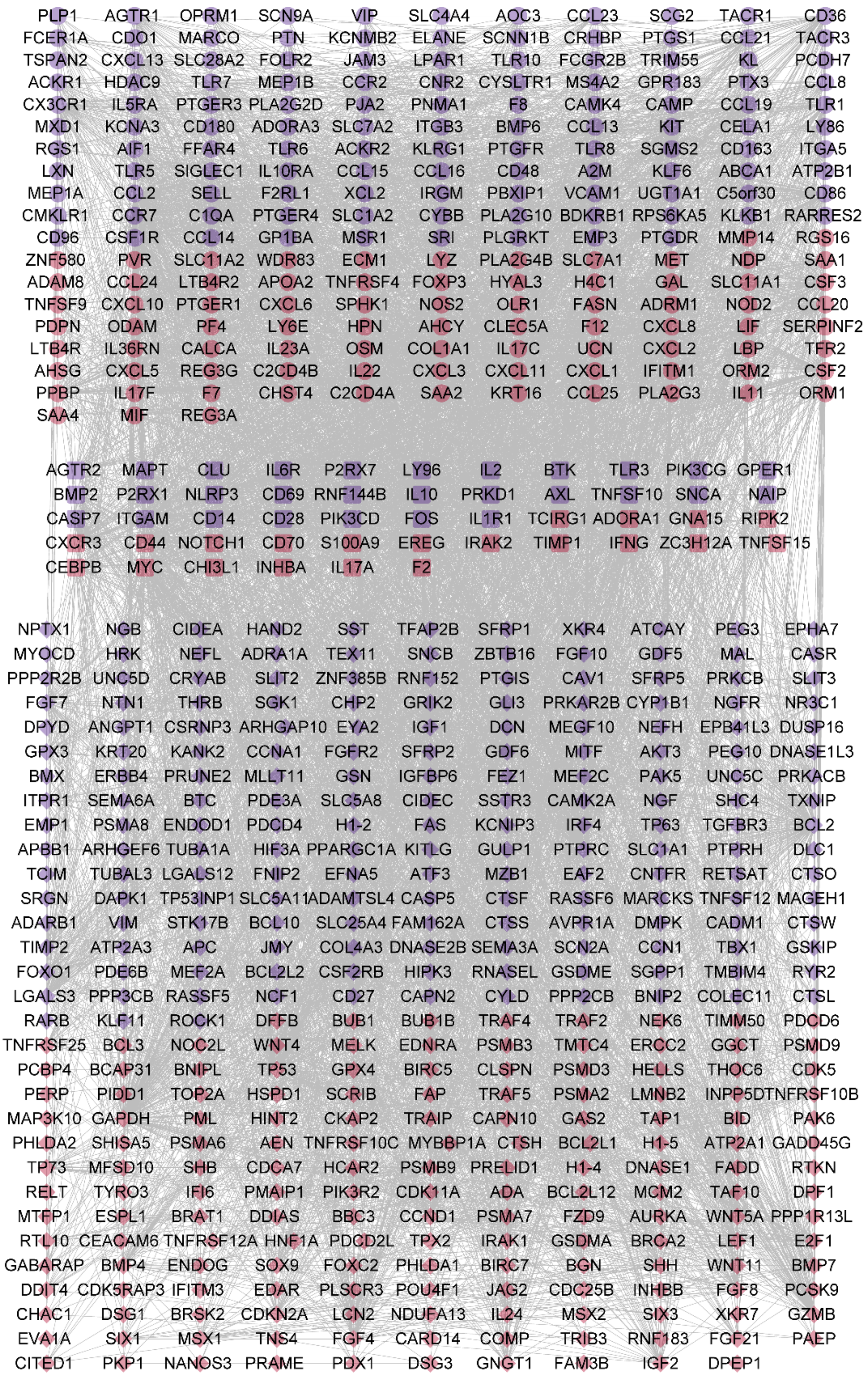


**Figure S16** PPI network between the deIRGs and deAGs in first-stage READ . The round nodes located at the upper section of the network represent the IRGs, the diamond-shaped nodes located at the lower section of the network represent the AGs and the rounded-square nodes located at the middle section of the network represent the genes that are IRGs as well as AGs. The red and blue nodes represent the upregulated and downregulated genes, respectively, in first-stage READ.

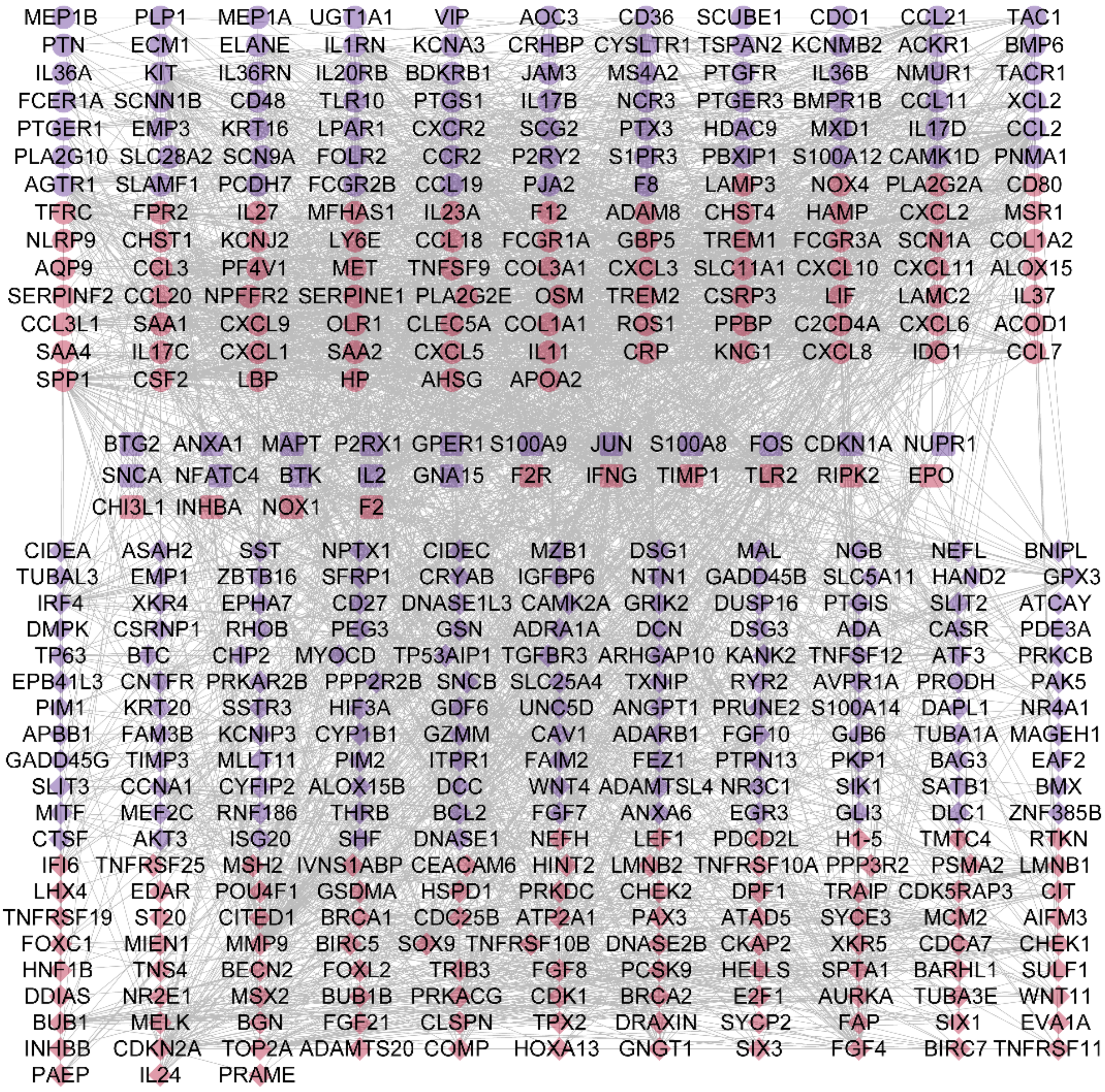


**Figure S17** PPI network between the deIRGs and deAGs in first-stage STAD . The round nodes located at the upper section of the network represent the IRGs, the diamond-shaped nodes located at the lower section of the network represent the AGs and the rounded-square nodes located at the middle section of the network represent the genes that are IRGs as well as AGs. The red and blue nodes represent the upregulated and downregulated genes, respectively, in first-stage STAD.

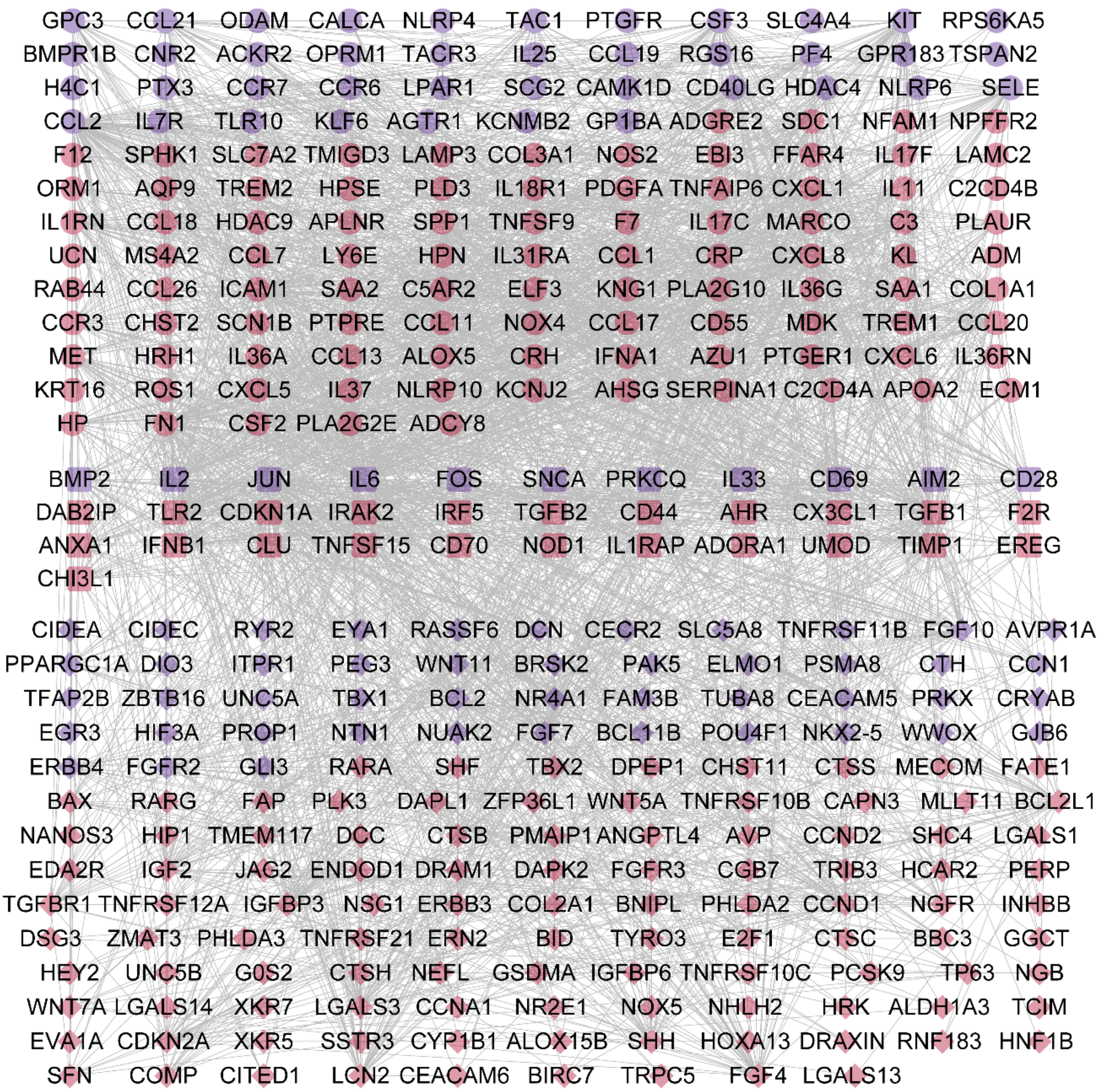


**Figure S18** PPI network between the deIRGs and deAGs in first-stage THCA . The round nodes located at the upper section of the network represent the IRGs, the diamond-shaped nodes located at the lower section of the network represent the AGs and the rounded-square nodes located at the middle section of the network represent the genes that are IRGs as well as AGs. The red and blue nodes represent the upregulated and downregulated genes, respectively, in first-stage THCA.

**Table S8** Number of DEMs obtained first-stage cancer and respective normal tissue samples

| Cancer type | No. of first-stage cancer samples | No. of normal cases | No. of DEMs | No. of upregulated DEMs | No. of downregulated DEMs |
|---|---|---|---|---|---|
| BRCA | 184 | 104 | 334 | 209 | 125 |
| CHOL | 19 | 9 | 141 | 90 | 51 |
| COAD | 75 | 8 | 508 | 215 | 293 |
| ESCA | 18 | 13 | 69 | 42 | 27 |
| HNSC | 27 | 44 | 275 | 141 | 134 |
| KICH | 21 | 25 | 320 | 143 | 177 |
| KIRC | 274 | 71 | 256 | 126 | 130 |
| KIRP | 173 | 34 | 353 | 155 | 198 |
| LIHC | 172 | 50 | 342 | 301 | 41 |
| LUAD | 281 | 46 | 382 | 221 | 161 |
| LUSC | 230 | 45 | 515 | 417 | 98 |
| READ | 29 | 3 | 260 | 164 | 96 |
| STAD | 58 | 45 | 222 | 174 | 48 |
| THCA | 284 | 59 | 166 | 123 | 43 |

The visualization of the differential miRNA expression analysis between the first-stage cancers and their respective normal tissue samples have been done using volcano plots. The statistically significant DEMs obtained for each cancer have been represented by red dots in the volcano plots. The gene names of top 5 upregulated and downregulated genes in each of the first-stage cancers have been represented in their volcano plots (**Error! Reference source not found.**).

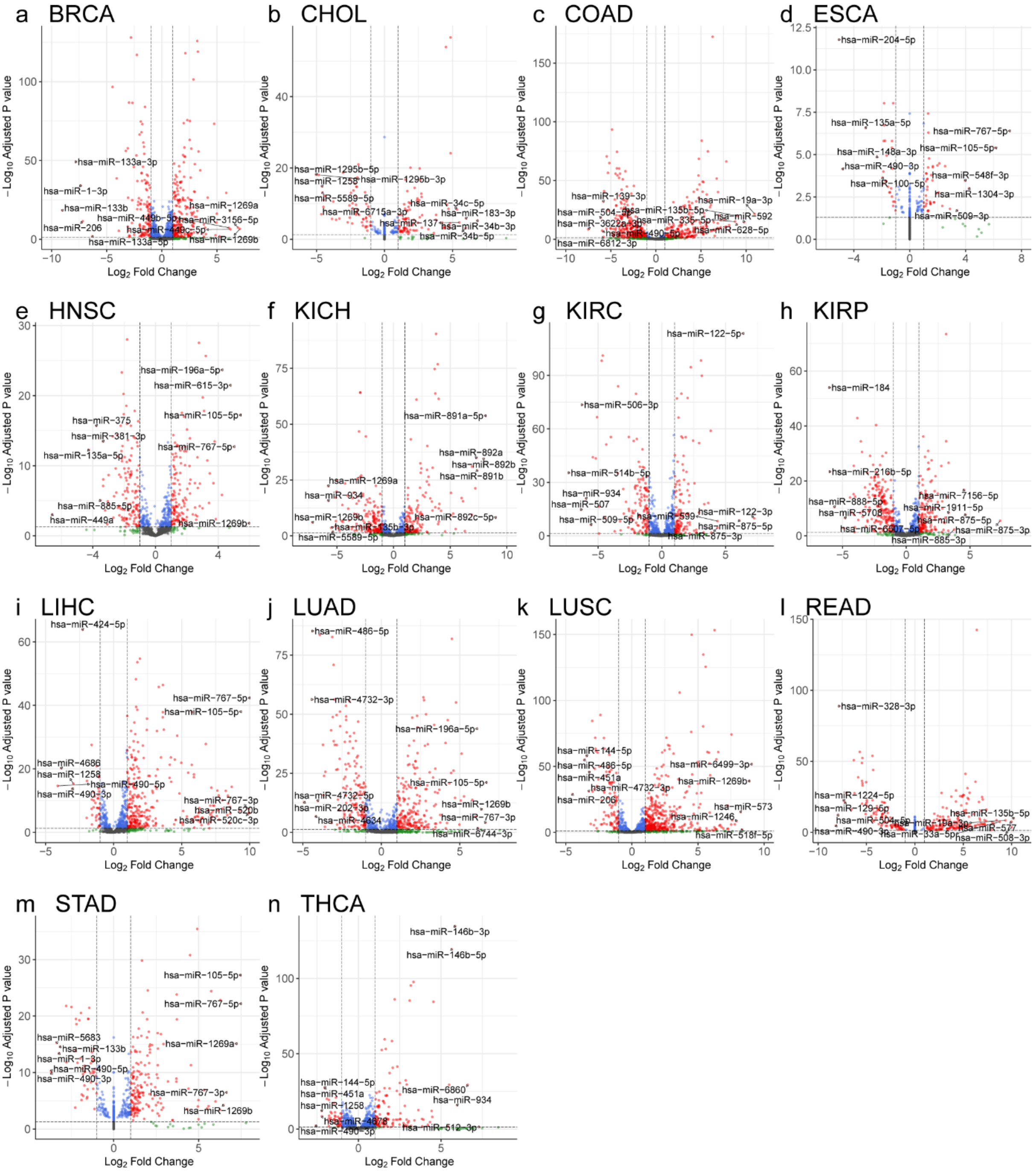


**Figure S19** Volcano plots for the differential miRNA expression analysis results of the 14 cancers. (a) BRCA (b) CHOL (c) COAD (d) ESCA (e) HNSC (f) KICH (g) KIRC (h) KIRP (i) LIHC (j) LUAD (k) LUSC (l) READ (m) STAD (n) THCA. The red dots indicate the statistically significant DEMs (|log2FC| > 1 and padj < 0.05) between normal and first-stage cancer. The green dots indicate miRNAs with padj >=0.05. The blue dots indicate the miRNAs with |log2FC| <= 1. The miRNAs dots represent the genes with |log2FC| < 1 and padj >= 0.05.

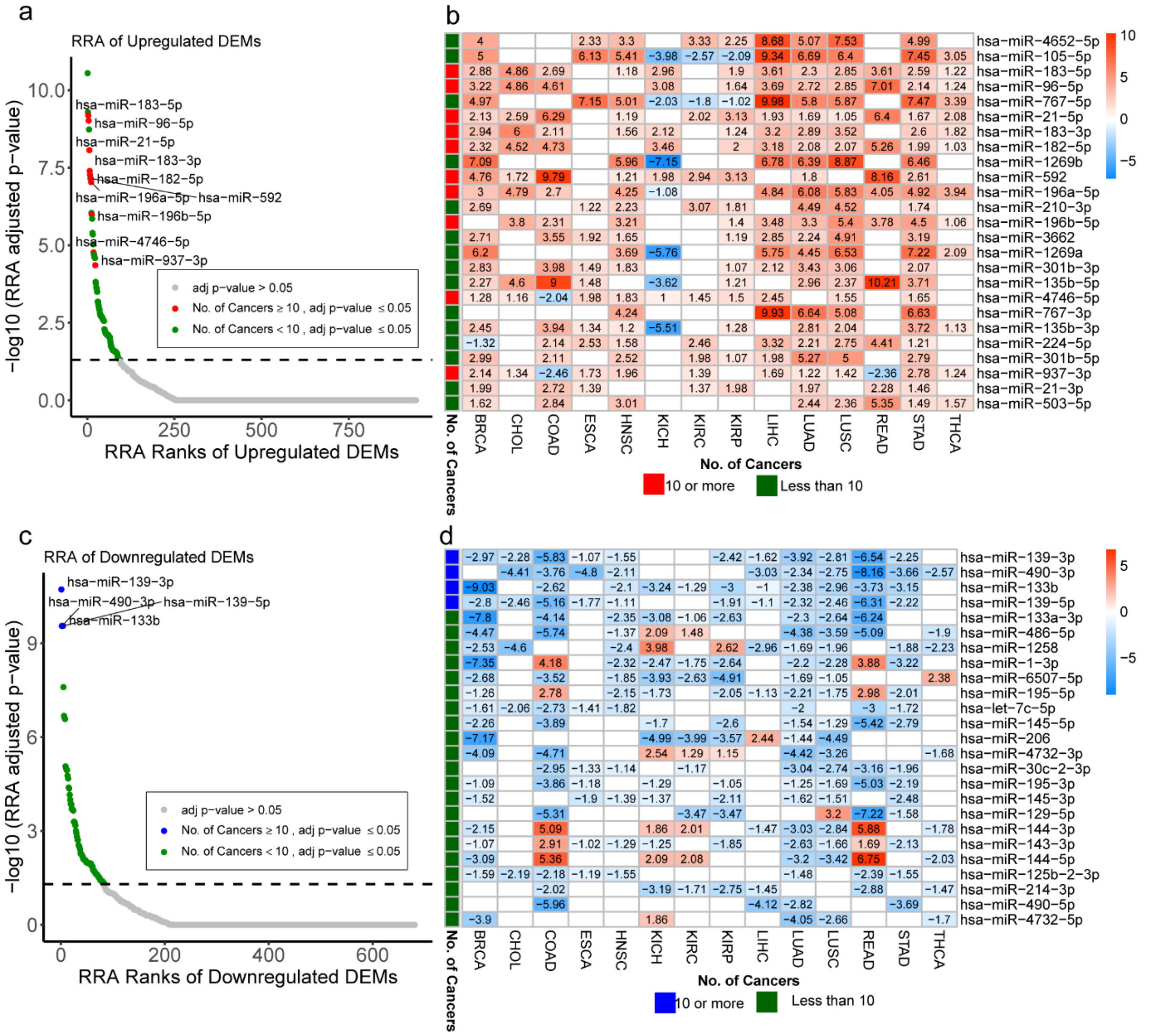


**Figure S20** RRA of Upregulated and Downregulated DEMs of 14 Cancers: (a) The dotted line represents the padj cutoff value of 0.05. The red dots represent the statistically significant RRA ranked miRNAs that are upregulated in atleast 10 cancers. The green dots represent the statistically significant RRA ranked miRNAs that are upregulated in less than 10 cancers. The grey dots represent the statistically insignificant RRA ranked miRNAs (padj>0.05). (b) Heatmap representing the log2FC of the top 25 upregulated DEMs in the 14 cancers. (c) The dotted line represents the padj cutoff value of 0.05. The blue dots represent the statistically significant RRA ranked miRNAs that are downregulated in atleast 10 cancers. The green dots represent the statistically significant RRA ranked miRNAs that are downregulated in less than 10 cancers. The grey dots represent the statistically insignificant RRA ranked miRNAs (padj>0.05). (d) Heatmap representing the log2FC of the top 25 downregulated DEMs in the 14 cancers.

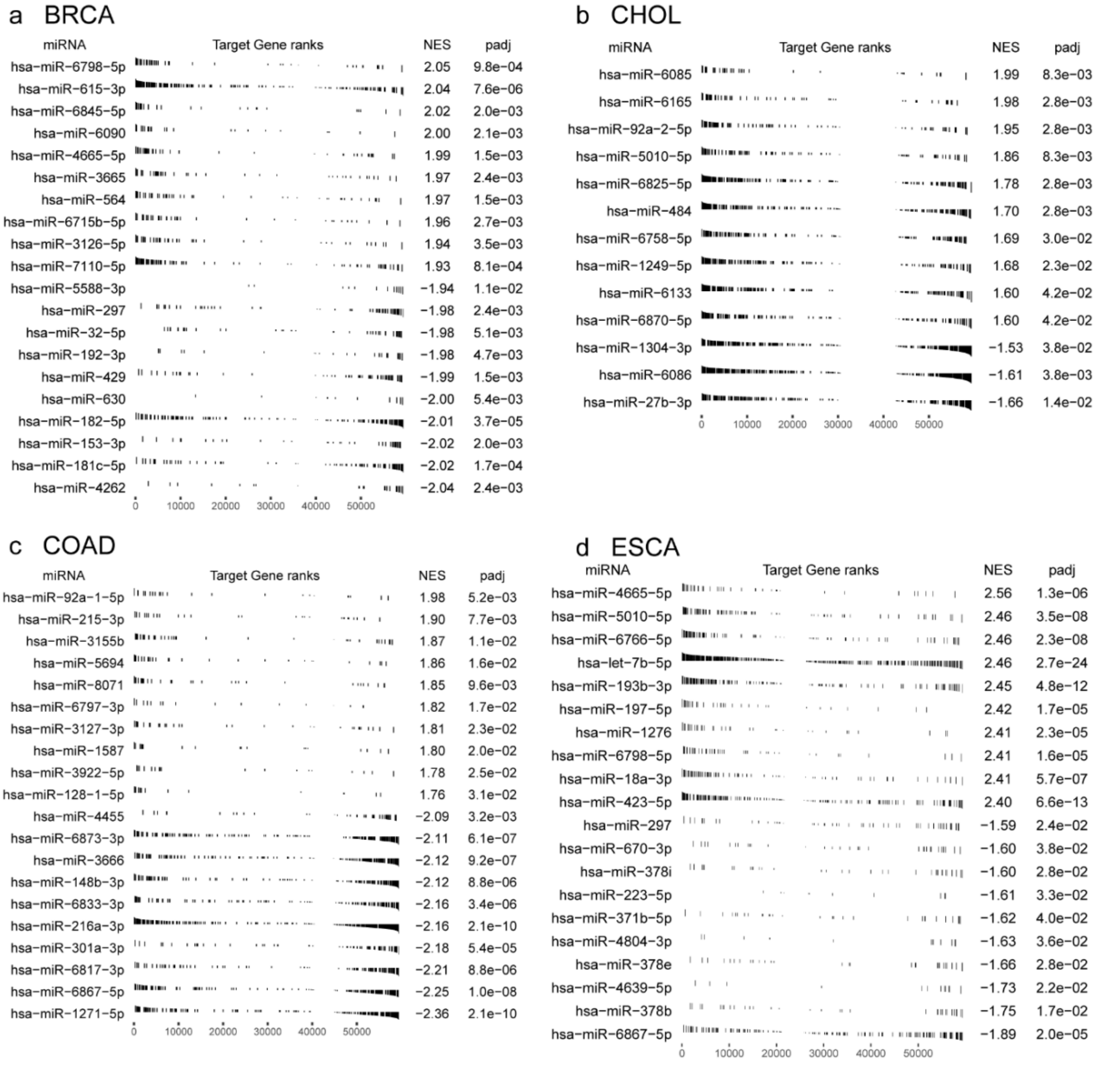


**Figure S21** GSEA tables representing the top miRNAs with dysregulated gene targets for 14 cancers. (a) BRCA (b) CHOL (c) COAD (d) ESCA (e) HNSC (f) KICH (g) KIRC (h) KIRP (i) LIHC (j) LUAD (k) LUSC (l) READ (m) STAD (n) THCA.
(Detailed results in Supplementary Table S4_4)

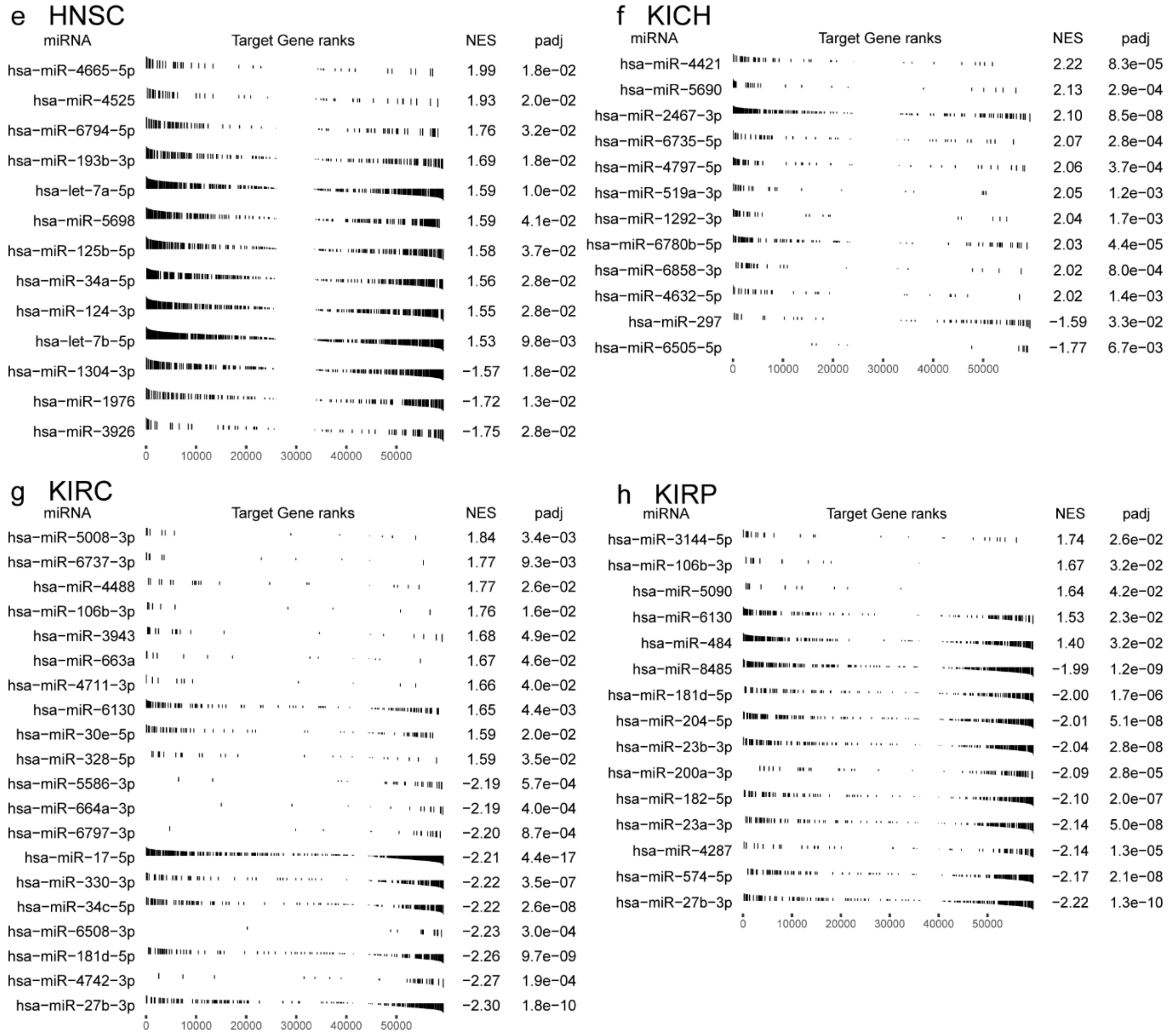


**Figure S21** (continued) GSEA tables representing the top miRNAs with dysregulated gene targets for 14 cancers. (a) BRCA (b) CHOL (c) COAD (d) ESCA (e) HNSC (f) KICH (g) KIRC (h) KIRP (i) LIHC (j) LUAD (k) LUSC (l) READ (m) STAD (n) THCA.
(Detailed results in Supplementary Table S4_4)

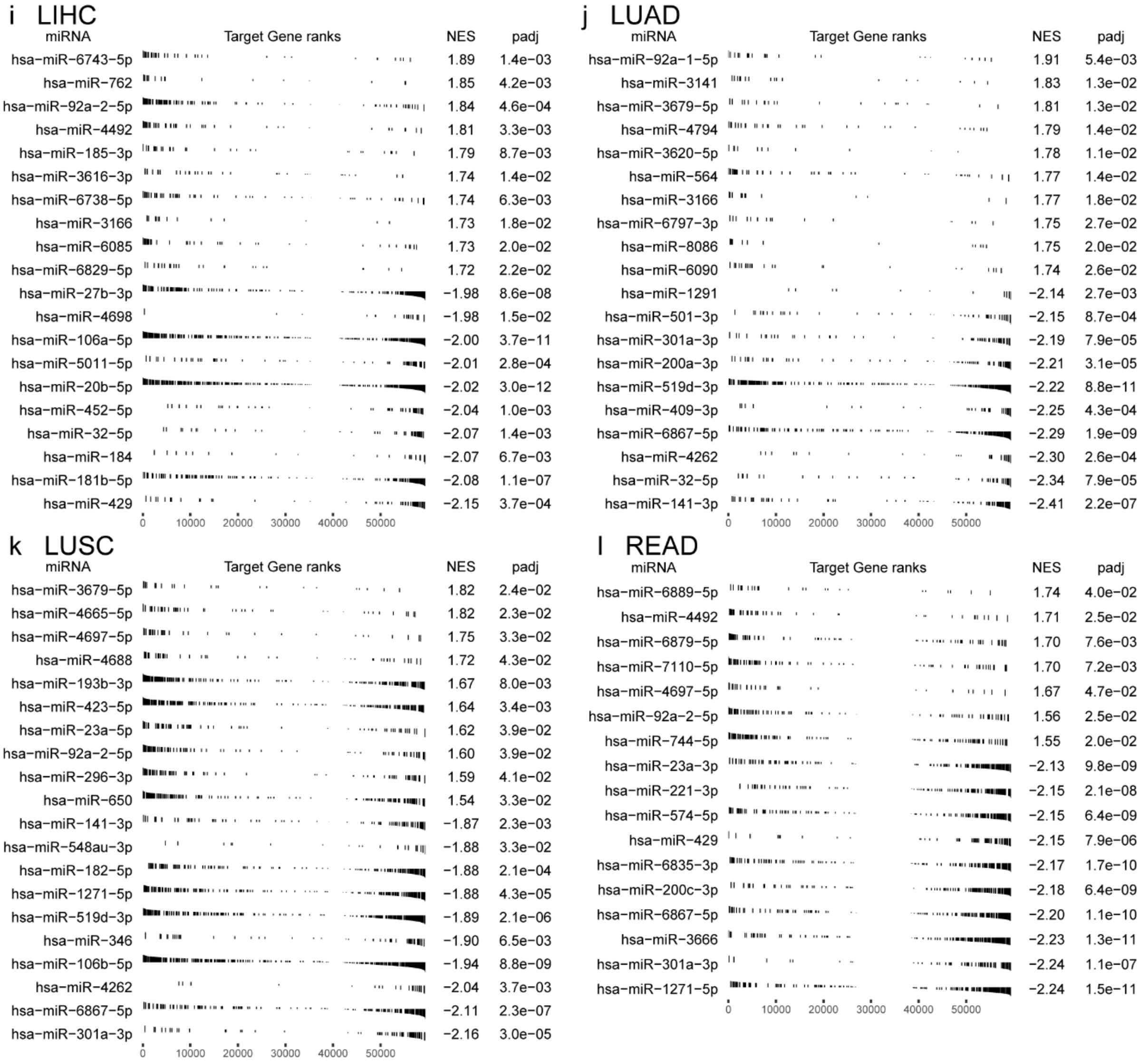


**Figure S21** (continued) GSEA tables representing the top miRNAs with dysregulated gene targets for 14 cancers. (a) BRCA (b) CHOL (c) COAD (d) ESCA (e) HNSC (f) KICH (g) KIRC (h) KIRP (i) LIHC (j) LUAD (k) LUSC (l) READ (m) STAD (n) THCA.
(Detailed results in Supplementary Table S4_4)

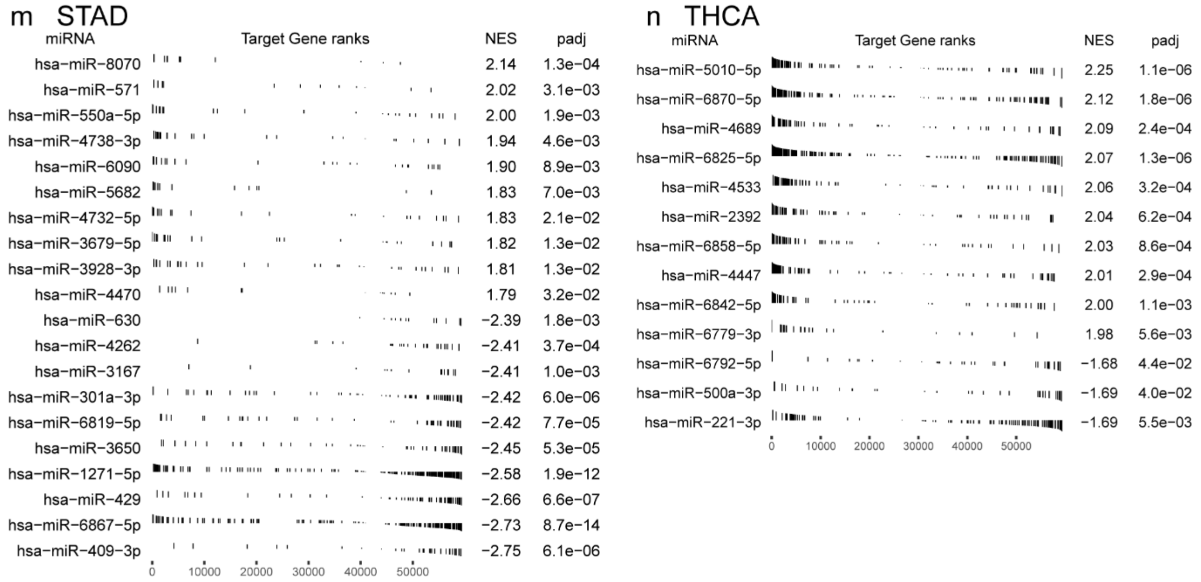


**Figure S21** (continued) GSEA tables representing the top miRNAs with dysregulated gene targets for 14 cancers. (a) BRCA (b) CHOL (c) COAD (d) ESCA (e) HNSC (f) KICH (g) KIRC (h) KIRP (i) LIHC (j) LUAD (k) LUSC (l) READ (m) STAD (n) THCA.
(Detailed results in Supplementary Table S4_4)

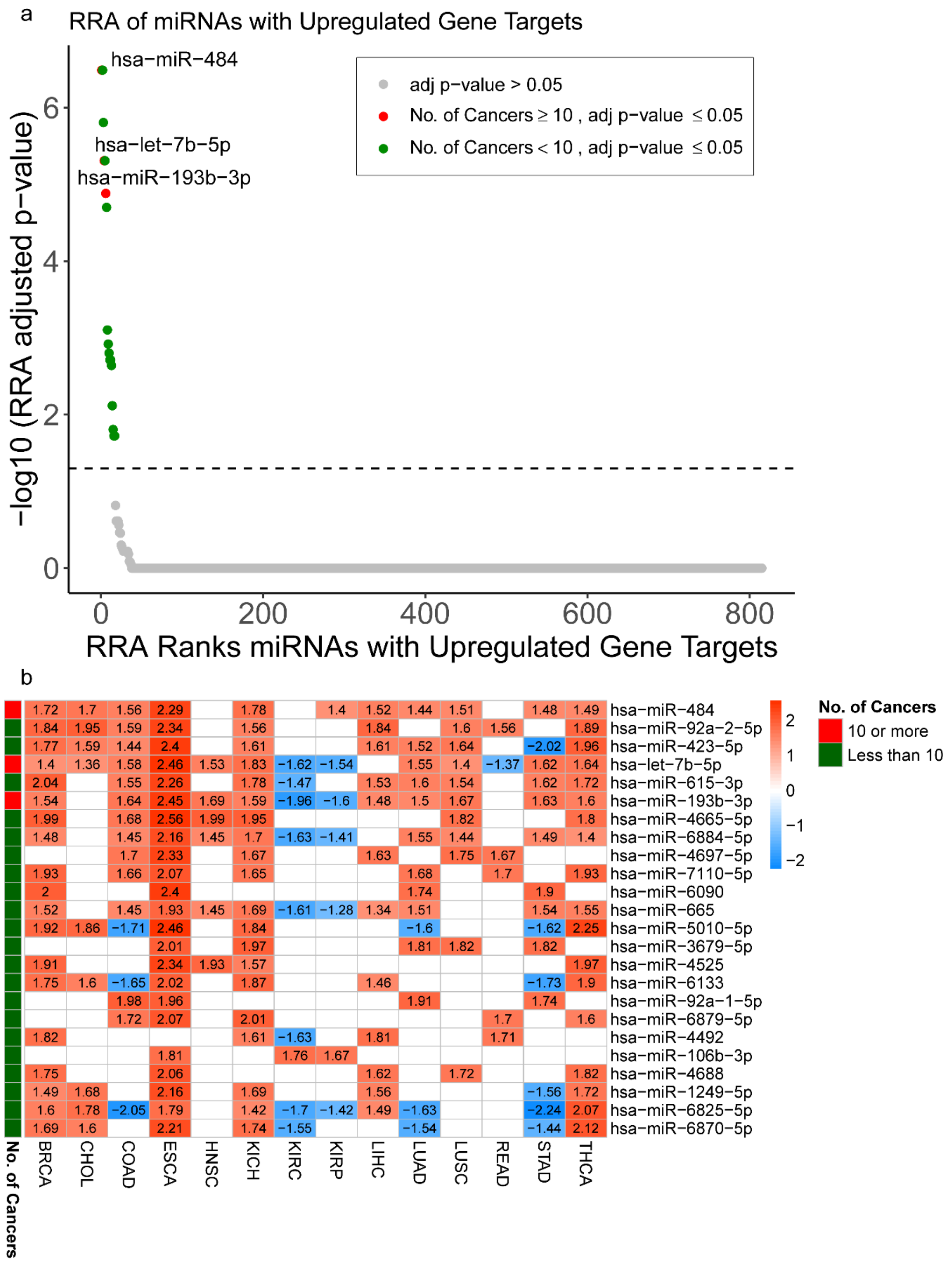


**Figure S22** RRA of miRNAs with Upregulated Gene Targets in 14 Cancers: (a) The dotted line represents the padj cutoff value of 0.05. The red dots represent the statistically significant RRA ranked miRNAs with upregulated gene targets in atleast 10 cancers. The green dots represent the statistically significant RRA ranked miRNAs with upregulated gene targets in less than 10 cancers. The grey dots represent the statistically insignificant RRA ranked miRNAs with upregulated gene targets (padj>0.05). (b) Heatmap representing the NES of the statistically significant RRA-ranked miRNAs with upregulated gene targets in the 14 cancers. (Details of RRA Analysis provided in Supplementary table S4_5)

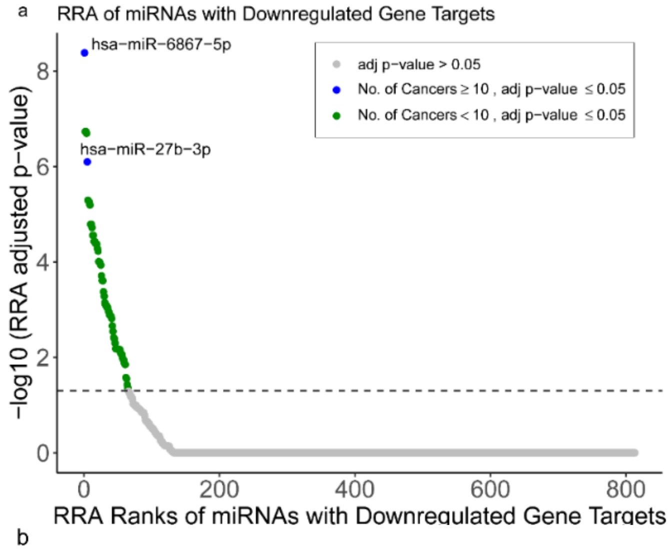


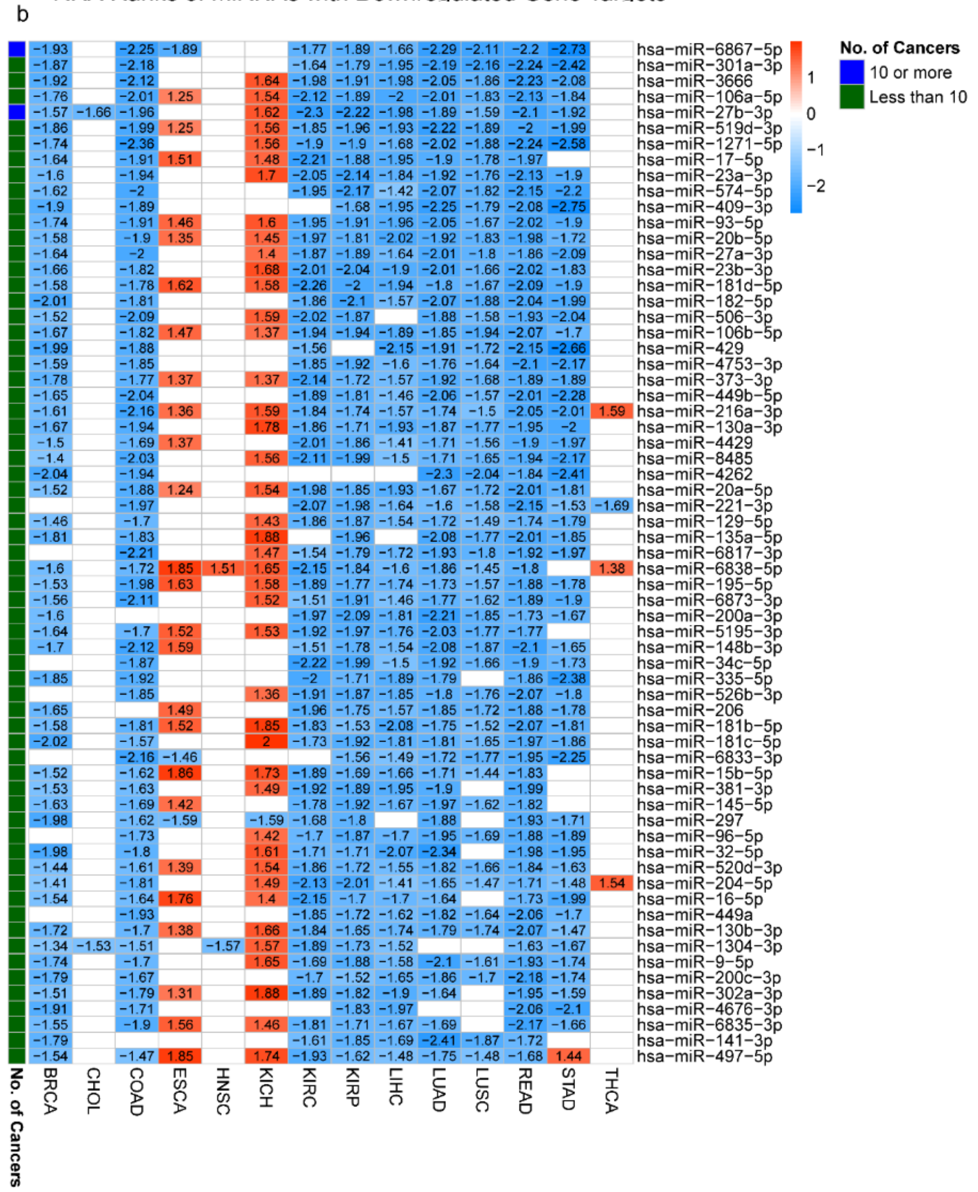


**Figure S23** RRA of miRNAs with Downregulated Gene Targets in 14 Cancers.

(a) The dotted line represents the padj cutoff value of 0.05. The blue dots represent the statistically significant RRA ranked miRNAs with downregulated gene targets in atleast 10 cancers. The green dots represent the statistically significant RRA ranked miRNAs with downregulated gene targets in less than 10 cancers. The grey dots represent the statistically insignificant RRA ranked miRNAs with downregulated gene targets (padj>0.05).

(b) Heatmap representing the NES of the statistically significant RRA-ranked miRNAs with downregulated gene targets in the 14 cancers.

Details of RRA Analysis provided in Supplementary table S4_6)

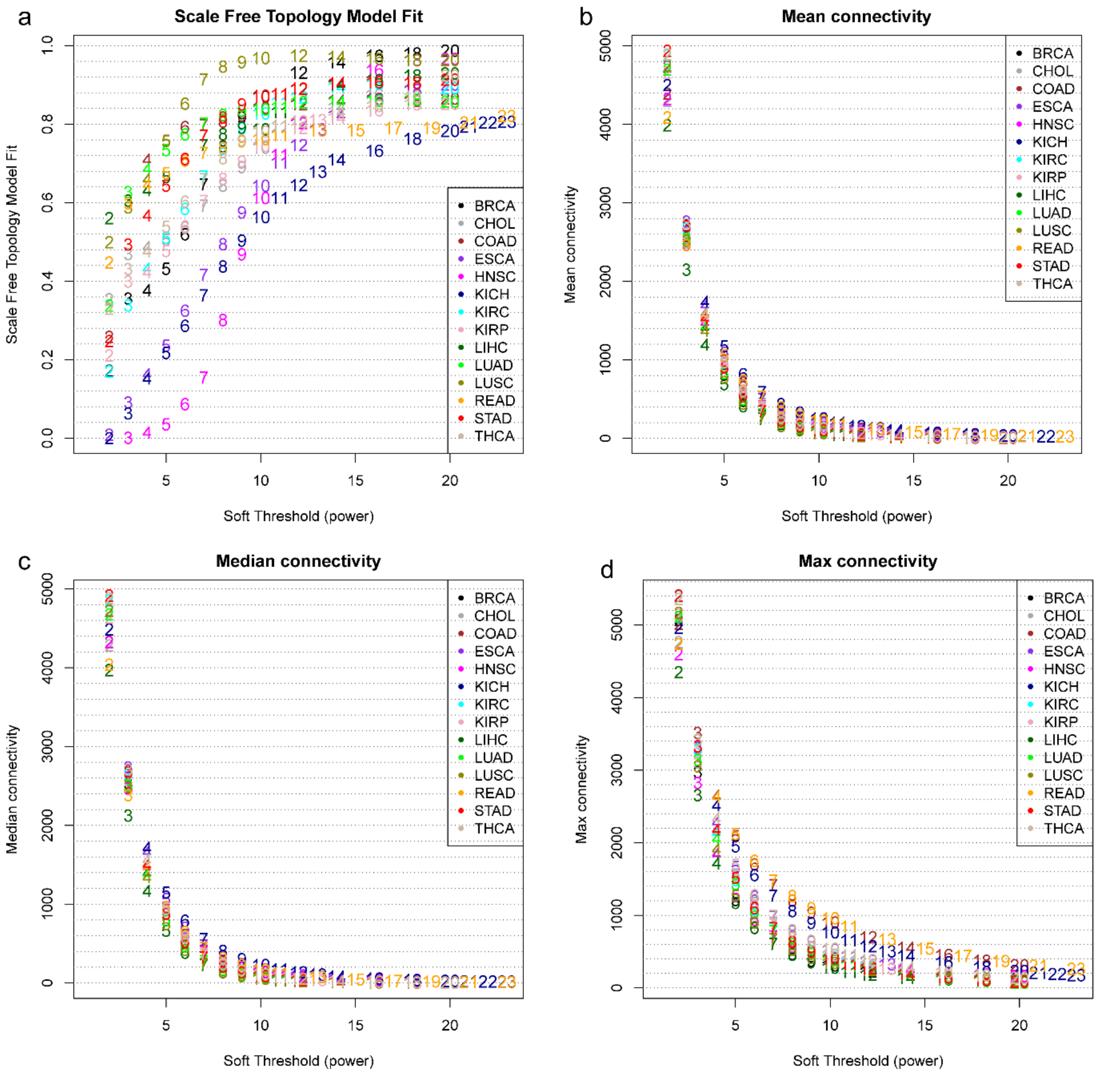


**Figure S24** Parameters for Individual cancer WGCNA of 14 cancers: (a) The soft thresholding power (β) selection was done keeping the scale-free topology fit index (R^2) of atleast 0.8. (b) The mean connectivity against the values of soft thresholding power for each type of cancer. (c) The median connectivity against the values of soft thresholding power for each type of cancer. (d) The mode connectivity against the values of soft thresholding power for each type of cancer.

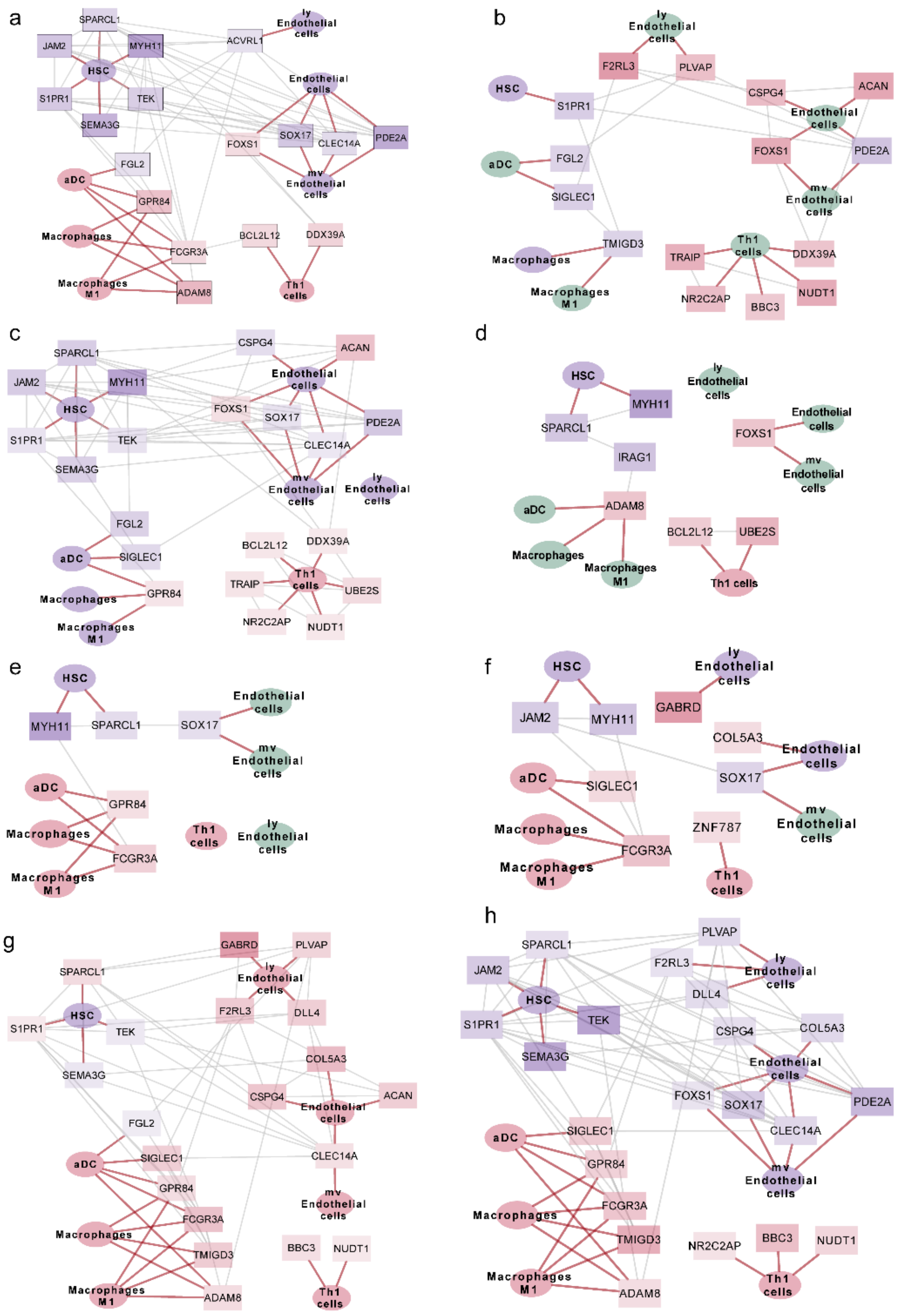


**Figure S25** The cancer-specific network models obtained by mapping the details of DEGs and PCI cells in the pan-cancer network model. (a) BRCA (b) CHOL (c) COAD (d) ESCA (e) HNSC (f) KICH (g) KIRC (h) KIRP (i) LIHC (j) LUAD (k) LUSC (l) READ (m) STAD (n) THCA. The red and blue colour gradients of rectangular gene nodes represent the upregulation and downregulation of the gene in that specific cancer, respectively. The oval nodes represent the PCI cells with red and blue colour signifying their increased and decreased infiltration the cancer tissue, respectively, compared to the respective normal tissue.

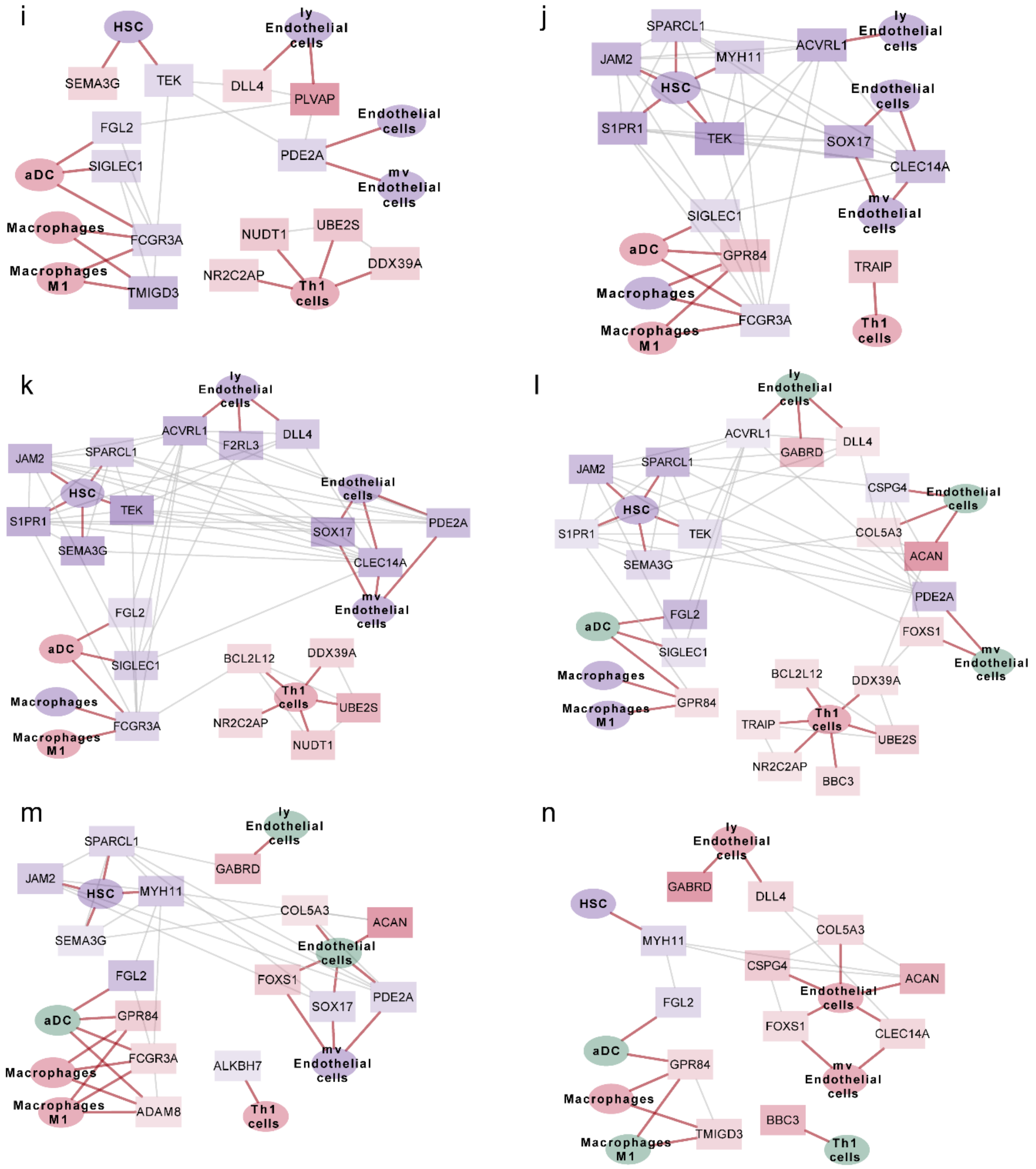


**Figure S25** (continued). The cancer-specific network models obtained by mapping the details of DEGs and PCI cells in the pan-cancer network model. (a) BRCA (b) CHOL (c) COAD (d) ESCA (e) HNSC (f) KICH (g) KIRC (h) KIRP (i) LIHC (j) LUAD (k) LUSC (l) READ (m) STAD (n) THCA. The red and blue colour gradients of rectangular gene nodes represent the upregulation and downregulation of the gene in that specific cancer, respectively. The oval nodes represent the PCI cells with red and blue colour signifying their increased and decreased infiltration the cancer tissue, respectively, compared to the respective normal tissue.
